# Seeing the unseen: A trifoliate (MYB117) mutant allele fortifies folate and carotenoids in tomato fruits

**DOI:** 10.1101/2021.09.28.462071

**Authors:** Kamal Tyagi, Anusha Sunkum, Meenakshi Rai, Supriya Sarma, Nidhi Thakur, Amita Yadav, Sanchari Sircar, Yellamaraju Sreelakshmi, Rameshwar Sharma

## Abstract

Micronutrient deficiency also termed hidden hunger affects a large segment of the human population, particularly in developing and underdeveloped nations. Tomato the second most consumed vegetable crop in the world after potato can serve as a sustainable source to alleviate micronutrient deficiency. In tomato, the mutations in the R2R3-MYB117 transcription factor elicit *trifoliate* leaves and initiate axillary meristems; however, its effect on fruit metabolome remains unexplored. The fruits of a new *trifoliate* (*tf*) allele (*tf-5*) were firmer, had higher °Brix, folate, and carotenoids. The transcriptome, proteome, and metabolome profiling of *tf-5* reflected a broad-spectrum change in homeostasis. The *tf-5* allele enhanced the fruit firmness by suppressing cell wall softening-related proteins. The *tf-5* fruit displayed a substantial increase in aminome, particularly γ-aminobutyric acid, with a parallel reduction in aminoacyl t-RNA synthases. The increased lipoxygenases proteins and transcripts seemingly elevated jasmonic acid. In addition, increased abscisic acid hydrolases transcripts coupled with reduced precursor supply lowered abscisic acid. The upregulation of carotenoids was mediated by modulation of methylerythreitol and plastoquinone pathways along with an increase in carotenoids isomerization proteins. The upregulation of folate in *tf*-5 was connoted by the increase in precursor *p*-aminobenzoic acid and transcripts of several folate biosynthesis pathway genes. The reduction in pterin-6-carboxylate and γ-glutamyl hydrolase activity indicated that the diminished folate degradation also enriched folate levels. Our study delineates that introgression of the *tf-5* can be used for the γ-aminobutyric acid, carotenoids, and folate fortification of tomato.

**One-sentence summary:** A tomato *trifoliate* allele encoding a truncated MYB117 transcription factor alters cellular homeostasis and fortifies γ-aminobutyric acid, folate, and carotenoids in tomato fruits.

## Introduction

Plant-based foods offer a wide range of nutraceuticals that are essential for human health. However, the staple food crops are often poor in some of the nutraceuticals. Thus, commonly available functional foods are of considerable importance to bridge the gap in the supply of essential nutraceuticals. In this respect, tomato (*Solanum lycopersicum*), a globally consumed fruit, is considered an ideal source for health-promoting compounds (Chaudhary et al., 2018). The improvement in nutraceutical levels of tomato has been an important target in classical breeding as well as by biotechnological interventions.

Since the tomato fruits are prominently red-colored, imparted by lycopene, the earliest efforts on fortification were directed towards colored carotenoids. The genetic analysis of fruit coloration mutants identified key loci regulating lycopene and β-carotene accumulation in tomato. The characterization of carotenoid biosynthesis mutants such as *r*, *delta*, *tangerine*, *beta,* and *beta old gold,* uncovered a fruit-specific carotenoids biosynthesis pathway, triggered at the onset of tomato ripening. The analysis of light signaling mutants such as *hp* and ripening mutants such as *nor*, *rin*, *Nr* highlighted that multiple loci modulate carotenoid levels in tomato (Bramley et al., 1993).

While classical breeding approaches used loci such as *hp* and *beta* to increase lycopene and β-carotene, respectively in tomato, the identification of carotenoid biosynthesis genes provided an impetus to enhance the carotenoids by transgenic manipulations. These studies coupled with other studies revealed carotenoid accumulation in tomato is regulated at multiple steps involving transcriptional, post-transcriptional, post-translational, protein-protein interaction, and feedback regulation by end products (Liu et al., 2015; Stanley and Yuan, 2019). The transcriptional regulation of carotenoid biosynthesis is modulated by multiple transcription factors (TFs), some belonging to MADs box family such as *RIN*, *TAGL1*, *FUL1*, and *FUL2*, which by directly binding to the promoters of early carotenoid biosynthesis pathway genes, positively regulate lycopene accumulation (Vrebalov et al., 2009; Fujisawa et al., 2013, 2014; Shima et al., 2013).

The positive regulation of carotenoids biosynthesis is also mediated by other TFs regulating light or hormonal signaling. The suppression of *SlAP2a* (ERF) and *slHB-1* (HD-Zip TF) expression inhibited lycopene accumulation probably by suppressing ethylene biosynthesis (Lin et al., 2008; Karlova et al., 2011). The light signaling TF, *SlPIF1a* binds to the phytoene synthase promoter and represses its expression under the low red/far-red ratio (Llorente et al., 2016). The increase in carotenoid accumulation in tomato fruits is not limited to the direct promotion of carotenoids biosynthesis, the other light-signaling regulators such as *hp1* (*DDB1*) and *hp2* (*DET1*) mutants increase carotenoids by an increase in plastid number, plastid size, and carotenoids sequestration protein (Azari et al., 2010; Levin et al., 2003; Kilambi et al., 2013).

Compared to carotenoids, little is known about the regulation of other nutraceuticals in tomato such as folate. Though, folate biosynthetic pathway in plants has been deciphered (Hanson and Gregory, 2011; Gorelova et al., 2017a), its regulation at the transcription level remains unknown. In tomato fruits, the folate levels widely vary among different cultivars, besides the onset of ripening induces a reduction in folate level (Upadhyaya et al., 2017). Similarly, the expression of key folate biosynthesis genes *GTP cyclohydrolase I* (*GCHI*), and *aminodeoxychorismate synthase*, (*ADCS*), and *aminodeoxychorismate ligase 1* declined during ripening (Waller et al., 2010). In the absence of information about regulatory factors, the fortification of folate levels in tomato, as well in other crop plants, was achieved by transgenic overexpression of folate biosynthesis genes *GCHI*, and *ADCS* alone or in combination with downstream pathway genes (de la Garza et al., 2004, 2007; Storozhenko et al., 2007; De Lepeleire et al., 2018; Liang et al., 2019).

The EcoTILLING of folate biosynthesis genes revealed little polymorphism among tomato cultivars indicating that folate level is governed by complex regulation of cellular homeostasis (Upadhyaya et al., 2017). Likewise, folate biosynthesis genes were recalcitrant to EMS-mutagenesis (Gupta et al., 2017), as mutations in key folate biosynthesis genes lead to embryo lethality (Gorelova et al., 2017b). The QTL analysis of rice found no QTL associated with folate biosynthesis genes, and folate level was linked with a *folate transport protein*, *serine hydroxymethyl transferase*, and *folate hydrolase* genes (Wei et al., 2014). The QTL analysis of maize identified a *SAM-dependent methyltransferase* and a *transferase protein*-containing folate-binding domain as putative genes determining folate level in kernels (Guo et al., 2019). In low- and high-folate clones of wild potato (*Solanum boliviense*) the folate levels positively correlated with the expression of *γ-glutamyl hydrolase 1* (*GGH1*) (Robinson et al., 2019).

The above studies though indicated that the C1 metabolism and folate degradation determines folate levels, but did not identify any regulatory gene modulating folate levels. Nonetheless, the steady decline in transcripts of folate biosynthesis genes during ripening in tomato fruits indicated a developmental regulation of folate biosynthesis (Waller et al., 2010).

However, the identification of TFs regulating folate biosynthesis remains elusive. In plants, R2R3-MYB-TFs control wide-ranging responses encompassing development, metabolism, and responses to stresses. Emerging evidences indicated that R2R3-MYB TFs also participate in tomato ripening. The overexpression of *SlMYB72* in tomato while did not affect carotenoid accumulation, its suppression leads to a reduction in lycopene and β-carotene level (Wu et al., 2020). Likewise, repression of *SlMYB70* accelerated fruit ripening, while overexpression delayed fruit ripening (Cao et al., 2020). The *SlMYB12* mutant alleles fail to accumulate naringenin chalcone in the peel leading to pink fruit phenotype in tomato (Fernandez-Moreno et al., 2016).

The mutations in *SlMYB117* in tomato cause a trifoliate (*tf*) phenotype and alter auxin homeostasis of developing leaflets (Naz et al., 2013; Martinez et al., 2021). However, its influence on the fruit phenotype was not examined. Using EMS-mutagenesis, we isolated a new allele of trifoliate, namely *tf-5,* in the Arka Vikas (AV) cultivar. We examined its influence on fruit ripening and compared it with other reported trifoliate alleles. The system analysis of mutant fruit using transcriptomic, proteomics, and metabolic approaches revealed that *tf-5* influenced a wide range of responses, thus affecting the overall cellular homeostasis of fruits. Particularly, the *tf-5* allele substantially upregulated health-promoting nutraceuticals, γ-aminobutyric acid, folate, and carotenoids in tomato fruits.

## Results

### Isolation of a new *trifoliate* allele

In a mutant screen of Arka Vikas (AV) cultivar, we isolated a non-phototropic mutant that bore trifoliate leaves. In backcrossed BC_1_F_2_ population, non-phototropic and trifoliate phenotypes segregated independently as monogenic recessive mutations. Homozygous BC_1_F_2_ trifoliate plants lacking non-phototropic trait were backcrossed to obtain BC_2_F_2_ plants (**Figure S1**) These BC_2_F_2_ trifoliate plants and their progeny lacked non-phototropic trait and were characterized in detail. In tomato, mutations in *R2R3-MYB117* TF cause a trifoliate phenotype. The mutant *MYB117* gene had a C1818T transition replacing glutamine with a stop codon and truncating protein at 365 amino acids (Q366*) compared to 417 amino acids in AV. The sequence alignment revealed that the mutated *MYB117* gene encoded a novel *tf* allele and was named *tf-5* (**Figure 1A**, **Figure S2; Table S1**). The overlaying of *tf-2*, *tf-3, tf-4*, and *tf-5* allele amino acid sequences on rat R2R3-MYB model (https://www.rcsb.org/structure/1MSE) and MYB117 protein revealed above *tf* alleles had good homology for the R2R3 domain (**Table S2**). Among tf alleles, *tf-5* showed a very wide shift in the structure of non-MYB intrinsically disordered regions at the C-terminus compared to WT.

**Figure 1.**
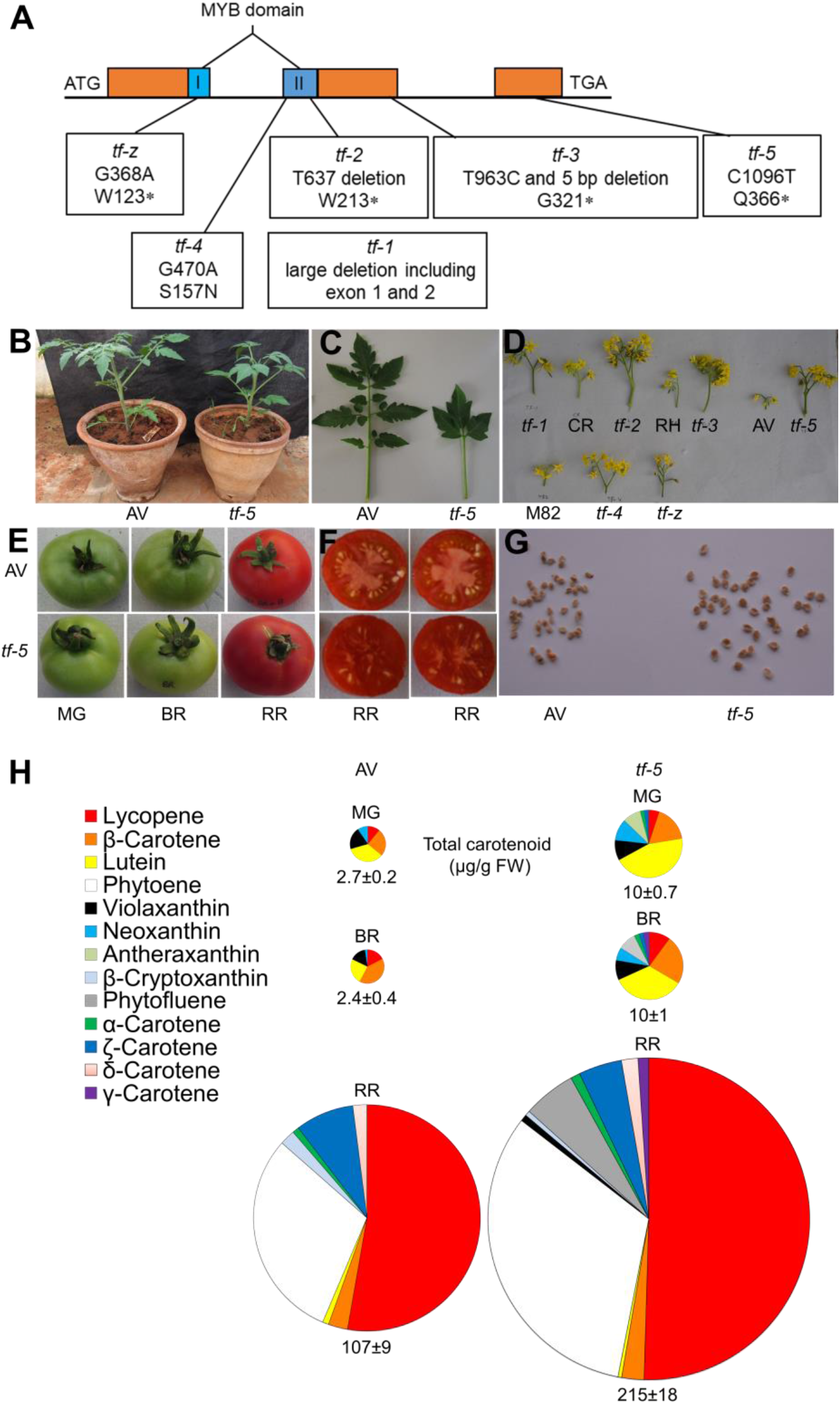
Genotypic, phenotypic, and carotenoids characterization of *tf-5.* (**A**) The location of different *tf* alleles in the *MYB117* gene (1976 bp, 417 amino acids). The *tf-5* mutation (C1818T) truncates the protein (Q366*). The MYB domains are shown in blue. Asterisks indicate the stop codon. (**B-G**) Phenotypic characterization of *tf-5* and its wild type (Arka Vikas –AV). (**B**) Plant stature. (**C**) Leaf morphology. (**D**) Inflorescence of different *tf* alleles with respective WT. (**E**) Fruit phenotype at mature green (MG), breaker (BR), and red-ripe (RR) stages of ripening. (**F**) Cut-halves of RR fruits. (**G**) Increased seeds size in *tf-5*. (**H**) The proportional amounts of different carotenoids in *tf-5* and AV fruits are depicted in the pie diagram. The values below the respective pie indicate the total carotenoid content (μg/g FW). Note that the area of the pie is proportional to the total carotenoid amount. The carotenoids data are means ± SE (n ≥ 4). See Table S4 for individual carotenoid levels and significance.

### *tf-5* fruits have high carotenoids levels

Since *tf-5* severely affected leaf and inflorescence development including smaller plant stature (**Figure 1B-D; Table S3**), we examined its effect on fruit ripening. The *tf-5* did not affect the duration of attainment of mature-green (MG), breaker (BR), and red-ripe (RR) stages of ripening, but fruits were more oblong-shaped and slightly smaller (82%) than AV. Markedly, *tf-5* fruits were darker-red, with red-colored placenta, and seeds were larger than AV (170%) (**Figure 1E-G).** However, similar to *tf-z* (Naz et al., 2013), it developed fewer seeds than WT. The darker-red colour indicated an elevation in the carotenoids level in *tf-5* fruits. Consistent with this, lycopene level in *tf-5* RR fruit was nearly 1.9-fold higher than AV. Wholly; at all ripening stages, most carotenoids were higher in *tf-5* fruits than AV (**Figure 1H**). We then examined carotenoid levels in other *tf* alleles (**Table S4**). In a like manner, *tf-*2 also had nearly 1.6-fold higher lycopene levels. The *tf-1* fruits were tangerine-coloured and accumulated little lycopene, but had a normal level of β-carotene and lutein (**Figure S3**; **Table S4**). Compared to other *tf* alleles, RR fruits of *tf-5* had higher °Brix, firmness, and bigger seeds than its wild type (**Figure S4**).

### *tf-5* fruits are rich in folate content

Considering that fruits of *tf-5* were enriched in carotenoids, we examined whether the *trifoliate* mutation also influenced folate levels. The folate level in *tf-5* fruits at all ripening stages was markedly high (3.8-fold at RR) than AV (**Figure 2A**). We then examined folate and pterin levels in other *tf* alleles (**Table S5**). Alike carotenoids, the RR fruits of *tf-2* also had higher folate (1.55-fold) levels than WT. Barring *tf*-*3*, *tf-4* and *tf-z* also had higher folate levels in RR fruits. We could not judge the relative folate level of *tf-1* as its WT background is not known. Corroborating higher folate, its precursor *p*-aminobenzoic acid (*p*ABA) was also high in *tf-5* fruits. A similar correlation between *p*ABA and folate level was seen in *tf-2*. The levels of other precursors- monopterin, and neopterin, were below detection level and 6-hydroxymethylpteridine (HMPt) was detected only in few alleles and/or WTs. Pterin-6-carboxylic acid (p6C), an intermediate in folate recycling, was detected in all alleles, and its level in *tf-2*, and *tf-5* was lower than respective WT (**Figure 2A, Table S5**).

**Figure 2.**
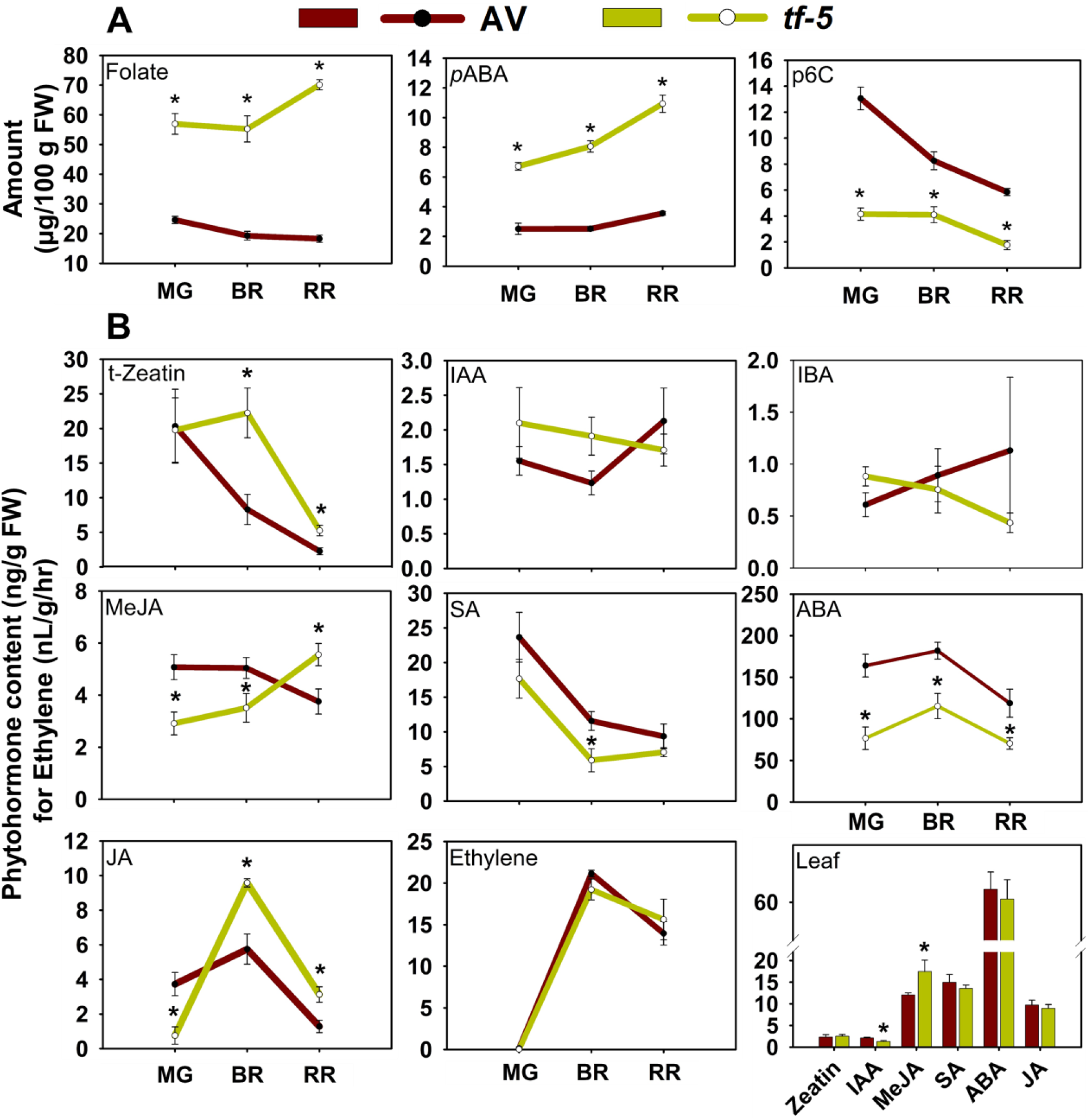
Folate, *p-*aminobenzoic acid (*p*ABA), pterin-6-carboxylate (p6C), and phytohormones levels in AV 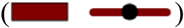 and *tf-5* 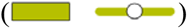 fruits at different ripening stages. (**A**) The level of total folate, *p*ABA, and p6C. (**B**) Phytohormones level in AV and *tf-* The levels of phytohormones were measured by LC-MS, except ethylene, which was measured by gas-chromatography and expressed as nL/g/hr. The bar diagram at the bottom right shows levels of different phytohormones in AV and *tf-5* leaves (5th node of 7-week-old plants). Data are means ± SE (n =5), * = p ≤ 0.05. An asterisk (*) shows a significant difference compared to AV. *Abbreviations*: IAA, indole acetic acid; IBA, indole butyric acid; MeJA, methyl jasmonate; SA, salicylic acid; ABA, abscisic acid, and JA, jasmonic acid.

*In vivo* folate exists as different vitamers whose abundances vary during development. In *tf-5* and AV fruits, four folate vitamers were detected. Among these, THF was a minor vitamer and detected only at a few stages. The abundance of folate vitamers varied during ripening. While the 5-CH_3_-THF level remained the same in *tf-5*, the levels of 5,10-CH^+^-THF increased during ripening. In contrast, the reduction in total folate levels in WT was due to the decline in 5-CH_3_-THF levels (**Figure S5A**) In the leaf of *tf* mutants, the folate levels were higher in *tf-4, tf-z,* and *tf-5*, while it was lower in *tf-2* (**Figure S5B**). Contrasting to tomato, leaves of Arabidopsis *MYB117* mutant (*lof1*) and its orthologue *MYB105* (*lof2*) (**Figure S6**) had reduced folate levels in WT (**Figure S5C**) (Lee et al., 2009). Unlike folate, there was no effect of *tf-5* on carotenoids levels of leaves except antheraxanthin, which was somewhat low (**Figure S5D-E**). Noting that *tf*-4 and *tf*-z are EMS-mutagenized M_2_ lines, and *tf*-3 had no folate stimulation, in later sections we compared *tf-5* mainly with *tf*-2 (LA0579) which is reportedly backcrossed.

### Hormonal homeostasis is altered in *tf-5* fruits

Taking cognizance that *tf*-2 affects auxin biosynthesis, transport, and signalling in leaf (Martinez et al., 2021), we examined, whether *tf-5* altered hormonal homeostasis in fruits and leaves. Surprisingly, while *tf-5* only moderately influenced ethylene emission, it up-regulated (↑) or down-regulated (↓) other hormones. In *tf-5* fruits, ABA and salicylic acid (SA) levels were lower, while IAA and IBA levels were nearly similar to AV. Jasmonic acid (JA) (MG↓, BR↑, RR↑), methyl jasmonate (MeJA) (MG↓, BR↓, RR↑), and zeatin (BR↑, RR↑) levels differed from AV in a stage-specific fashion (**Figure 2B**). Levels of GA and brassinosteroids were presumably lower than the limits of detection, henceforth could not be determined. In the *tf-5* leaf, the IAA level was lower and the MeJA level was higher than AV.

### Mutation in *MYB117* upregulates amino acid levels in *tf-5*

The higher levels of folate in *tf-5* implied that altered folate levels might also affect the overall metabolic homeostasis. Equally, MYB being a broad-spectrum transcriptional regulator, *tf-5* may directly affect homeostasis. Consistent with this, out of 77 detected primary metabolites, most metabolites levels in *tf-5* substantially differed from AV (**Figure 3A**). In accordance therewith, the metabolite cluster of *tf-5* fruits in PCA was distinct from AV (**Figure 3B**). The hierarchical classification of metabolites revealed that the majority of the amino acids and amines were substantially higher in *tf-5* than AV (**Figure 3C, Dataset S1**). Notably, the levels of glutamate, tryptophan, and γ-aminobutyric acid (45-fold) were massively upregulated in RR fruits. Consistent with higher °Brix in *tf-5* fruits, sucrose level was higher at RR. The levels of sugar derivatives and fatty acids levels were affected either way during ripening with lower or higher levels at one or two stages in *tf-5.* Among the TCA cycle components, only cis-aconitate significantly varied, it was at a lower level in early ripening stages and became equal to AV at the RR stage. The regulation patterns were also variable as seen for oxalic acid (MG↑, BR↓, RR≈).

**Figure 3.**
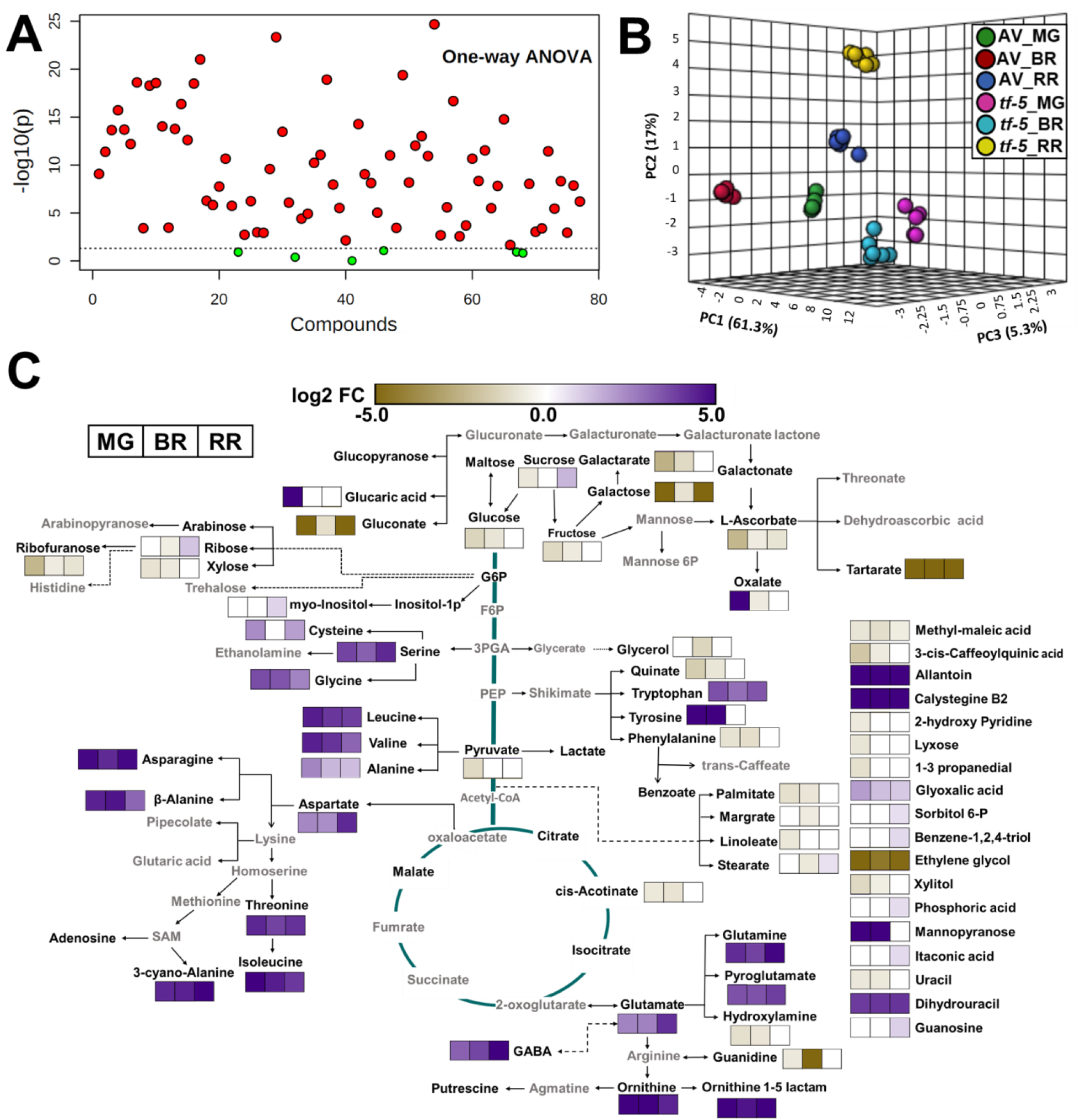
The levels of primary metabolites in AV and *tf-5* fruits at different ripening stages. **(A)** Important features (solid red circle) selected by One-way ANOVA plot (P≤0.05). One-way ANOVA showed that 71 of the 77 metabolites present at MG, BR, and RR differed in AV and *tf-5*. (**B**) Principal component analysis (PCA) of AV and *tf-5* at MG, BR, and RR stage. The PCA was plotted using MetaboAnalyst 4.0. (**C**) The metabolic shifts in *tf-5* fruits during ripening in comparison to AV. The relative changes in the metabolite levels at MG, BR, and RR stage in *tf-5* fruits were determined by calculating the *tf-5*/AV ratio at respective ripening phases. Only significantly changed metabolites are depicted on the metabolic pathway (Log2 fold ≥ ± 0.584 and p-value ≤ 0.05). Bold black letters indicate the identified metabolites. Grey letters indicate metabolites below the detection limit. The white box indicates no significant change in metabolite level. Data are means ± SE (n ≥ 5), p ≤ 0.05. See Dataset S1 for detailed metabolite data.

### Systems analysis of *tf-5*

Considering the broad-spectrum effect of *tf-5* on metabolome, we profiled the proteome of AV and *tf-5* fruits. Consistent with the wide-ranging effect of *tf-5* on metabolome, it also influenced the proteome profile. Label-free quantification of proteins identified 516, 643, and 531 differentially expressed proteins (Log2 fold ± 0.58, p-value ≤ 0.05) in *tf-5* at MG, BR, and RR respectively. At MG and RR, most proteins were upregulated (337↑, 179↓, MG; 257↑, 274↓, RR), while in BR most were downregulated (270↑, 373↓, BR) (**Dataset S2, S3**). The complement of up/down-regulated proteins varied in a stage-specific fashion, and only a few proteins overlapped at two or more stages (**Figure S7A-B**). The overlapping upregulated proteins were mainly storage proteins, proteinase inhibitors, and chitinases, whereas proteases were downregulated. The GO classification of differentially expressed proteins was in conformity with the wide-ranging influence of *tf*-5 on cellular homeostasis (**Figure S7C**).

Considering that *tf*-5 affected homeostasis, with a very prominent effect on carotenoids and folate at RR, we compared the transcriptome of WT and *tf-5* at RR. Among significantly affected transcripts (Log2 fold ± 0.58, p-value ≤ 0.05), 2531 and 1273 were upregulated and downregulated respectively (**Dataset S4**, **Figure S8**). The GO classification of up/downregulated transcripts revealed that major upregulated classes (> 100 transcripts) were enzyme classifications, RNA biosynthesis, protein modifications, solute transport, and cell wall organization (**Figure S8**). The *tf-5* also influenced a large number of TFs (187↑, 167↓). Among TFs, the *MYB* was the major group (30↑, 5↓) (**Figure S8**) followed by *bHLH* (15↑, 6↓), *bZIP* (14↑, 7↓), and *ERF* (11↑, 7↓) (**Dataset S4**).

The Mapman and LycoCyc depiction of proteome and transcriptome revealed the influence of *tf-5* on almost all aspects of cellular metabolism (**Figure S9-S10**). Comparison of metabolome, proteome, and transcriptome at the RR stage brought forth the complexities of cellular homeostasis, with some direct correlations. Conforming to enhanced firmness, several cell wall modifying enzymes- expansins, pectin esterases, and pectinacetyl esterases were downregulated in *tf-5* (**Figure 4A, Dataset S3**), while those modulating redox metabolism and fasciclin like arabinogalactan proteins were upregulated. Congruent with higher °Brix level in *tf-5*, sucrose synthase protein and its transcript (Solyc12g009300, RR) was upregulated, while both cytosolic and vacuolar invertases were downregulated. The distinct upregulation of the amino acids in *tf-5* seems to involve lowering of t-RNA charging at BR and RR, as several aminoacyl-tRNA synthases were downregulated {[10↓ BR, 11↓, 1↑ RR, in proteome], [5↓, in transcriptome]} (**Figure 4B, Dataset S3**). Amino acid biosynthesis and degradation were similarly affected in *tf-5* (**Figure 4C-D**).

**Figure 4.**
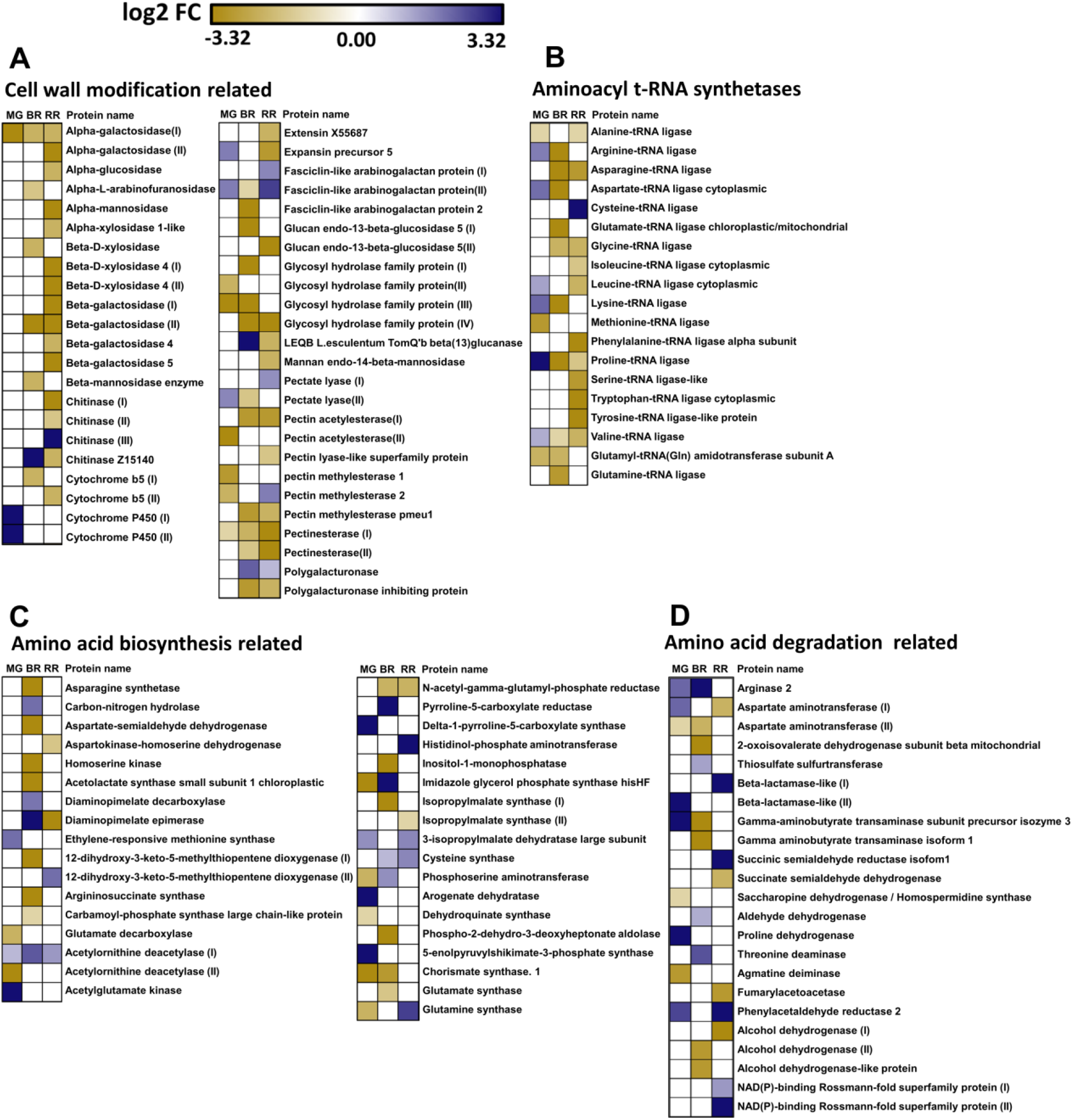
Heat maps representation of differentially expressed proteins affecting metabolic pathways in *tf-5* during fruit ripening. **(A)** Cell wall modification. **(B)** Aminoacyl t-RNA synthetases. **(C)** Amino acid biosynthesis. **(D)** Amino acid degradation. Only significantly different proteins (Log2 fold ≥±0.58, p ≤ 0.05) are shown in heat maps. Data are means ± SE (n = 3), p ≤ 0.05. See Dataset S2, S3 for detailed proteome data.

A global comparison of differentially expressed proteins with respective transcripts showed little correlation between them. Among downregulated proteins, 25 proteins and transcripts showed correlation, while 44 upregulated proteins correlated with respective transcripts. Strikingly, for 16 upregulated and 31 downregulated proteins, respective transcripts were down- and up-regulated, respectively (**Dataset S2**). The mismatch between proteome and transcriptome was noticeably evident for secondary metabolism, where *tf-5* affected several transcripts, but only a few proteins at RR (**Figure S9**).

### Several proteins/transcripts modulating hormonal levels are affected in *tf-5*

In consonance with the elevation of jasmonate levels, the levels of several proteins/transcripts regulating jasmonate synthesis were modulated in *tf-5.* Particularly *lipoxygenases*, *CytP450s*, *12-oxophytodienoate reductases* transcripts along with lipoxygenases proteins were upregulated. For the MeJA pathway, the *jasmonate-o-methyl transferase* transcript was upregulated and *methyl jasmonate esterase* transcript and protein were downregulated (**Figure 5A**). Correspondingly, the transcripts of four TFs related to jasmonate were also upregulated (Solyc08g036640, Solyc12g009220, Solyc07g042170, and Solyc03g118540). Conversely, the genes contributing to ABA biosynthesis were downregulated, particularly violaxanthin epoxidase with reduced protein and transcript levels. Contrarily, the genes related to ABA degradation; specifically three isoforms of *abscisic acid 8’-hydroxylase* were upregulated (**Figure 5B**). Though ACO1 protein at BR and RR, and *ACO2* transcripts at RR were upregulated, the ethylene emission from *tf-5* was nearly similar to WT. Oddly, ACO6 protein, with yet undetermined function (Houben and Van de Poel, 2019), was also upregulated in *tf-5* at BR (**Dataset S2**).

**Figure 5.**
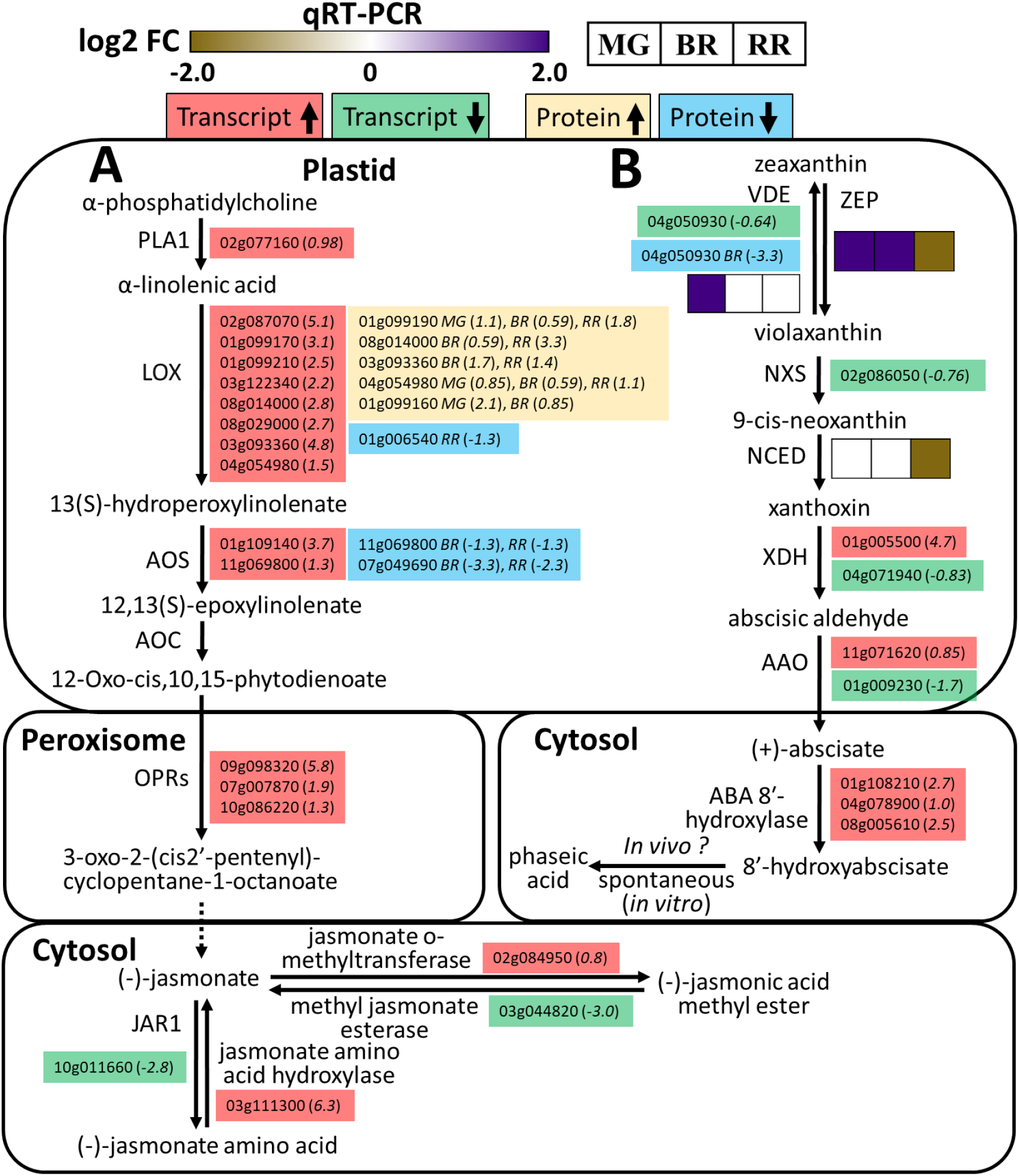
Jasmonate biosynthesis and ABA biosynthesis/degradation pathway showing changes in proteins/transcripts abundance in *tf-5*. **(A)** Jasmonic acid pathway. **(B)** ABA biosynthesis/degradation pathway. The SOLYC prefix before each gene/protein number is removed for convenience. Upregulated (↑) and downregulated (↓) transcripts and proteins are shown in different colors. The transcriptome data are for the RR stage only. The value in parentheses next to the transcript number represents the log2 fold change. The *MG*, *BR*, and *RR* in italics placed next to the protein number with values in parenthesis depict log2 fold change at that ripening stage. The heat maps show log2 fold change in the transcript of respective gene determined by qRT-PCR at MG, BR, and RR stage. Only significantly different proteins/transcripts (Log2 fold ≥ ±0.58, p ≤ 0.05) are shown See Dataset S2, S3 and Dataset S4 for detailed proteome and transcriptome data, respectively. Data are means ± SE (n = 3), p ≤ 0.05. *Abbreviations*: PLA1, phospholipase A1, LOX, lipoxygenase; AOS, allene oxide synthase; AOC, allene-oxide cyclase; OPRs, 12-oxophytodienoate reductase; JAR1, jasmonate amino acid synthetase; VDE, violaxanthin de-epoxidase, ZEP, zeaxanthin epoxidase; NXS, neoxanthin synthase; NCED, 9-cis-epoxycarotenoid dioxygenase; XDH xanthoxin dehydrogenase; AAO, abscisic aldehyde oxidase; ABA 8′-hydroxylase, (+)-abcisic acid 8′-hydroxylase.

### *tf-5* upregulates carotene isomerases in RR Fruits

In transcriptome/proteome, several proteins and transcripts of the MEP pathway, which provides carotenoids precursors, as well as of carotenoid biosynthetic pathway were detected (**Figure 6**). In MEP-pathway, isopentenyl diphosphate Δ-isomerase protein, *farnesyl diphosphate synthase*, and *geranylgeranyl diphosphate synthase* (*GGPPS2),* transcripts were upregulated. In the carotenoid biosynthesis pathway, two key proteins ζ-carotene isomerase (ZISO) and carotenoid isomerase (CRTISO) were upregulated in *tf-5* at RR. As observed in earlier studies (Kilambi et al., 2017, 2021), the transcripts contributing to lycopene synthesis had little correlation with increased protein levels at the RR stage. The transcript/proteins profiles also revealed modulation of plastoquinone synthesis in *tf-5*, particularly, upregulation of *tyrosine aminotransferase*. The *tf-5* also showed higher levels of carotenoids sequestration protein, PAP3 (Solyc08g076480) at RR. Curiously, remorin1 (Solyc03g025850), which reportedly upregulates carotenoids levels in tomato fruits (Cai et al., 2018), was high in *tf-5* at BR and RR (**Dataset S2**).

**Figure 6.**
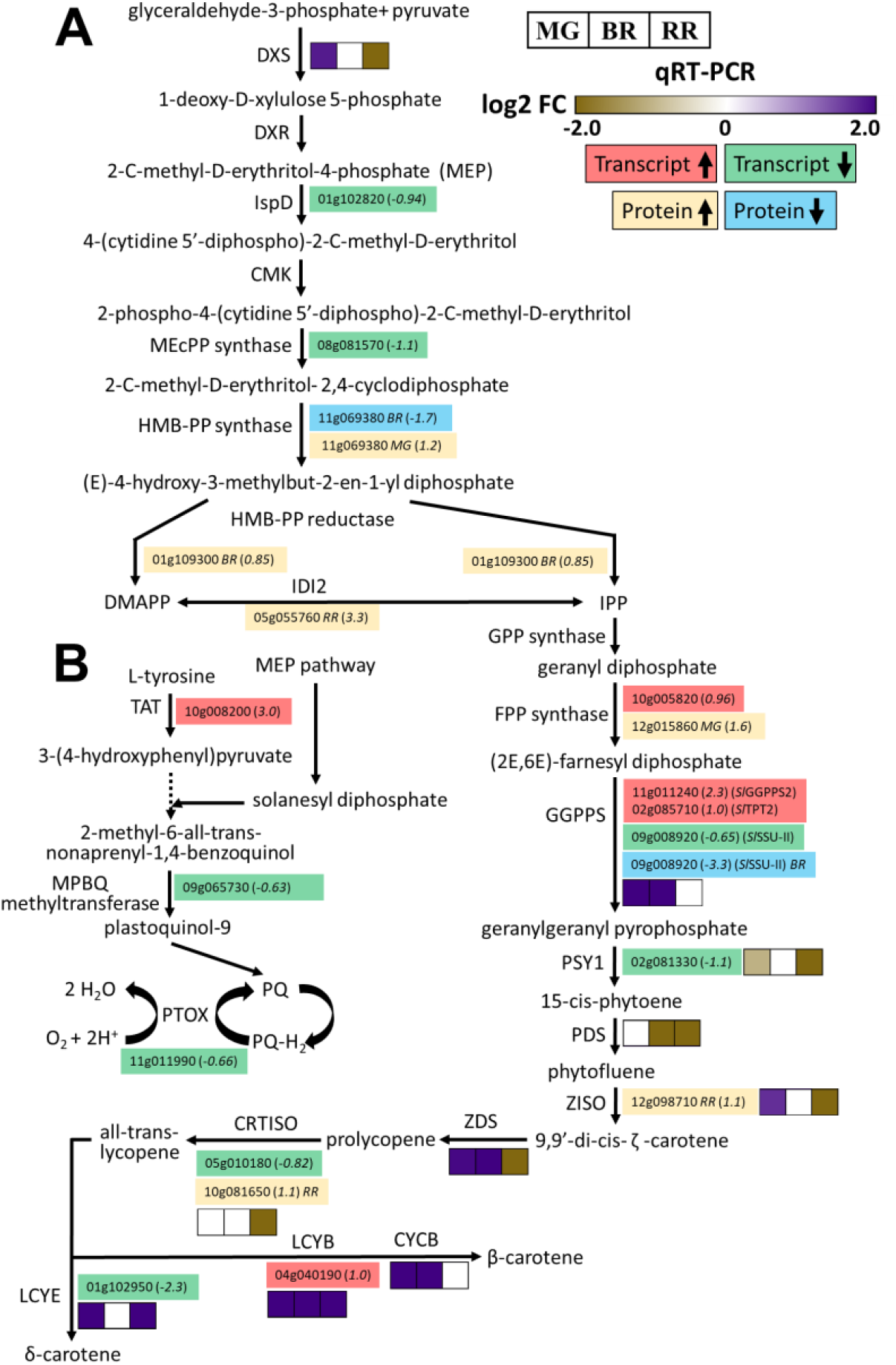
Plastoquinone, methylerythritol, and carotenoid biosynthesis pathways showing changes in proteins/transcripts abundance in *tf-5.* The SOLYC prefix before each gene/protein number is removed for convenience. Upregulated (↑) and downregulated (↓) transcripts and proteins are shown in different colors. The transcriptome data are for the RR stage only. The value in parentheses next to the transcript number represents the log2 fold change. The *MG*, *BR*, and *RR* in italics placed next to the protein number with values in parentheses depict log2 fold change at that ripening stage. The heat maps show log2 fold change in the transcript of respective gene determined by qRT-PCR at MG, BR, and RR stage. Only significantly different proteins/transcripts (Log2 fold ≥±0.58, p ≤ 0.05) are shown in heat maps. See Dataset S2, S3 and Dataset S4 for detailed proteome and transcriptome data, respectively. Data are means ± SE (n = 3), p ≤ 0.05. *Abbreviations*: DXS, 1-deoxy-D-xylulose-5-Phosphate synthase; DXR, 1-deoxy-D-xylulose-5-phosphate reductoisomerase; IspD, 2-C-methyl-D-erythritol-4-phosphate cytidylyltransferase; CMK, 4-(cytidine 5’-diphospho)-2-C-methyl-D-erythritol kinase; MEcPP synthase, 2-C-methyl-D-erythritol 2,4-cyclodiphosphate synthase; HMB-PP synthase, 4-hydroxy-3-methylbut-2-en-1-yl diphosphate synthase; IDI2, isopentenyl-diphosphate Δ-isomerase; GPP synthase, geranyl diphosphate synthase; FPP synthase, farnesyl diphosphate synthase; GGPPS, geranylgeranyl diphosphate synthase; PSY1, phytoene synthase; PDS, phytoene desaturase; ZISO, ζ-carotene isomerase; ZDS, ζ-carotene desaturase; CRTISO, carotenoid isomerase; LCYE, lycopene ε-cyclase; LCYB, lycopene β-cyclase; CYCB-lycopene β-cyclase (chromoplastic); TAT, tyrosine aminotransferase; MPBQ methyltransferase, 2-methyl-6-solanyl-1,4-benzoquinone methyltransferase; PTOX, plastid terminal oxidase; IPP, isopentenyl pyrophosphate; DMAPP, dimethylallyl diphosphate; PQ, plastoquinone.

### *tf-5* fruits show altered C1 metabolism and reduced GGH enzyme activity

For folate biosynthesis, perhaps due to low abundance, only *dihydropteroate synthase* (Solyc05g012090↓) and *dihydrofolate reductase* (*DHFR)* (Solyc04g074950↑) transcripts (**Dataset S4**), but no proteins were detected. Nonetheless, qRT-PCR indicated upregulation of several folate biosynthesis genes at one or more ripening stages in *tf-5* and *tf-2* (**Figure 7A, Figure S11A).** Notably, *GCHI* and *ADCS*, the key genes contributing to the first step of folate biosynthesis were upregulated in *tf-5*. We also examined whether *trifoliate* mutation affected the expression of the *MYB117* gene, which in turn may have led to stimulation of folate synthesis genes. The expression of *MYB117* was significantly higher at the MG and RR in *tf-5 (***Figure 7B**), and *tf-2* fruits and leaves (**Figure S11**). We then examined *in silico* whether promoters of folate and carotenoid biosynthesis genes have MYB binding motif using PLACE software. The majority of these genes had one or more MYB binding motif in their promoters (**Figure S12**). Since *MYB117* promoter too has a MYB binding motif, its upregulation may be related to its auto-regulation alone or with other TFs.

**Figure 7.**
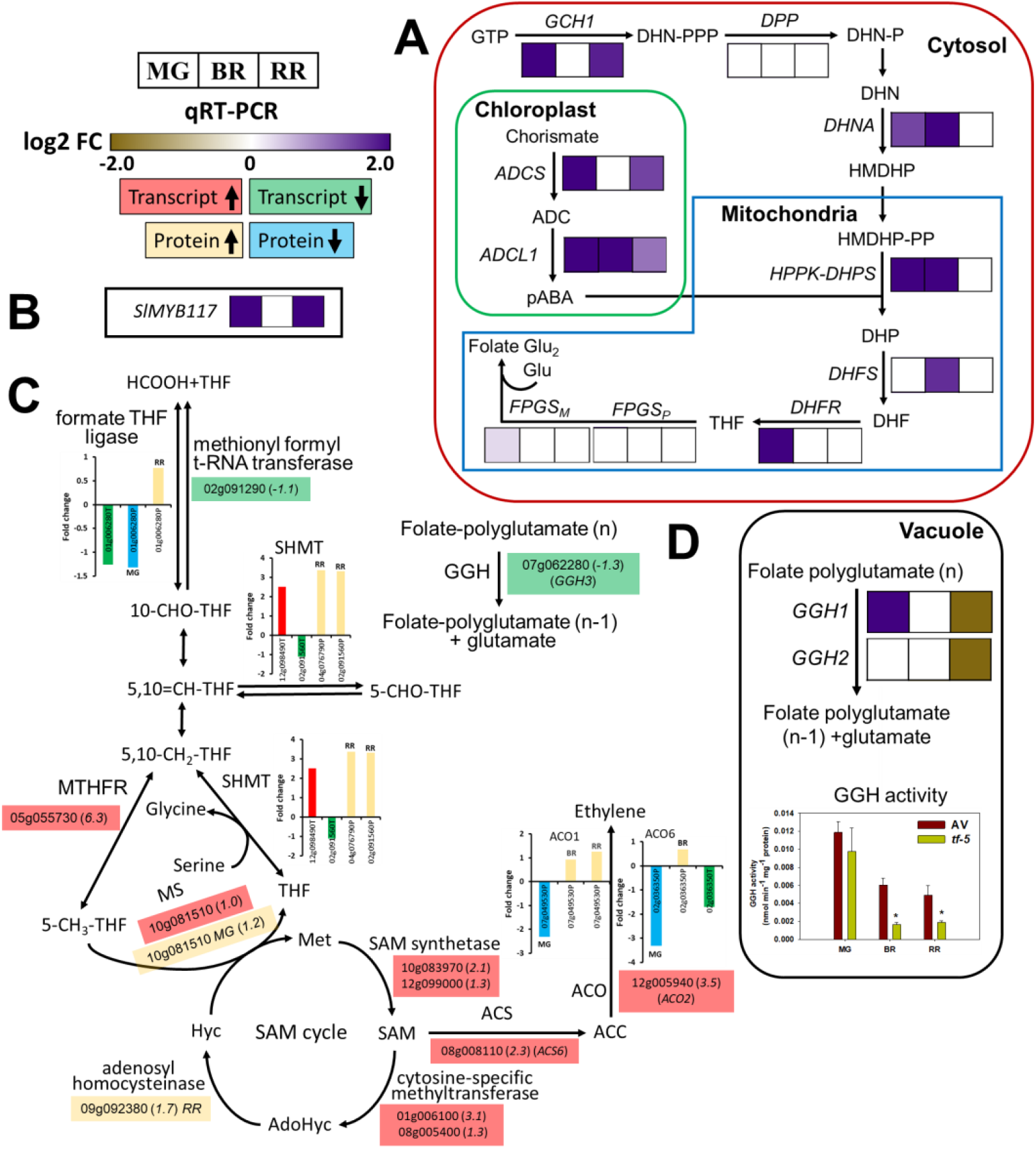
Folate biosynthesis and C-1 metabolism pathway in *tf-5*. **(A-B)** The heat maps show log2 fold change in transcripts of folate biosynthesis pathway genes (**A**), and *MYB117* (**B**) in *tf-5* determined by qRT-PCR at MG, BR, and RR stage. (**C)** Changes in abundance of transcripts and proteins of C-1 metabolism in *tf-5.* (**D**) Changes in *GGH1*, *GGH2* transcripts, and total GGH enzyme activity in *tf-5* at MG, BR, and RR stage. The SOLYC prefix before each gene/protein number is removed for convenience. Upregulated (↑) and downregulated (↓) transcripts and proteins are shown in different colors. The transcriptome data are for the RR stage only. The value in parentheses next to the transcript number represents the log2 fold change. The log2 fold changes in protein abundance are shown as the histograms. The MG, BR, and RR placed next to the protein number show log2 fold change at that ripening stage. Only significantly different proteins/transcripts (Log2 fold ≥ ± 0.58, p ≤ 0.05) are shown in heat maps. See Dataset S2, S3 and Dataset S4 for detailed proteome and transcriptome data, respectively. Data are means ± SE (n = 3), p ≤ 0.05. *Abbreviations:* GCH1, GTP cyclohydrolase I; DPP, dihydroneopterin (DHN) triphosphate diphosphatase; DHNA, DHN aldolase; ADCS, aminodeoxychorismate (ADC) synthase; ADCL, aminodeoxychorismate lyase; HPPK-DHPS, 6-hydroxymethyl-7,8-dihydropterin (HMDHP) pyrophosphokinase (HPPK) and dihydropteroate (DHP) synthase; DHFS, dihydrofolate (DHF) synthase; DHFR, DHF reductase; FPGS, folylpolyglutamate synthase (p-plastic, m-mitochondrial), GGH, γ-glutamyl hydrolase; SHMT, serine hydroxymethyl transferase; MTHFR, methylenetetrahydrofolate reductase; MS, methionine synthase; SAM synthetase, S-adenosylmethionine synthetase; ACS, 1-aminocyclopropane-1-carboxylic acid synthase; ACO, 1-aminocyclopropane-1-carboxylate oxidase; Glu, glutamate; *p*ABA, *p*-aminobenzoate; THF, tetrahydrofolate, -P, phosphate;P2, diphosphate; P3, triphosphate; 10-CHO-THF, 10-formyl-THF; 5,10=CH-THF, 5,10 methenyl-THF; 5,10-CH_2_-THF, 5,10 methylene-THF; 5-CHO-THF, 5-formyl-THF; Met, methionine; SAM, S-adenosylmethionine; AdoHyc, adenosyl-L-homocysteine; Hyc, homocysteine.

Mapman analysis revealed that *tf-5* also influenced the C1 metabolism modulated by folate (**Figure S9**). Markedly, several proteins participating in C1 metabolism were upregulated in *tf-5,* mainly at the RR stage. The S-adenosylhomocysteinase, formate-tetrahydrofolate ligase, methylenetetrahydrofolate reductase (MTHFR), and serine hydroxymethyl transferase (SHMT) proteins were upregulated in *tf-5* (**Figure 7C**). Oddly, the transcripts related to C1 metabolism did not match with the above protein isoforms and showed both up- and down-regulation. Specifically, upregulated transcripts were *methionine synthase*, *SAM synthase*, *SHMT*, *MTHFR*, and *Cytosine-dependent-methyltransferase*.

The transcriptome profiling of *tf-5* revealed a reduction in the folate catabolism gene, *GGH3* (Solyc07g062280), which though encodes an inactive protein, can dimerize with GGH1 and GGH2 and reduces their activity (Akthar et al., 2008). The qRT-PCR analysis showed at RR both *GGH1* and *GGH2* transcripts were downregulated in *tf-5* (**Figure 7D**). The enzymatic assay of GGH activity in WT and *tf-5* fruits showed a substantial reduction in the deglutamylation of folate in *tf-5*. Consistent with lower GGH activity, the polyglutamylation was higher in *tf-5* fruits at the RR stage of ripening (**Figure S13**). In a like manner, *tf-2* also showed lower levels of *GGH* transcripts and enzyme activity (**Figure S11**). To validate the role of GGH in modulating cellular folate levels, we generated gene-knockouts for *GGH1* and *GGH2* in tomato. Similar to *tf-5*, both *GGH1* and *GGH2* knockouts in T_0_ showed 2-3 fold higher folate levels in RR fruits (**Figure S14**). We are advancing these lines to validate the above results in T_1_ and T_2_ generation.

## Discussion

### Phenotypic effect of *tf-5* ensues from modifications of multiple processes

The new trifoliate allele *tf-5*, displayed altered inflorescence architecture, had deep-red colored fruits and bigger-sized seeds than the wild type (AV). It also had reduced placenta, higher °Brix, and firmer fruits. Ostensibly, besides leaf architecture, the *tf-5* also affected the fruit phenotype. Contrarily to leaf, where five *tf* alleles (barring *tf-4*) show similar phenotype, their action in fruit differs, at least on folate and carotenoids levels. Among six *tf* alleles, *tf-2* and *tf-5* affected a similar subset of the responses, viz. increase in carotenoids and folate levels in fruits. However, this similarity may be specific to the fruits, as, in leaf, *tf-2*, and *tf-5* show the opposite effect on the folate level.

The physiochemical analysis of the *tf-2* leaf indicated that altered auxin transport and biosynthesis are important contributors to its trifoliate phenotype (Naz et al., 2013; Martinez et al., 2021). Analogous to *tf-2*, the *tf-5* leaf showed reduced auxin level, and besides had a higher MeJA level. Contrarily, auxin levels were not altered in *tf-5* fruits, but RR fruits had enhanced JA and reduced ABA levels. In Arabidopsis, reduced ABA level in *abi5* and *aba2* mutant silique is correlated with the formation of larger size seeds (Cheng et al., 2014; Li et al., 2019). As *tf-5* fruits have reduced ABA level that may have contributed to increased seed size.

In plants, TFs such as MYB regulate diverse developmental responses. In tomato, the MYB family consists of nearly 127 genes, and several of these including *MYB117* express in the fruit (Li et al., 2016). Similar to the *tf-2* leaf (Martinez et al., 2021), the expression of *MYB117* is higher in *tf-5* and *tf-2* fruits. In plants, MYBs and bHLHs TFs have unique properties, signified by their propensity to act as homo- or hetero-dimers (Pireyre and Burow, 2015). Thus, MYBs can form MYB/bHLH complexes, or even associate with other proteins to modulate the cellular processes. Consistent with this, *tf-5* shows upregulation of a large number *MYBs*, *bHLH,* and other TFs in fruits. In turn, this seemingly influenced overall homeostasis, which is reflected by wide changes in transcriptome, metabolome, and proteome.

### Reduced t-RNA charging likely elevates amino acids levels

The metabolic shifts are considered as a penultimate manifestation of homeostasis alteration at the systems level. Corresponding with phenotypic alteration, the metabolome profile of *tf-5* was markedly different from WT, a feature seen with ripening-related TFs (Osorio et al., 2011), or hormonal mutants (Sharma et al., 2021). A distinctive feature of the shift was higher amino acid levels in *tf-5*. In parallel, the protein levels of several aminoacyl-tRNA synthetases in *tf-5* declined during ripening. Consecutively, reduced t-RNA charging may elevate free amino acids due to lowered flux towards protein synthesis.

Similar to amino acids, the firmness of *tf-5* fruits correlated to the lowering of the several cell wall softening enzymes, such as expansin, pectin esterases, β-D-xylosidases, α- and β-mannosidases. Conversely, upregulation of fasciclin-like arabinogalactan-proteins may add to the strengthening of cell walls (Huang et al., 2013). The higher °Brix level of *tf-5* may stem from an increase in sucrose synthase protein and transcript. However, these correlations may be fortuitous, as altered protein levels may not accurately reflect their *in vivo* activities, which is determined by other factors too. Nonetheless, on a broader scale, the metabolic shifts in *tf-5* were an outcome of wide changes in transcriptome and proteome.

### Upregulation of lipoxygenases may be responsible for increased JA levels

Being a climacteric fruit, tomato ripening is modulated by the upregulation of ethylene biosynthesis (Grierson, 2013). Though *tf-5* affected a wide range of responses, it did not influence ethylene emission. Contrarily, other hormones particularly JA, MeJA, and ABA levels were modulated by *tf-5*. Nonetheless, modulation of these hormones did not affect the duration of the transition from MG to BR in *tf-5*. In strawberry during early ripening, the ABA biosynthetic enzymes are suppressed, while ABA-degrading enzymes show higher activities (Kim et al., 2019). Analogous to strawberry, the transcripts of *ABA 8*′ *hydroxylases* are upregulated in *tf-5*, which in conjugation with downregulation of violaxanthin de-epoxidase protein/transcript likely contributed to reduced ABA level. Consistent with this, the level of violaxanthin, a precursor for ABA is high in *tf-5* fruits (Neuman et al., 2014).

In tomato, JA is necessary for *MYB21* transcription, which in turn regulates ovule development in flowers (Schubert et al., 2019). The high level of JA in *tf-5* may ensue from the upregulation of lipoxygenases (LOX) proteins and transcripts. Likewise, higher MeJA levels may arise from the upregulation of *jasmonate-o-methyl transferase* transcript and downregulation of methyl jasmonate esterase protein/transcripts.

### *tf-5* may be a gain of function mutation

Emerging evidences have indicated that carotenoid biosynthesis/accumulation in ripening tomato fruits is governed by multiple factors (Liu et al., 2015; Stanley and Yuan, 2019). Our *in silico* analysis of promoters revealed MYB binding sites in several genes related to carotenoids and folate biosynthesis. Thus, MYB117 may influence carotenogenesis by binding to the promoters alone or in concert with other TFs. In *Citrus reticulate* fruits, *CrMYB68* TF negatively regulates the expression of *β-carotene hydroxylase2* and *NCED5* by binding to their respective promoters (Zhu et al., 2017). In tomato, *MYB72* binds to *phytoene synthase 1* and ζ*-carotene isomerase* promoters; thereby its suppression by RNAi reduced the synthesis of lycopene in fruits (Wu et al., 2020). The contribution of *MYB72* to increased carotenoids in *tf-5* seems to be remote, as, in the *tf-5* transcriptome, the *MYB72* transcript level was not affected.

Considering that *tf-1*, with total deletion of the *MYB117* gene, is compromised in lycopene synthesis, a functional MYB may be needed for stimulation of carotenoid levels. Excepting *tf-3*, other *tf* alleles-*tf-2*, *tf-4*, *tf-z* showed elevated carotenoids indicating a linkage between *MYB117* mutation and carotenoids accumulation. Notably, these *tf* alleles enhanced levels of only a few carotenoids; none showed stimulation of the majority of carotenoids like *tf-5*. The varying carotenoid levels in *tf* alleles likely reflect the diversity in the expression of individual alleles and/or interaction with other partners. Likewise, in barley, mutations in the *Sln1* gene encoding DELLA protein lead to either overgrowth or dwarfing phenotypes, highlighting different alleles can diversely affect the phenotype (Chandler and Harding, 2013).

Since in the *tf-5* majority of differentially expressed genes were upregulated, the positive influence of *tf-5* on the transcriptome may be due to a gain of function. This gain of function may be imparted by C-terminus truncation, as in R2R3-MYBs, the non-MYB domains that constitute the intrinsically disordered region are important for the diversity in their activity (Millard et al., 2019). Since *tf-5*-MYB117 shows a major alteration in C-terminus structure, it is therefore likely that the above change in the intrinsically disordered region relieved the repressive action of MYB117. Seemingly above alteration in the C-terminus potentiated the wide-ranging effect of *tf-5* on homeostasis.

Analogously, the mutations in the C-terminus of *AtMYB4, AtMYB32,* and *AtMYB7* abolished their transcriptional repressive activity (Zhou et al., 2015). Similarly, truncation of C-terminus in Lilium *virescens* gene (*LhMYB12*) enhanced fruit exocarp color in oil palm (Singh et al., 2014). The relieving of transcription repression by *Myb117* truncation is also evident by the upregulation of transcripts of secondary metabolism in *tf-5*, which are actively repressed by R2R3-MYB TFs (Ma and Constabel, 2019).

### Upregulation of carotene isomerases seems to be the driver for increased carotenoids

Notwithstanding, the complexities underlying the regulation of carotenoids biosynthesis in tomato fruits, the expression of carotenogenic genes involved in biosynthesis, degradation, and esterification is the key determinant of the carotenoids level. Combined proteome and transcriptome analysis of *tf-5* highlighted that the MEP pathway and plastoquinone (PQ) biosynthesis too are modulated and likely contribute to enhanced carotenoids. The activity of PDS protein is extensively determined by a constant supply of PQ, which is an essential cofactor for carotenoid desaturation carried out by phytoene desaturase (PDS) (Rosso et al., 2009). Considering that transcripts contributing to the first step of plastoquinone biosynthesis viz. *tyrosine aminotransferase* are upregulated that may increase plastoquinone supply to PDS. The likely increased plastoquinone supply may offset the effect if any due to a reduction in *plastid terminal oxidase* (PTOX) transcript.

The *tf-5* also upregulated the expression of several genes/proteins contributing to the MEP pathway. In tomato, even before the onset of ripening, the MEP pathway primes geranylgeranyl diphosphate (GGPP) supply to later ripening-triggered carotenoid biosynthesis (Botella□Pavía et al., 2004). Therefore, the upregulation of (E)-4-Hydroxy-3-methyl-but-2-enyl pyrophosphate synthase and farnesyl diphosphate synthase proteins at MG may give a head start to carotenoid synthesis in *tf-5* fruit during ripening. Similarly, increased dimethylallyl-diphosphate:NAD(P) oxidoreductase and isopentenyl- diphosphate:NAD(P) oxidoreductase at BR and isopentenyl- diphosphate Δ-isomerase (IDI2) proteins at RR can boost the supply of GGPP. At the pinnacle of the MEP pathway, the *GGPPS2*, a major GGPP contributor for fruit-specific carotenoid synthesis (Barja et al., 2021), is upregulated in *tf-5* at all ripening stages. The modulation of the MEP pathway at multiple levels in *tf-5* seemingly enhances precursor supply to carotenoids biosynthesis.

Since *tf-5* had substantially higher levels of lycopene, and intermediates such as phytoene and phytofluene, it indicated a strong upregulation of carotenoids biosynthesis. In tomato, phytoene synthase and lycopene β-cyclase (CYCB) are two key regulatory enzymes for fruit-specific carotenoids synthesis. The transcriptome and qRT-PCR did not reveal any significant upregulation of genes modulating the conversion of GGPP to lycopene. Seemingly, higher carotenoids levels in *tf-5* fruit were most likely executed by increased protein levels of CRTISO and ZISO. These two enzymes are crucial for the isomerization of carotene precursors, leading to lycopene formation (Isaacson et al., 2002; Chen et al., 2010). While in leaf, the absence of these enzymes can be compensated by sunlight-mediated isomerization, in fruit the light is not effective, as it fails to penetrate to deeper layers. Consequently, tomato *CRTISO* (*tangerine*) mutant accumulates lycopene only at the fruit surface, where light intensity is sufficient for isomerization (Isaacson et al., 2002).

An increase in carotenoids in *tf-5* may also be causally related to an increase in JA, as JA-deficient tomato mutants show a reduction in lycopene level (Liu et al., 2012). Alike, reduced ABA may enhance carotenoids as ABA deficiency promotes carotenoids levels (Galpaz et al; 2008; Sun et al., 2012). Tomato *hp* fruits bearing high carotenoid levels also have increased carotenoid sequestration proteins (Kilambi et al., 2013). Since in *tf-5* fruit PAP3 protein is high, increased sequestration may contribute to enhanced carotenoids. The increased remorin1 protein in *tf-5* too may contribute to increased carotenoids, as transgenic overexpression of *remorin1* upregulates lycopene in tomato (Cai et al., 2018). However, unlike *remorin1* overexpressing fruits, in *tf-5* the expression *CNR, RIN, and NOR* transcripts was suppressed (**Figure S15**). In sum, ZISO and CRITSO proteins, in concert with phytohormones JA and ABA, PAP3, and remorin1 proteins seem to be the main contributors to increased carotenoid levels in *tf-5*.

### C1 metabolism proteins are upregulated in *tf-5*

The C1 metabolism mediated by folate contributes to nucleotide and amino acid biosynthesis and thus higher amino acids in *tf-5* may arise from upregulation of C1 transfers. Expectedly in *tf-5,* several proteins and transcripts contributing to the folate cycle and methionine cycle were upregulated. Among the four detected folate vitamers, 5-CHO-THF acts as an inhibitor of SHMT (Goyer et al., 2005). Moreover, the 5-CHO-THF level is doubled in *tf-5*, similar to Arabidopsis 5-CHO-THF cycloligase mutant. Therefore, it is plausible that increased 5-CHO-THF, by inhibiting SHMT catalytic activity, may counteract C1 transfer stimulation by increased SHMT protein. Nonetheless, such possibility was discounted in Arabidopsis, where metabolic resilience annulled the influence of increased 5-CHO-THF (Goyer et al., 2005). The inhibitory influence of 5-CHO-THF is not limited to SHMT, recent evidence indicated that it binds to a large number of proteins and can modulate C/N metabolism (Li et al., 2021). Several 5-CHO-THF binding proteins such as sucrose synthase, glutamine synthase are upregulated in the *tf-5* proteome. Since sucrose and glutamine levels are high in *tf-5,* the inhibition of sucrose and glutamine synthase activity by 5-CHO-THF, if any, is minor. Nonetheless, it remains possible that increased 5-CHO-THF may influence the overall metabolome by inhibiting other enzymes.

### Higher folate results from reduced GGH activity

Folates are essential for central metabolism, therefore; mutations in the folate biosynthesis pathway are rare and cause developmental abnormalities. As *tf-5* encodes a *MYB* TF, it is reasonable to expect that increase in folate results from the upregulation of its biosynthesis. Conforming to this expectation, transcripts of the first two enzymes of the folate biosynthesis *GCHI* and *ADCS* along with few other downstream genes were upregulated in *tf-5*. Likewise, in *tf-2* also the folate biosynthesis genes are upregulated. Congruent with upregulation of *ADCS, tf-5* fruits have a higher *p*ABA level, which is akin to higher *p*ABA levels in *ADCS* overexpressing transgenic tomato fruits (Garza et al., 2007) and also in high folate pak choi accessions (Shohag et al. 2020). Our inability to detect a similar increase in pterin may be related to its efficient conversion to downstream intermediates. Even in transgenic tomato fruits overexpressing *GCHI*, the relative accumulation of pterin is far lower than *p*ABA (de la Garza et al., 2004, 2007).

The *in vivo* levels of folate are determined by a combination of synthesis, degradation, and polyglutamylation. In plants, a salvage pathway for folate degradation products regenerates pterin and *p*ABA. This pathway is mediated by GGHs, *p*ABA-glucose hydrolase, pterin aldehyde reductase (PTAR). Since the pterin-6-carboxylate level, an intermediate of salvage pathway in *tf-5* is reduced to half, though *tf-5* has 4-fold higher folate, points to reduced folate degradation. Pterin-6-carboxylate is considered a dead-end product; therefore, its reduced level indicates more efficient folate protection (Orsomondo et al., 2006). Considering that 6-hydroxymethylpterin while detected in WT (MG, BR) is not detected in *tf-5*, the pterin salvage, if any mediated by PTAR, likely takes place by conversion of dihydropterin-6-aldehyde to 6-hydroxymethyldihydropterin (Gorelova et al., 2017a).

In potato cultivars, high *GGH1* expression coincides with high folate levels (Robinson et al., 2019), contrariwise *tf-5* has lower *GGH1* and *GGH2* expressions and reduced GGH enzyme activity. In potato, it was surmised that high *GGH1* expression is either a check valve to limit folate levels to a threshold or it recycles the pABA-glutamate for the fresh synthesis of folate (Robinson et al., 2019). The reduced *GGH1* and *GGH2* transcripts and GGH activity in *tf-2* and *tf-5* favors reduced folate degradation and efficient salvage, which likely elevates folate level. The elevation of folate in tomato *GGH1*, and *GGH2* gene-knockout lines, conforms to the above possibility. Since silencing of both *GGH1* and *GGH2* in Arabidopsis caused a ca. 30% increase in leaf folate (Akhtar et al., 2010), lower GGH activity likely boosts folate level in *tf-5*.

Previous attempts to enhance folate levels in crop plants have focused on overexpressing folate biosynthesis enzymes. Our study highlights a likely dual action of the *tf-5* allele, wherein enhanced expression of folate biosynthesis genes and reduced folate degradation by GGH boost folate levels in fruits. Being a TF, the *tf-5* allele influences a wide range of the responses including γ-aminobutyric acid and carotenoids. The mechanism by which the *tf-5* allele upregulates three different metabolic pathways, amino acids, carotenoids, and folate remains to be deciphered. In general, the mutation in TFs and associated proteins has a broad-spectrum influence on cellular response. Similarly, in tomato mutations in the *high pigment* genes stimulate diverse pathways, such as higher vitamins, flavonoids, and carotenoids levels in fruits (Levin et al., 2006). To sum, our study delineates a novel role for *MYB117* in regulating three important nutraceuticals, γ-aminobutyric acid, folate, and carotenoids in plants. The introgression of the above *MYB117* allele can be used for non-transgenic biofortification of tomato cultivars for improved levels of the above nutraceuticals.

## Experimental Procedures

### Plant materials

The following tomato wild types; Arka Vikas (AV), M82, Rheinland Ruhm (RH), Condine Red (CR) and trifoliate alleles; *tf-1* (LA0512, cv. unknown); *tf-2* (LA0579, cv. CR); *tf-3* (cv. RH); *tf-4*, *tf-z* (cv. M82); and *tf-5* (cv. AV) were used (**Table S1**). The *tf-5* mutant was isolated from the EMS-mutagenized AV M_2_ population (Sharma et al., 2021) and was used in BC_2_F_2_ generation and its progeny. Based on similarity to *trifoliate* mutants (Naz et al., 2013), the *MYB117* gene (Solyc05g007870) of *tf-5* was sequenced and designated as a new *tf-5* allele. The CEL-I assay was used for ascertaining homozygosity of *tf-5* mutation in backcrossed progeny (**Figure S2**, **Table S6**). The gene-edited plants for *GGH1* and *GGH2* genes were raised using protocols described in Kilambi et al. (2021) (**Figure S16**). The tomato plants were grown in the greenhouse as described in Bodanapu et al. (2016). The Arabidopsis wild type (CS_70000) and *lof1* (Salk_025235) and *lof2* (Salk_064076) mutant seedlings were grown at 22°C for 21 days under long day (16h light/8h dark). The leaves were harvested from the 5^th^ node of 6-7 week old plants. The fruits at mature-green (MG), breaker (BR), and red-ripe (RR) stages were harvested from the first and second truss of plants.

### Biochemical Characterizations

The protocols used for different analyses were as follows: Gupta et al. (2014) for °Brix, fruit firmness, and qRT-PCR (**Table S7-S8**); Gupta et al. (2015) for carotenoids; Kilambi et al. (2013) for ethylene emission; Tyagi et al. (2015) for folate; Blancquaert et al. (2013) for pABA; Martín-Tornero et al. (2016) for pterins, and Bodanapu et al. (2016) for phytohormones and metabolites. Only metabolites with a ≥1.5-fold change (Log2 ≥ ± 0.584) and p-value ≤ 0.05 were mapped on the metabolic pathway.

### RNA-Seq

Total RNA from RR fruits was isolated using RNeasy Plant-Mini-Kit as per the manufacturer’s protocol. Isolated RNA was treated with DNase using TURBO DNA-free™ Kit (Ambion, USA). The quality check of RNA was performed by Bioanalyzer 2100, before library construction and three independent biological replicates each of AV and *tf-5* were sequenced. The transcriptome was analyzed by paired-end sequencing using the BGI Seq-500 platform (Shenzhen, China) generating about 11.02 GB bases per sample. The sequencing reads with low-quality, adaptor-polluted, and high content of unknown base (N) reads were removed before analyses using SOAPnuke (v1.5.2) (Chen et al., 2017: Zhu et al., 2018) (**Table S9**). The clean reads were then aligned to the tomato reference genome (ITAG4.0) using HISAT2 (Kim et al., 2015). On average, 97.54% reads mapped, and the uniformity of the mapping result for each sample suggests that the samples are comparable (**Table S10**). Alignment files generated were then used for assigning genomic features using feature counts (Liao et al., 2014) along with the reference annotations. The read counts generated using feature counts were taken as input for identifying differentially expressed genes (DEGs) using DeSeq2 (Love et al., 2014) based on ≥ 1.5-fold change (Log2 ≥ ± 0.58) and p-value ≤ 0.05.

### Proteome profiling

Proteins (70 μg) were separated and identified as described in (Kilambi et al., 2016). *Solanum lycopersicum* iTAG4.1 proteome sequence (ftp://ftp.solgenomics.net/tomato_genome/annotation/ITAG4.1_release/ITAG4.1_proteins.fasta downloaded on August 28, 2020, 34430 sequences, and 11584472 residues) was used as the database against which the searches were done. The functional annotation and classification of the ITAG4.1 proteins was generated using the Mercator 4 tool (https://www.plabipd.de/portal/mercator4) on October 29, 2020 (http://www.plabipd.de/projects/Mercator4_output/GFA-62a085d6ce7e994eb0d871c612e7f528.zip) with the job id GFA-62a085d6ce7e994eb0d871c612e7f528. The mass spectrometry proteomics data are available via ProteomeXchange (Vizcaíno et al., 2014) with the dataset identifier PXD027940. The proteins with ≥ 1.5-fold change (Log2 ≥ ± 0.58) and p-value ≤ 0.05 were considered as differentially expressed. For proteins detected only in *tf-5* or in AV, the missing values were replaced with 1/10 of the lowest value in the dataset.

### PCA and metabolite pathways

Principal Component Analysis (PCA) was performed using Metaboanalyst 4.0 (http://www.metaboanalyst.ca/) after the One-way ANOVA plot (P ≤ 0.05) identified 71 of the 77 metabolites which have a significant contribution. The differentially expressed proteins and transcripts were mapped to the general metabolic pathways using LycoCyc (http://solcyc.solgenomics.net/) and MetGenMap (http://bioinfo.bti.cornell.edu/cgi-bin/MetGenMAP/home.cgi) to highlight patterns of change in *tf-5* compared to AV.

### GGH enzyme assay

GGH enzyme assay was modified from the Orsomando et al. (2005). The fruit tissue was homogenized in liquid N_2_ to a fine powder and to the homogenate 100 mM potassium phosphate, pH 6.0, 10% (v/v) glycerol, 10 mM β-mercaptoethanol, and 3% (w/v) polyvinylpolypyrrolidone was added. After centrifugation (20,000*g*, 20 min, 4°C) the supernatant was desalted using a 10 kDa cut-off membrane filter (Pall Corporation, USA). The GGH assay mixture contained 50 μM *p*ABAGlu7 substrate, 100 mM potassium phosphate, pH 6.0, 10% (v/v) glycerol, 10 mM β-mercaptoethanol, and supernatant (30 μg total protein) in a final volume of 100 μL. After 4 hr incubation at 37°C, the reaction mixture was boiled for 3 min, followed by centrifugation (16,000*g*, 10 min). The appearance of the product pABAGlu was measured using Orbitrap-Exactive LC-MS in a reversed-phase Luna C18 column (5 μm particle size, 250 mm x 4.60 mm ID) (Phenomenex, USA) using gradient elution program. The gradient comprised of a binary solvent system consisting of 0.1% (v/v) formic acid in water (solvent A) and 0.1% (v/v) formic acid in acetonitrile (solvent B) at a flow rate of 0.7 mL per minute. The injection volume was 7.5 μL and the run time was 17 min. The retention time of *p*ABAGlu was 6.8 min and m/z 265.08. Enzyme activity was measured using the *p*ABAGlu as an external standard and expressed as nmol min^-1^ mg^-1^ protein. Other parameters of Orbitrap-Exactive LC-MS were the same as described by Tyagi et al. (2015).

### Promoter analysis

The promoter sequences (2□Kb) of analyzed genes (**Table S11)** were retrieved from the SOL genomics network (https://solgenomics.net/) and analyzed for MYB TFs binding sites using PlantCARE (http://bioinformatics.psb.ugent.be/webtools/plantcare/html/).

### Statistical analysis

A minimum of three independent biological replicates was used for all experiments. Statistical Analysis On Microsoft Excel (https://prime.psc.riken.jp/MetabolomicsSoftware/StatisticalAnalysisOn-MicrosoftExcel) was used to obtain significant differences between data points. A Student’s *t*-test was also performed to determine significant differences (* for p-value ≤ 0.05). Heat maps and 3D-PCA plots were generated using Morpheus (https://software.broadinstitute.org/morpheus/) and MetaboAnalyst 4.0 (https://www.metaboanalyst.ca/), respectively.

## Acknowledgments

This work was supported by the Department of Biotechnology (DBT), India grants, BT/PR11671/PBD/16/828/2008, BT/PR/7002/PBD/16/1009/2012, and BT/COE/34/SP15209/2015 to RS and YS. We also acknowledge fellowships by CSIR to KT, UGC to AS, DST-N-PDF to NT, and DBT-RA to SS.

## Author Contributions

KT, YS, and RS designed this project and wrote the manuscript. KT. performed most of the experiments AS did proteome profiling, MR, AY and SSi did transcriptome profiling, SSa and NT raised genome edited GGH lines.

## Supporting Information

**Figure S1.** *Trifoliate* mutant stabilization and backcrossing

**Figure S2.** Mutation detection in *tf-5* by Sanger Sequencing, and CEL- I endonuclease assay

**Figure S3**. The phenotype of ripe *tf-1* fruit.

**Figure S4. C**hanges in °Brix, fruit firmness, and seed weight in *tf* alleles.

**Figure S5**. Total folate in the leaf of Arabidopsis *lof* mutants, *tf* alleles, and carotenoids in *tf-5* leaves

**Figure S6**. Dendrogram showing maximum likelihood phylogeny of selected R2R3-MYBs.

**Figure S7**. Proteome profiling of *tf-5* and its wild type.

**Figure S8**. Transcriptome profiling of *tf-5* and its wild type

**Figure S9.** Mapman representations of proteome and transcriptome changes in *tf-5* fruits

**Figure S10**. LycoCyc representations of proteome and transcriptome changes in *tf-5* fruits

**Figure S11**. Folate biosynthesis pathway, *MYB 117* expression, and GGH activity in *tf-2*.

**Figure S12**. MYB binding domains in the promoter of carotenoids and folate biosynthesis genes.

**Figure S13**. Percent polyglutamylation in red ripe fruits of Arka Vikas and *tf-5*

**Figure S14.** Total folate in red ripe fruits of T0 *GGH1* and *GGH2* gene-edited lines.

**Figure S15**. Expression of ripening regulatory genes in *tf-5* and its wild type.

**Figure S16.** Generation and confirmation of genome-edited *GGH1* and *GGH2* plants.

**Table S1.** List of different *tf* alleles, their cultivars, and the site of mutations.

**Table S2.** MYB proteins of *tf* alleles showing structural diversity compared to wild-type MYB117.

**Table S3.** Morphological characterization of AV and *tf-5*.

**Table S4.** Carotenoids profiles of *tf* alleles and their respective WTs.

**Table S5.** Folate, *p*ABA, p6C, and HMPt content in *tf* alleles and their respective WTs.

**Table S6.** Primers set used for mutation detection in *tf-5* using CEL-I assay.

**Table S7.** Primers used for qRT-PCR of folate metabolism and *MYB117* gene.

**Table S8.** Primers used for qRT-PCR of carotenoid biosynthesis and ripening regulators genes.

**Table S9.** The filtered read statistics of AV and *tf-5* transcriptome samples.

**Table S10.** Summary of genome mapping statistics using HISAT2 in AV and *tf-5*.

**Table S11.** Genes name and their promoter sequence coordinates used for MYB binding site.

**Dataset S1.** Metabolites profile of *tf-5*, its WT and qRT-PCR data during fruit ripening.

**Dataset S2.** Proteome profile of *tf-5* and its WT during fruit ripening.

**Dataset S3.** Differentially expressed proteins of *tf-5* mentioned in text/Figures

**Dataset S4.** Differentially expressed transcripts of *tf-5* and its WT at red ripe stage.

## References

Akhtar TA, Orsomando G, Mehrshahi P, Lara□Núñez A, Bennett MJ, Gregory III JF, Hanson AD. (2010) A central role for gamma□glutamyl hydrolases in plant folate homeostasis. The Plant Journal. 64: 256–66.

Azari R, Tadmor Y, Meir A, Reuveni M, Evenor D, Nahon S, Shlomo H, Chen L, Levin I (2010) Light signaling genes and their manipulation towards modulation of phytonutrient content in tomato fruits. Biotechnol Adv 28: 108–118

Barja MV, Ezquerro M, Beretta S, Diretto G, Florez□Sarasa I, Feixes E, Fiore A, Karlova R, Fernie AR, Beekwilder J, Rodríguez□Concepción M. (2021) Several geranylgeranyl diphosphate synthase isoforms supply metabolic substrates for carotenoid biosynthesis in tomato. New Phytologist. https://doi.org/10.1111/nph.17283

Blancquaert D, Van Daele J, Storozhenko S, Stove C, Lambert W, Van Der Straeten D. (2013) Rice folate enhancement through metabolic engineering has an impact on rice seed metabolism, but does not affect the expression of the endogenous folate biosynthesis genes. Plant Molecular Biology. 83: 329–49.

Bodanapu R, Gupta SK, Basha PO, Sakthivel K, Sreelakshmi Y, Sharma R. (2016) Nitric oxide overproduction in tomato shr mutant shifts metabolic profiles and suppresses fruit growth and ripening. Frontiers in Plant Science. 7: 1714.

Botella□Pavía P, Besumbes Ó, Phillips MA, Carretero□Paulet L, Boronat A, Rodríguez□Concepción M. (2004) Regulation of carotenoid biosynthesis in plants: evidence for a key role of hydroxymethylbutenyl diphosphate reductase in controlling the supply of plastidial isoprenoid precursors. The Plant Journal. 40: 188–99

Bramley PM, Bird CR, Schuch W. (1993) Carotenoid biosynthesis and manipulation. In Biosynthesis and manipulation of plant products. Ed. Grierson D., Springer, Dordrecht. pp. 139–177

Cai J, Qin G, Chen T, Tian S. (2018) The mode of action of remorin1 in regulating fruit ripening at transcriptional and post□transcriptional levels. New Phytologist. 219: 1406–20.

Cao H, Chen J, Yue M, Xu C, Jian W, Liu Y, Song B, Gao Y, Cheng Y, Li Z. (2020) Tomato transcriptional repressor MYB70 directly regulates ethylene-dependent fruit ripening. The Plant Journal. 104: 1568–81.

Chandler PM, Harding CA. (2013) ‘Overgrowth’ mutants in barley and wheat: new alleles and phenotypes of the ‘Green Revolution’ DELLA gene. Journal of Experimental Botany. 64: 1603–13

Chaudhary P, Sharma A, Singh B, Nagpal AK. (2018) Bioactivities of phytochemicals present in tomato. Journal of Food Science and Technology. 55: 2833–49

Chen Y, Li F, Wurtzel ET. (2010) Isolation and characterization of the Z-ISO gene encoding a missing component of carotenoid biosynthesis in plants. Plant Physiology. 153: 66–79.

Cheng ZJ, Zhao XY, Shao XX, Wang F, Zhou C, Liu YG, Zhang Y, Zhang XS. (2014) Abscisic acid regulates early seed development in Arabidopsis by ABI5-mediated transcription of SHORT HYPOCOTYL UNDER BLUE1. The Plant Cell. 26: 1053–68.

de La Garza RD, Gregory JF, Hanson AD. (2007) Folate biofortification of tomato fruit. Proceedings of the National Academy of Sciences. 104: 4218–22.

de la Garza RD, Quinlivan EP, Klaus SM, Basset GJ, Gregory JF, Hanson AD. (2004) Folate biofortification in tomatoes by engineering the pteridine branch of folate synthesis. Proceedings of the National Academy of Sciences. 101:1 3720–5.

De Lepeleire J, Strobbe S, Verstraete J, Blancquaert D, Ambach L, Visser RG, Stove C, Van Der Straeten D. (2018) Folate biofortification of potato by tuber-specific expression of four folate biosynthesis genes. Molecular Plant. 11: 175–88.

Fernandez-Moreno JP, Tzfadia O, Forment J, Presa S, Rogachev I, Meir S, Orzaez D, Aharoni A, Granell A. (2016) Characterization of a new pink-fruited tomato mutant results in the identification of a null allele of the SlMYB12 transcription factor. Plant Physiology. 171: 1821–36.

Fujisawa M, Nakano T, Shima Y, Ito Y. (2013) A large-scale identification of direct targets of the tomato MADS box transcription factor RIPENING INHIBITOR reveals the regulation of fruit ripening. The Plant Cell. 25: 371–86.

Fujisawa M, Shima Y, Higuchi N, Nakano T, Koyama Y, Kasumi T, Ito Y. (2012) Direct targets of the tomato-ripening regulator RIN identified by transcriptome and chromatin immunoprecipitation analyses. Planta. 235: 1107–22.

Galpaz N, Wang Q, Menda N, Zamir D, Hirschberg J. (2008) Abscisic acid deficiency in the tomato mutant high□pigment 3 leading to increased plastid number and higher fruit lycopene content. The Plant Journal. 53: 717–30.

Gorelova V, Ambach L, Rébeillé F, Stove C, Van Der Straeten D. (2017a) Folates in plants: research advances and progress in crop biofortification. Frontiers in Chemistry. 5: 21.

Gorelova V, De Lepeleire J, Van Daele J, Pluim D, Meï C, Cuypers A, Leroux O, Rébeillé F, Schellens JH, Blancquaert D, Stove CP, Van Der Straeten D. (2017b) Dihydrofolate reductase/thymidylate synthase fine-tunes the folate status and controls redox homeostasis in plants. The Plant Cell. 29: 2831–53.

Goyer A, Collakova E, de la Garza RD, Quinlivan EP, Williamson J, Gregory JF, Shachar-Hill Y, Hanson AD. (2005) 5-Formyltetrahydrofolate is an inhibitory but well tolerated metabolite in Arabidopsis leaves. Journal of Biological Chemistry. 280: 26137–42.

Grierson D (2013) Ethylene and the Control of Fruit Ripening. In The Molecular Biology and Biochemistry of Fruit Ripening. Eds. Seymour GB, Tucker GA, Poole M, Giovannoni, J. Wiley-Blackwell Chichester. pp 43–74.

Guo W, Lian T, Wang B, Guan J, Yuan D, Wang H, Safiul Azam FM, Wan X, Wang W, Liang Q, Wang H. (2019) Genetic mapping of folate QTLs using a segregated population in maize. Journal of Integrative Plant Biology. 61: 675–90.

Gupta P, Reddaiah B, Salava H, Upadhyaya P, Tyagi K, Sarma S, Datta S, Malhotra B, Thomas S, Sunkum A, Devulapalli S., Till BJ, Sreelakshmi Y, Sharma R (2017) NGS-based identification of induced mutations in a doubly mutagenized tomato (*Solanum lycopersicum*) population. The Plant Journal 92: 495–508

Gupta SK, Sharma S, Santisree P, Kilambi HV, Appenroth K, Sreelakshmi Y, Sharma R. (2014) Complex and shifting interactions of phytochromes regulate fruit development in tomato. Plant, Cell and Environment. 37: 1688–702.

Hanson AD, Gregory III JF. (2011) Folate biosynthesis, turnover, and transport in plants. Annual Review of Plant Biology.62: 105–25.

Houben M, Van de Poel B. (2019) 1-Aminocyclopropane-1-carboxylic acid oxidase (ACO): the enzyme that makes the plant hormone ethylene. Frontiers in Plant Science. 10: 695.

Huang GQ, Gong SY, Xu WL, Li W, Li P, Zhang CJ, Li DD, Zheng Y, Li FG, Li XB. (2013) A fasciclin-like arabinogalactan protein, GhFLA1, is involved in fiber initiation and elongation of cotton. Plant Physiology. 161: 1278–90.

Isaacson T, Ronen G, Zamir D, Hirschberg J. (2002) Cloning of tangerine from tomato reveals a carotenoid isomerase essential for the production of β-carotene and xanthophylls in plants. The Plant Cell 14; 333–42.

Karlova R, Rosin FM, Busscher-Lange J, Parapunova V, Do PT, Fernie AR, Fraser PD, Baxter C, Angenent GC, de Maagd RA. (2011) Transcriptome and metabolite profiling show that APETALA2a is a major regulator of tomato fruit ripening. The Plant Cell. 23: 923–41.

Kilambi HV, Dindu A, Sharma K, Nizampatnam NR, Gupta N, Thazath NP, Dhanya AJ, Tyagi K, Sharma S, Sharma R, Sreelakshmi Y. (2021) The new kid on the block: A dominant□negative mutation of phototropin1 enhances carotenoid content in tomato fruits. The Plant Journal. 106: 844–61.

Kilambi HV, Kumar R, Sharma R, Sreelakshmi Y. (2013) Chromoplast-Specific carotenoid-associated protein appears to be important for enhanced accumulation of carotenoids in hp1 tomato fruits. Plant Physiology 161: 2085–2101

Kilambi HV, Manda K, Rai A, Charakana C, Bagri J, Sharma R, Sreelakshmi Y (2017) Green-fruited *Solanum habrochaites* lacks fruit-specific carotenogenesis due to metabolic and structural blocks. Journal of Experimental Botany 68: 4803–4819.

Kilambi HV, Manda K, Sanivarapu H, Maurya VK, Sharma R, Sreelakshmi Y. (2016) Shotgun proteomics of tomato fruits: evaluation, optimization and validation of sample preparation methods and mass spectrometric parameters. Frontiers in Plant Science. 7: 969.

Kim D, Langmead B, Salzberg SL. (2015) HISAT: a fast spliced aligner with low memory requirements. Nature Methods. 12: 357–60.

Kim J, Lee JG, Hong Y, Lee EJ. (2019) Analysis of eight phytohormone concentrations, expression levels of ABA biosynthesis genes, and ripening-related transcription factors during fruit development in strawberry. Journal of Plant Physiology. 239: 52–60.

Lee DK, Geisler M, Springer PS. (2009) LATERAL ORGAN FUSION1 and LATERAL ORGAN FUSION2 function in lateral organ separation and axillary meristem formation in Arabidopsis. Development. 136: 2423–32.

Levin I, Frankel P, Gilboa N, Tanny S, Lalazar A. (2003) The tomato dark green mutation is a novel allele of the tomato homolog of the DEETIOLATED1 gene. Theoretical and Applied Genetics. 106: 454–60.

Levin I, De Vos CR, Tadmor Y, Bovy A, Lieberman M, Oren-Shamir M, Segev O, Kolotilin I, Keller M, Ovadia R, Meir A. (2006) High pigment tomato mutants— more than just lycopene (a review). Israel Journal of Plant Sciences. 54: 179–90.

Li N, Xu R, Li Y. (2019) Molecular networks of seed size control in plants. Annual Review of Plant Biology. 70: 435–63.

Li W, Liang Q, Mishra RC, Sanchez-Muñoz R, Wang H, Chen X, Van Der Straeten D, Zhang C, Xiao Y. (2021) The 5-formyl-tetrahydrofolate proteome links folates with C/N metabolism and reveals feedback regulation of folate biosynthesis. The Plant Cell. https://doi.org/10.1093/plcell/koab19

Li Z, Peng R, Tian Y, Han H, Xu J, Yao Q. (2016) Genome-wide identification and analysis of the MYB transcription factor superfamily in Solanum lycopersicum. Plant and Cell Physiology. 57: 1657–77.

Liang Q, Wang K, Liu X, Riaz B, Jiang L, Wan X, Ye X, Zhang C. (2019) Improved folate accumulation in genetically modified maize and wheat. Journal of Experimental Botany. 70: 1539–51.

Liao Y, Smyth GK, Shi W. (2014) FeatureCounts: an efficient general purpose program for assigning sequence reads to genomic features. Bioinformatics. 30: 923–30.

Lin Z, Hong Y, Yin M, Li C, Zhang K, Grierson D. (2008) A tomato HD□Zip homeobox protein, LeHB□1, plays an important role in floral organogenesis and ripening. The Plant Journal. 55: 301–10.

Liu L, Shao Z, Zhang M, Wang Q. (2015) Regulation of carotenoid metabolism in tomato. Molecular Plant. 8: 28–39

Liu L, Wei J, Zhang M, Zhang L, Li C, Wang Q. (2012) Ethylene independent induction of lycopene biosynthesis in tomato fruits by jasmonates. Journal of Experimental Botany. 63: 5751–61.

Llorente B, D’Andrea L, Ruiz□Sola MA, Botterweg E, Pulido P, Andilla J, Loza□Alvarez P, Rodriguez□Concepcion M. (2016) Tomato fruit carotenoid biosynthesis is adjusted to actual ripening progression by a light□dependent mechanism. The Plant Journal. 85: 107–19.

Love MI, Huber W, Anders S. (2014) Moderated estimation of fold change and dispersion for RNA-seq data with DESeq2. Genome Biology. 15: 1–21.

Ma D, Constabel CP. (2019) MYB repressors as regulators of phenylpropanoid metabolism in plants. Trends in Plant Science. 24: 275–89.

Martinez CC, Li S, Woodhouse MR, Sugimoto K, Sinha NR. (2021) Spatial transcriptional signatures define margin morphogenesis along the proximal-distal and medio-lateral axes in tomato (Solanum lycopersicum) leaves. The Plant Cell. 33: 44–65.

Martín-Tornero E, Gómez DG, Durán-Merás I, Espinosa-Mansilla A. (2016) Development of an HPLC-MS method for the determination of natural pteridines in tomato samples. Analytical Methods. 8: 6404–14.

Millard PS, Kragelund BB, Burow M. (2019) R2R3 MYB transcription factors–Functions outside the DNA-binding domain. Trends in Plant Science. 24: 934–46.

Naz AA, Raman S, Martinez CC, Sinha NR, Schmitz G, Theres K. (2013) Trifoliate encodes an MYB transcription factor that modulates leaf and shoot architecture in tomato. Proceedings of the National Academy of Sciences. 110: 2401–6.

Neuman H, Galpaz N, Cunningham Jr FX, Zamir D, Hirschberg J. (2014) The tomato mutation nxd1 reveals a gene necessary for neoxanthin biosynthesis and demonstrates that violaxanthin is a sufficient precursor for abscisic acid biosynthesis. The Plant Journal. 78: 80–93.

Orsomando G, Bozzo GG, de la Garza RD, Basset GJ, Quinlivan EP, Naponelli V, Rébeillé F, Ravanel S, Gregory III JF, Hanson AD. (2006) Evidence for folate□salvage reactions in plants. The Plant Journal. 46: 426–35.

Osorio S, Alba R, Damasceno CM, Lopez-Casado G, Lohse M, Zanor MI, Tohge T, Usadel B, Rose JK, Fei Z, Giovannoni JJ. (2011) Systems biology of tomato fruit development: combined transcript, protein, and metabolite analysis of tomato transcription factor (nor, rin) and ethylene receptor (Nr) mutants reveals novel regulatory interactions. Plant Physiology. 157: 405–25.

Pireyre M, Burow M. (2015) Regulation of MYB and bHLH transcription factors: a glance at the protein level. Molecular Plant. 8: 378–88.

Robinson BR, Garcia Salinas C, Ramos Parra P, Bamberg J, Diaz de la Garza RI, Goyer A. (2019) Expression levels of the γ-glutamyl hydrolase I gene predict vitamin B9 content in potato tubers. Agronomy. 9: 734.

Rosso D, Bode R, Li W, Krol M, Saccon D, Wang S, Schillaci LA, Rodermel SR, Maxwell DP, Huner NP. (2009) Photosynthetic redox imbalance governs leaf sectoring in the Arabidopsis thaliana variegation mutants immutans, spotty, var1, and var2. The Plant Cell. 21: 3473–92.

Schubert R, Dobritzsch S, Gruber C, Hause G, Athmer B, Schreiber T, Marillonnet S, Okabe Y, Ezura H, Acosta IF, Tarkowska D. (2019) Tomato MYB21 acts in ovules to mediate jasmonate-regulated fertility. The Plant Cell. 31: 1043–62.

Sharma K, Gupta S, Sarma S, Rai M, Sreelakshmi Y, Sharma R. (2021) Mutations in tomato 1□aminocyclopropane carboxylic acid synthase2 uncover its role in development beside fruit ripening. The Plant Journal. 106: 95–112.

Shima Y, Kitagawa M, Fujisawa M, Nakano T, Kato H, Kimbara J, Kasumi T, Ito Y. (2013). Tomato FRUITFULL homologues act in fruit ripening via forming MADS-box transcription factor complexes with RIN. Plant Molecular Biology, 82: 427–438.

Shohag MJ, Wei Y, Zhang J, Feng Y, Rychlik M, He Z, Yang X. (2020) Genetic and physiological regulation of folate in pak choi (Brassica rapa subsp. Chinensis) germplasm. Journal of Experimental Botany. 71: 4914–29.

Singh R, Low ET, Ooi LC, Ong-Abdullah M, Nookiah R, Ting NC, Marjuni M, Chan PL, Ithnin M, Manaf MA, Nagappan J. (2014) The oil palm VIRESCENS gene controls fruit colour and encodes a R2R3-MYB. Nature Communications. 5: 1–8.

Stanley L, Yuan YW. (2019) Transcriptional regulation of carotenoid biosynthesis in plants: so many regulators, so little consensus. Frontiers in Plant Science. 10: 1017

Storozhenko S, De Brouwer V, Volckaert M, Navarrete O, Blancquaert D, Zhang GF, Lambert W, Van Der Straeten D. (2007) Folate fortification of rice by metabolic engineering. Nature Biotechnology. 25: 1277–9.

Sun L, Yuan B, Zhang M, Wang L, Cui M, Wang Q, Leng P. (2012) Fruit-specific RNAi-mediated suppression of SlNCED1 increases both lycopene and β-carotene contents in tomato fruit. Journal of Experimental Botany. 63: 3097–108.

Tyagi K, Upadhyaya P, Sarma S, Tamboli V, Sreelakshmi Y, Sharma R. (2015) High performance liquid chromatography coupled to mass spectrometry for profiling and quantitative analysis of folate monoglutamates in tomato. Food Chemistry. 179: 76–84.

Upadhyaya P, Tyagi K, Sarma S, Tamboli V, Sreelakshmi Y, Sharma R. (2017) Natural variation in folate levels among tomato (Solanum lycopersicum) accessions. Food Chemistry. 217: 610–9.

Vizcaíno JA, Deutsch EW, Wang R, Csordas A, Reisinger F, Rios D, Dianes JA, Sun Z, Farrah T, Bandeira N, Binz PA. (2014) ProteomeXchange provides globally coordinated proteomics data submission and dissemination. Nature Biotechnology. 32: 223–6.

Vrebalov J, Pan IL, Arroyo AJ, McQuinn R, Chung M, Poole M, Rose J, Seymour G, Grandillo S, Giovannoni J, Irish VF. (2009) Fleshy fruit expansion and ripening are regulated by the tomato SHATTERPROOF gene TAGL1. The Plant Cell. 21: 3041–62.

Waller JC, Akhtar TA, Lara-Núñez A, Gregory III JF, McQuinn RP, Giovannoni JJ, Hanson AD. (2010) Developmental and feedforward control of the expression of folate biosynthesis genes in tomato fruit. Molecular Plant. 3: 66–77.

Wei DO, Cheng ZJ, Xu JL, Zheng TQ, Wang XL, Zhang HZ, Jie WA, Wan JM. (2014) Identification of QTLs underlying folate content in milled rice. Journal of Integrative Agriculture. 13: 1827–34

Wu M, Xu X, Hu X, Liu Y, Cao H, Chan H, Gong Z, Yuan Y, Luo Y, Feng B, Li Z. (2020) SlMYB72 regulates the metabolism of chlorophylls, carotenoids, and flavonoids in tomato fruit. Plant Physiology. 183: 854–68.

Zhu F, Luo T, Liu C, Wang Y, Yang H, Yang W, Zheng L, Xiao X, Zhang M, Xu R, Xu J. (2017) An R2R3□MYB transcription factor represses the transformation of α□and β□branch carotenoids by negatively regulating expression of CrBCH2 and CrNCED5 in flavedo of Citrus reticulate. New Phytologist. 216: 178–92.

Zhu FY, Chen MX, Ye NH, Qiao WM, Gao B, Law WK, Tian Y, Zhang D, Zhang D, Liu TY, Hu QJ. (2018) Comparative performance of the BGISEQ-500 and Illumina HiSeq4000 sequencing platforms for transcriptome analysis in plants. Plant Methods. 14: 1–4.

