## Supplementary material for "Seeing the unseen: A trifoliate (MYB117) mutant allele fortifies folate and carotenoids in tomato fruits": Table S1

**Table S1.** List of different *tf* alleles, their cultivars, and the site of mutations.

| Alleles | Site of mutation based on mRNA coordinates | Protein | Comments | Seed source |
|---|---|---|---|---|
| Wild type | mRNA length 1254 (Gene length 1976 nucleotides) | Protein length 417 amino acids | Has two MYB domains MYB I (115-166) and MYB II (167-221) | - |
| *tf-1* (LA0512, cv. unknown) | Gene cannot be amplified ( macro lesion was detected in *tf-1*, where Naz et al., (2013) were unable to amplify genomic fragments including exon 1, exon 2, and part of the promoter region) | No protein | | (www.tgrc.ucdavis.edu) |
| *tf-2* (LA0579, cv. CR) | Frameshift resulting from deletion of T at nucleotide 637 | Tryptophan to Stop at position 213 | MYB I domain intact (115-166) but MYB II domain truncated (115-212) | (www.tgrc.ucdavis.edu) and Prof. Klaus Theres, MPIPZ, Germany |
| *tf-3* (cv. RH) | Splicing defect caused by T to C at nucleotide 963 followed by the deletion of 5 nucleotides | Glycine to Stop at position 321 | Both MYB I, and MYB II domains are intact | IPK Gatersleben, Germany |
| *tf-4* (cv. M82) | G to A at nucleotide 470 | Serine to Asparagine at position 157 (MYB domain) | Mutation in MYB domain I (115-166), and MYB II domain intact | IPK Gatersleben, Germany |
| *tf-z* (e0761B, cv. M82) | G to A at nucleotide 368 | Tryptophan to Stop at position 123 | Both MYB I and MYB II domains are lost | Prof. Dani Zamir, HUJI, Israel |
| *tf-5* (cv. AV) | C to T at nucleotide 1096 (genomic position 1818) | Glutamine to stop codon at 366 | Both MYB I and MYB II domains are intact | In-house isolated |

Table updated from Naz et al., 2013.

**Table S2.** MYB proteins of *tf* alleles showing structural diversity compared to wild-type MYB117.

**A.** Structures of tomato MYB117 and different *tf* alleles generated using ITASSER. The mouse R2R3 domain (1MSE) structure is from Protein data bank https://www.rcsb.org/structure/1MSE.

**B.** The alignment of different *tf* allele's MYB protein with 1MSE and wild type MYB117 protein.

**Table S2 A**.

| Sr no. | Alleles | Protein | Structure | Estimated score from ITASSER |
|---|---|---|---|---|
| 1 | Wild type | Protein length 417 amino acids | 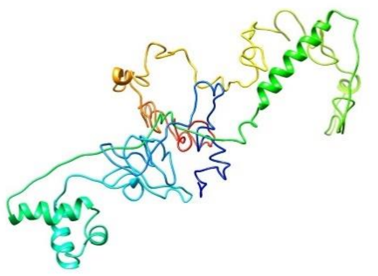 | C-score = -3.69<br>Estimated TM-score = 0.31±0.10<br>Estimated RMSD = 16.2±3.1Å |

| Sr no. | Alleles | Protein | Structure | Estimated score from ITASSER |
|---|---|---|---|---|
| 3 | 1MSE (MYB DNA-BINDING DOMAIN of Mouse) | Protein length 105 amino acid | 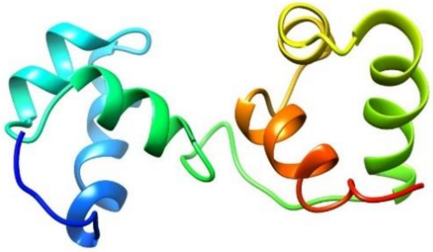 | From Protein data bank https://www.rcsb.org/structure/1MSE |
| 2 | *tf-1* (cv. unknown) | No protein | - | - |
| 3 | *tf-2* (cv. CR) | Tryptophan to Stop at position 213 | 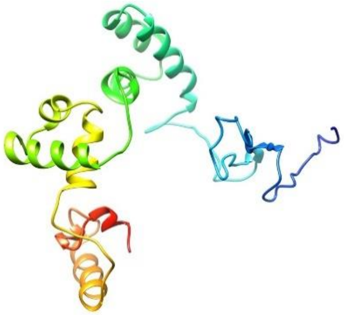 | C-score = -1.34<br>Estimated TM-score = 0.55±0.15<br>Estimated RMSD = 8.4±4.5Å |

| Sr no. | Alleles | Protein | Structure | Estimated score from ITASSER |
|---|---|---|---|---|
| 4 | *tf-3* (cv. RH) | Glycine to Stop at position 321 | 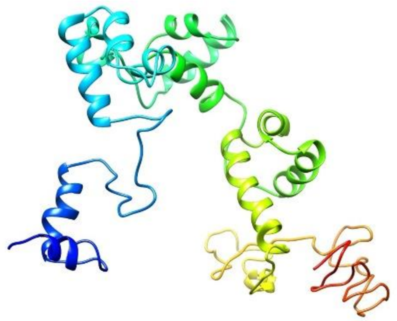 | C-score = -3.16<br>Estimated TM-score = 0.36±0.12<br>Estimated RMSD = 14.0±3.9Å |
| 5 | *tf-4* (cv. M82) | Serine to Asparagine at position 157 (MYB domain) | 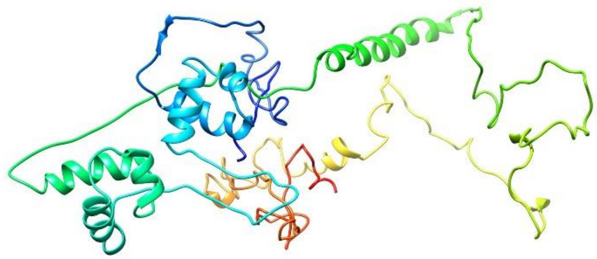 | C-score = -3.66<br>Estimated TM-score = 0.31±0.10<br>Estimated RMSD = 16.1±3.1Å |
| 6 | *tf-z* (cv. M82) | Tryptophan to Stop at position 123 | - | - |

| Sr no. | Alleles | Protein | Structure | Estimated score from ITASSER |
|---|---|---|---|---|
| 7 | *tf-5* (cv. AV) | Glutamine to stop codon at 366 | 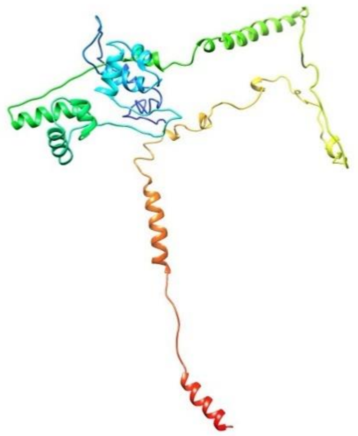 | C-score = -2.53<br>Estimated TM-score = 0.42±0.14<br>Estimated RMSD = 12.7±4.3Å |

The protein structure was generated by homology modelling threading method using ITASSER (https://zhanggroup.org/I-TASSER/) and the best protein structure was selected based on the score.
Score includes 1. C score: confidence score for estimating the quality of a predicted model by ITASSER
2. TM Score and RMSD: TM-score and RMSD are known standards for measuring structural similarity between two structures, which are usually used to measure the accuracy of structure modeling when the native structure is known.
TM Score: the template modeling score or TM-score is a measure of similarity between two protein structures.
RMSD: Root mean square deviation (RMSD) is used for measuring the difference between the backbones of a protein from its initial structural conformation to its final position. The stability of the protein relative to its conformation can be determined by the deviations produced during its simulation.

**Reference for scores and literature:**
https://zhanggroup.org//I-TASSER/example/cscore.txt#:~:text = C%2Dscore%20is%20a%20confidence,of%20the%20structure%20assembly%20simulations.

https://zhanggroup.org/I-TASSER/FAQ.html

**Roy A, Kucukural A, Zhang Y.** (2010) I-TASSER: a unified platform for automated protein structure and function prediction. *Nature Protocols* **5:** 725-738.

**Yang J, Yan R, Roy A, Xu D, Poisson J, Zhang Y.** (2015). The I-TASSER Suite: Protein structure and function prediction. *Nature Methods* **12:** 7-8.

**Yang J, Zhang Y**. (2015) I-TASSER server: new development for protein structure and function predictions. *Nucleic Acids Research* **43:** W174-W181.

**Table S2B**

| Sr no | Alignment | TM align Structure | TM align Score |
|---|---|---|---|
| 1 | *tf2* aligned with 1MSE | 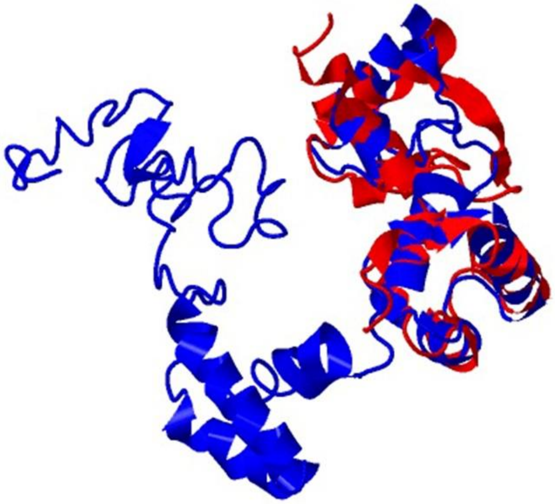 | *tf2* (blue) = 212 residues<br><br>1MSE (red) = 105 residues<br><br>Aligned length = 95<br><br>RMSD = 2.43<br><br>Seq_ID = n_identical/n_aligned = 0.389<br><br>TM-score = 0.38656 (if normalized by the length of *tf-2*)<br><br>TM-score = 0.70302 (if normalized by the length of 1MSE) |

| Sr no | Alignment | TM align Structure | TM align Score |
|---|---|---|---|
| 2 | *tf-2* aligned with wild type MYB117 | 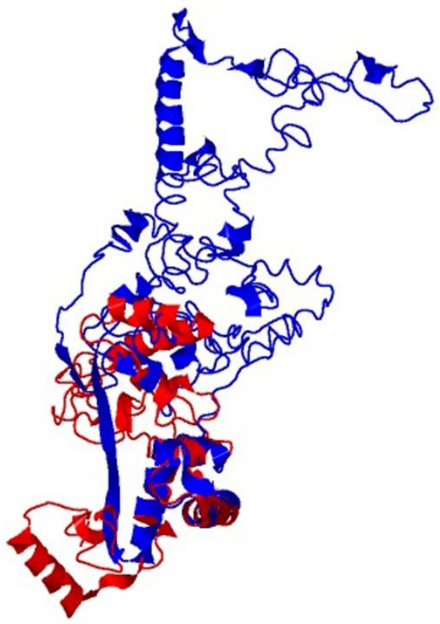 | Wild (blue): 417 residues<br><br>*tf-2* (red): 212 residues<br><br>Aligned length = 109<br><br>RMSD = 3.79<br><br>Seq_ID = n_identical/n_aligned = 0.505<br><br>TM-score = 0.22065 (if normalized by the length of wild type MYB117)<br><br>TM-score = 0.39828 (if normalized by the length of *tf-2*) |

| Sr no | Alignment | TM align Structure | TM align Score |
|---|---|---|---|
| 3 | *tf-2* aligned with *tf-5* | 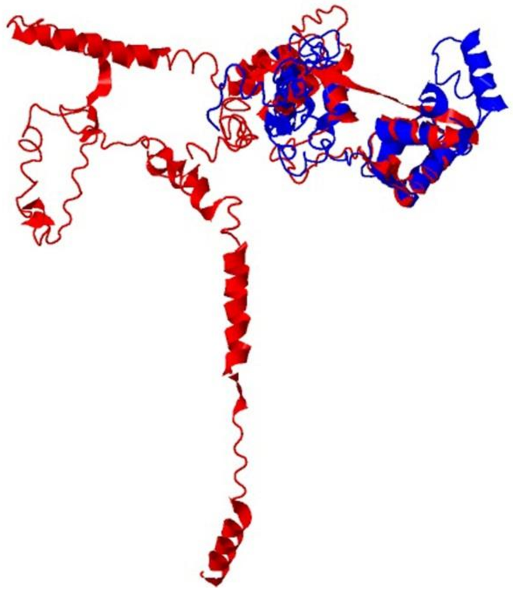 | *tf-2* (blue): 212 residues<br><br>*tf-5* (red): 365 residues<br><br>Aligned length = 130<br><br>RMSD = 3.71<br><br>Seq_ID = n_identical/n_aligned = 0.423<br><br>TM-score = 0.47697 (if normalized by the length of *tf-2*)<br><br>TM-score = 0.29629 (if normalized by the length of *tf-5*) |

| Sr no | Alignment | TM align Structure | TM align Score |
|---|---|---|---|
| 4 | *tf-5* aligned with wild type MYB117 | 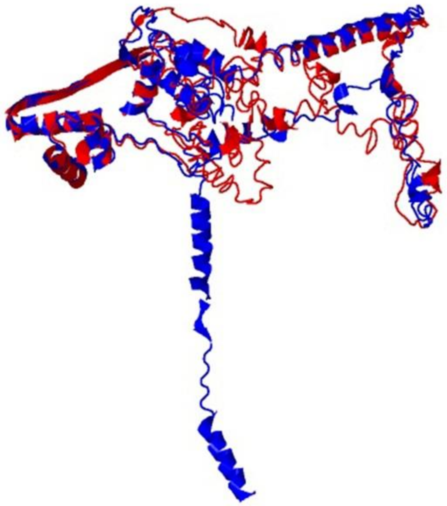 | *tf*-5 (blue): 365 residues<br><br>wild type MYB117 (red): 417 residues<br><br>Aligned length = 280<br><br>RMSD = 4.58<br><br>Seq_ID = n_identical/n_aligned = 0.854<br><br>TM-score = 0.61036 (if normalized by the length of *tf-5*)<br><br>TM-score = 0.54275 (if normalized by the length of wild type MYB117) |

| Sr no | Alignment | TM align Structure | TM align Score |
|---|---|---|---|
| 5 | *tf-5* aligned with 1MSE | 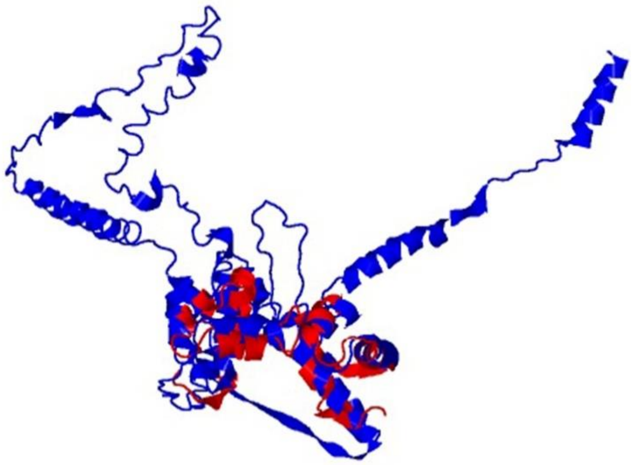 | *tf-5* (blue) = 212 residues<br><br>1MSE (red) = 105 residues<br><br>Aligned length = 102<br><br>RMSD = 2.87<br><br>Seq_ID = n_identical/n_aligned = 0.137<br><br>TM-score = 0.24489 (if normalized by the length of *tf-5*)<br><br>TM-score = 0.69685 (if normalized by the length of 1MSE) |

TM align:

TM-align is an algorithm for sequence-independent protein structure comparisons. For two protein structures of unknown equivalence, TM-align first generates optimized residue-to-residue alignment based on structural similarity using heuristic dynamic programming iterations. An optimal superposition of the two structures built on the detected alignment, as well as the TM-score value which scales the structural similarity, will be returned. TM-score has the value in (0,1], where 1 indicates a perfect match between two structures. (https://zhanggroup.org/TM-align/)

**Reference:**

**Zhang Y, Skolnick J**. (2005) TM-align: A protein structure alignment algorithm based on TM-score. *Nucleic Acids Research* **33**: 2302-2309

**Table S3.** Morphological characterization of AV and *tf-5*. Values in bold show the significant difference ($P \leq 0.05$) as compared to the AV. Data are mean ± SE ($n \geq 4$).

| | **Plant morphology** | **AV** | ***tf-5*** |
|---|---|---|---|
| Vegetative (2- month old) | Height (mm) | 440.0±9.00 | **259.0±10.15** |
| | Internode length (mm) | 26.33±0.43 | **15.50±0.12** |
| | No. of branches | 4.125±0.52 | **7.25±0.25** |
| Reproductive (3-4 month old) | No. of inflorescences | 6.75±0.41 | **12.25±0.49** |
| | No. of fruits/plant | 8.25±0.52 | **12.25±0.59** |
| Fruit phenotype (~3-month-old, first truss) | Size (horizontal in mm) | 47.59±1.33 | **48.76±0.67** |
| | Size (vertical in mm) | 36.73±0.99 | **29.61±1.03** |

**Table S4.** Carotenoids profiles of *tf* alleles and their respective WT. Values in bold show the significant difference (P ≤ 0.05) with respect to corresponding WT. Data are mean ± SE (n ≥4) and expressed as µg/gm FW. The MG and BR stage of *tf-1* could not be ascertained, hence the carotenoids for these two stages were not examined, and therefore the respective cells are labeled as NE (Not examined). The ND (Not detected) refers to the stages where the carotenoid analysis was carried out, but respective carotenoid was not detected, perhaps due to being below the limit of the detection.

| **Carotenoids** | **Fruit stage** | ***tf-1*** | **CR** | ***tf-2*** | **RH** | ***tf-3*** | **M82** | ***tf-4*** | ***tf-z*** | **AV** | ***tf-5*** |
|---|---|---|---|---|---|---|---|---|---|---|---|
| Phytoene | MG | NE | ND | ND | ND | ND | ND | ND | ND | ND | ND |
| | BR | NE | ND | ND | ND | ND | ND | ND | ND | ND | ND |
| | RR | 0.15±0.01 | ND | **47.32±5.96** | 43.78±3.25 | 37.42±4.26 | 20.82±2.72 | 28.99±3.57 | 28.78±4.21 | 32.33±2.55 | **69.62±6.02** |
| Phytofluene | MG | NE | ND | ND | ND | ND | ND | ND | ND | ND | ND |
| | BR | NE | ND | ND | ND | ND | ND | ND | ND | ND | ND |
| | RR | 0.15±0.01 | ND | **3.4±0.59** | 3.1±0.23 | 3.83±0.48 | 2.17±0.23 | **1.09±0.17** | 1.69±0.23 | ND | **11.46±0.41** |
| ζ-Carotene | MG | NE | ND | ND | ND | ND | ND | ND | ND | ND | **0.22±0** |
| | BR | NE | ND | ND | ND | ND | ND | ND | ND | ND | **0.22±0.02** |
| | RR | ND | 16.82±1.47 | **ND** | ND | ND | 7.01±0.72 | ND | ND | 8.98±0.92 | 9.35±1.6 |
| Lycopene | MG | NE | ND | ND | ND | ND | ND | ND | ND | 0.3±0.02 | 0.51±0.09 |
| | BR | NE | ND | ND | ND | ND | ND | ND | ND | 0.42±0.04 | 1.02±0.2 |
| | RR | 2.11±0.19 | 40±4.26 | **63.6±6.63** | 24.7±2.19 | 20.85±2 | 39.19±1.87 | 40.04±6.05 | 33.2±3.76 | 56.63±4.56 | **108.48±9.15** |
| β-Carotene | MG | NE | 0.91±0.12 | 0.88±0.09 | 1.05±0.05 | **0.76±0.03** | 0.84±0.1 | 0.69±0.02 | 1.18±0.1 | 0.68±0.09 | **1.75±0.17** |
| | BR | NE | 1.28±0.11 | 1.21±0.18 | 1.11±0.08 | 1.2±0.16 | 0.95±0.05 | 1.06±0.11 | 1.05±0.12 | 0.99±0.23 | **2.37±0.26** |
| | RR | 8.11±0.87 | 7.86±0.42 | **5.81±0.52** | 6.25±0.36 | **4.65±0.2** | 4.39±0.43 | 5.4±0.25 | **8.75±0.46** | 2.95±0.21 | **4.81±0.21** |
| γ-Carotene | MG | NE | ND | ND | ND | ND | ND | ND | ND | ND | ND |
| | BR | NE | ND | ND | ND | ND | ND | ND | ND | ND | **0.29±0.03** |

| **Carotenoids** | **Fruit stage** | ***tf-1*** | **CR** | ***tf-2*** | **RH** | ***tf-3*** | **M82** | ***tf-4*** | ***tf-z*** | **AV** | ***tf-5*** |
|---|---|---|---|---|---|---|---|---|---|---|---|
| | RR | ND | ND | ND | ND | ND | ND | ND | ND | ND | **2.32±0.15** |
| β-Cryptoxanthin | MG | NE | ND | ND | ND | ND | ND | ND | ND | ND | ND |
| | BR | NE | ND | ND | ND | ND | ND | ND | ND | ND | ND |
| | RR | ND | ND | ND | ND | ND | ND | ND | ND | 2.33±0.49 | **1.09±0.09** |
| Antheraxanthin | MG | NE | ND | ND | ND | ND | ND | ND | ND | ND | **0.88±0.01** |
| | BR | NE | ND | ND | ND | ND | ND | ND | ND | ND | **0.87±0** |
| | RR | ND | ND | ND | ND | ND | ND | ND | ND | ND | ND |
| Violaxanthin | MG | NE | 0.38±0.07 | 0.51±0.04 | 0.46±0.04 | 0.45±0.02 | 0.25±0.02 | 0.28±0.02 | **0.52±0.1** | 0.54±0.04 | **0.98±0.04** |
| | BR | NE | 0.24±0.01 | **0.37±0.04** | 0.32±0.05 | 0.4±0.08 | 0.22±0.01 | 0.24±0.02 | **0.31±0.02** | 0.38±0.04 | **0.95±0.18** |
| | RR | ND | ND | ND | ND | ND | ND | ND | ND | ND | **1.43±0.16** |
| Neoxanthin | MG | NE | 1.08±0.17 | 1.28±0.11 | 0.83±0.04 | **1.07±0.08** | 0.31±0.05 | **0.49±0.03** | **1.32±0.18** | 0.26±0.02 | **1.05±0.19** |
| | BR | NE | 0.68±0.04 | 0.79±0.07 | 0.46±0.08 | **0.89±0.15** | 0.28±0.05 | 0.48±0.13 | **0.69±0.09** | 0.06±0.01 | **0.63±0.12** |
| | RR | ND | ND | ND | ND | ND | ND | ND | ND | ND | ND |
| δ-Carotene | MG | NE | ND | ND | ND | ND | ND | ND | ND | ND | ND |
| | BR | NE | ND | ND | ND | ND | ND | ND | ND | ND | ND |
| | RR | ND | ND | 5.87±0.5 | 5.38±0.36 | **3.94±0.26** | ND | 4.46±0.44 | 5.25±0.4 | 2.15±0.47 | **3.56±0.28** |
| α-Carotene | MG | NE | ND | ND | ND | ND | 0.38±0.03 | 0.34±0.06 | 0.38±0.04 | ND | **0.2±0.02** |
| | BR | NE | ND | ND | ND | ND | 0.38±0.07 | 0.32±0.05 | 0.18±0.02 | ND | **0.25±0.01** |
| | RR | ND | ND | ND | ND | ND | ND | **1.35±0.05** | **1.11±0.04** | 1.02±0.08 | **2.04±0.15** |
| Lutein | MG | NE | 1.71±0.28 | **2.52±0.19** | 1.69±0.11 | 1.86±0.12 | 1.25±0.13 | 1.54±0.12 | **2.34±0.33** | 0.94±0.06 | **4.5±0.25** |
| | BR | NE | 1.36±0.14 | 1.75±0.2 | 1.4±0.12 | 1.82±0.25 | 1.27±0.09 | **1.69±0.09** | 1.58±0.12 | 0.6±0.05 | **3.52±0.35** |
| | RR | 1.18±0.2 | 1.14±0.06 | 1.69±0.29 | 1.59±0.11 | 1.52±0.03 | 1.28±0.08 | **1.82±0.16** | 1.56±0.14 | 0.94±0.12 | 0.79±0.05 |

| **Carotenoids** | **Fruit stage** | ***tf-1*** | **CR** | ***tf-2*** | **RH** | ***tf-3*** | **M82** | ***tf-4*** | ***tf-z*** | **AV** | ***tf-5*** |
|---|---|---|---|---|---|---|---|---|---|---|---|
| Total carotenoid | MG | NE | 4.08±0.64 | 5.19±0.43 | 4.03±0.24 | 4.14±0.25 | 3.03±0.33 | 3.34±0.25 | 5.74±0.75 | 2.73±0.22 | 10.1±0.77 |
| | BR | NE | 3.56±0.3 | 4.12±0.49 | 3.29±0.33 | 4.31±0.64 | 3.1±0.27 | 3.79±0.4 | 3.81±0.37 | 2.45±0.36 | 10.13±1.2 |
| | RR | 11.7±1.3 | 65.82±6.21 | 127.7±14.5 | 84.8±6.5 | 72.2±7.23 | 75±6.05 | 83.2±10.7 | 80.3±9.24 | 107.34±9.4 | 215±18.1 |

**NE, not examined; ND, not detected**

**Table S5.** Folate, *p*-aminobenzoic acid (*p*ABA), pterin-6-carboxylic acid (p6C), and 6-hydroxymethylpteridine (HMPt) content in *tf* alleles and their respective WT fruits. Folate and pathway intermediates value are in µg/100 gm FW. Data are means ± SE (n=5), and the values in bold show the significant difference (P ≤ 0.05). NE, not examined; ND, not detected (below the detection limit).

| Cultivar/mutant | MG | BR | RR |
|---|---|---|---|
| **Folate** | | | |
| *tf-1* | NE | NE | 37.47±1.02 |
| CR | 27.43±2.14 | 19.7±0.42 | 25.78±1.37 |
| *tf-2* | **50.43±5.08** | **34.91±1.21** | **39.85±3.14** |
| RH | 26.54±0.35 | 27.19±1.34 | 33.05±2.38 |
| *tf-3* | **29.45±0.56** | 30.79±1.19 | **26.61±0.85** |
| M82 | 29.83±0.65 | 31.97±1.86 | 27.26±1 |
| *tf-4* | 33.12±1.48 | 31.22±1.23 | **32.63±1.31** |
| *tf-z* | **42.93±2.15** | **41.18±0.93** | **39.92±1.78** |
| AV | 24.6±1.21 | 19.3±1.5 | 18.3±1.2 |
| *tf-5* | **56.9±3.5** | **55.2±4.4** | **70.1±1.7** |
| **pABA** | | | |
| *tf-1* | NE | NE | 21.88±1.9 |
| CR | 3.89±0.18 | 4.65±0.33 | 5.75±0.25 |
| *tf-2* | 4.27±0.34 | **6.29±0.38** | **8.05±0.49** |
| RH | 3.22±0.06 | 3.83±0.2 | 4.53±0.16 |
| *tf-3* | 3.36±0.15 | 3.98±0.16 | 4.24±0.26 |
| M82 | 8.58±0.36 | 10.56±0.22 | 11.92±0.54 |
| *tf-4* | **11.18±0.87** | **15.4±0.43** | **19.9±0.96** |
| *tf-z* | 8.88±0.8 | **8.52±0.33** | **9.75±0.47** |
| AV | 2.5±0.4 | 2.5±0.14 | 3.6±0.13 |
| *tf-5* | **6.7±0.25** | **8.1±0.38** | **10.9±0.58** |

| Cultivar/mutant | MG | BR | RR |
|---|---|---|---|
| **p6C** | | | |
| *tf-1* | NE | NE | 6.76±0.77 |
| CR | 3.46±0.55 | 2.91±0.3 | 3.99±1.05 |
| *tf-2* | **2.07±0.2** | 2.38±0.54 | **1.29±0.32** |
| RH | 4.58±0.47 | 4.46±0.34 | 5.64±0.51 |
| *tf-3* | 5.13±0.44 | 4.57±0.45 | 6.07±0.32 |
| M82 | 7.37±1.05 | 7.27±1.23 | 11.09±0.73 |
| *tf-4* | **4.05±0.71** | 5.3±0.59 | **4.36±0.67** |
| *tf-z* | **3.62±0.36** | 5.11±0.14 | **3.88±0.68** |
| AV | 13.1±0.87 | 8.2±0.69 | 5.8±0.27 |
| *tf-5* | **4.1±0.47** | **4.1±0.62** | **1.8±0.35** |
| **HMPt** | | | |
| *tf-1* | NE | NE | ND |
| CR | 0.42±0.11 | 0.43±0.14 | ND |
| *tf-2* | ND | ND | ND |
| RH | 0.2±0.12 | 0.2±0.05 | ND |
| *tf-3* | 0.28±0.07 | 0.31±0.11 | 0.35±0.11 |
| M82 | 1.41±0.35 | ND | ND |
| *tf-4* | ND | 0.6±0.09 | 1.64±0.31 |
| *tf-z* | ND | 0.32±0.06 | 0.63±0.16 |
| AV | 2.88±0.16 | 2.51±0.35 | ND |
| *tf-5* | ND | ND | ND |

**Table S6.** List of primers set used for mutation detection in *tf-5* using CEL I assay. The primers were designed using SOL ITAG 2.3.

| **Gene** | **Primers** | | **Primer length** | **Primer Sequence (5′→3′)** | **Start-End position (bp)** |
|---|---|---|---|---|---|
| *MYB117* (Solyc05g007870) | Set 1 | FP | 21 | CTTGTTCTTGTTCTCCTATGG | 20 |
| | | RP | 20 | GAGCAGCAAATGGTGAGTAT | 1240 |
| | Set 2 | FP | 20 | TTATGGACCAAATGGCTCAC | 1131 |
| | | RP | 21 | AGTGGCTCCTACTCCAAGAAA | 1973 |
| | Sub-Set 2 | FP | 23 | AACACGGGTGCGTATGACTTTAT | 1554 |
| | | RP | 24 | GGCTCCTACTCCAAGAAAGTCTAT | 1970 |

**Table S7.** List of primers used for qRT-PCR analysis for folate biosynthesis pathway, *MYB117*, and *FLAs* in AV and *tf-5*. The primers were designed using SOL ITAG 2.3.

| ′Gene | SGN id. | Primer Sequence (5′→3′) | | Amplicon size (bp) |
|---|---|---|---|---|
| *ADCS* | Solyc04g049360 | Fp | TGAAAGAAGGGCTCATTATGCT | 148 |
| | | Rp | CTGGATGTGATAGATTTAAGGTTCC | |
| *ADCL1* | Solyc11g071280 | Fp | TCGATAGGGAAAGCATAAGACAGA | 110 |
| | | Rp | ATAGTTGAAAATCACCAGGTCCTG | |
| *GCH1* | Solyc06g083230 | Fp | AGGATGCTGTTAGAGTCCTATTGC | 103 |
| | | Rp | TGTTCCTTGTCTTAGAGCCTTAGC | |
| *DPP (DHPN)* | Solyc03g043860 | Fp | ATGAGGAAAATCTTGCATCACACT | 116 |
| | | Rp | CATACCATTCCCATCCATCACATT | |
| *DHNA (DHPA)* | Solyc10g079830 | Fp | CAAGTATCCAGAGGTATCTGCTGTT | 99 |
| | | Rp | GTATCTAATGATCTCGACACCCAAG | |
| *DHFS* | Solyc06g051900 | Fp | CACTTCGAGTACCCTATGTAGCAAT | 113 |
| | | Rp | CGTAAAGTTCAGATGCTACATCCTT | |
| *DHFR* | Solyc01g109830 | Fp | GTTCAGGAAGTTTTGACATTGCTAC | 103 |
| | | Rp | AATAGAGAGACAATAAGGCGAGGAT | |
| *HPPK-DHPS* | Solyc05g012090 | Fp | ACTTTATGCAACTTAGGATCAGTGG | 114 |
| | | Rp | GTTGACGTTAGCTTTAGACTTCTGC | |
| *FPGSm* | Solyc04g016550 | Fp | GAGCT TGGACAAACG GTAGTATTTA | 114 |
| | | Rp | TTTCCAGGAGTGTATGAGGGTATAG | |
| *FPGSp* | Solyc05g052920 | Fp | TACTTCTCTGCTC TACGCTTCAAAT | 111 |
| | | Rp | AGTTGGCTTTATCTCATCCTTACCT | |
| *GGH1* | Solyc07g062270 | Fp | GAAGATTTCCCCCCGTGTTGCTAAAGA | 128 |
| | | Rp | CAATACCCTGAAAAAGCTACTCAAA | |
| *GGH2* | Solyc10g007410 | Fp | CCCGATTATATATACAGAGCCTCCT | 140 |
| | | Rp | CTTCAAAGTAGAGACCCTTCTTGCT | |
| *MYB117* | Solyc05g007870 | Fp | CTTGTTCTTGT TCTCCTATGG GTA | 85 |
| | | Rp | AGACTTTCTGGAGTGTTAATAGAG | |
| *FLAs10* | Solyc10g005960 | Fp | CCATGGTGGTCTTCTCTTTAGGT | 106 |
| | | Rp | TGGTTGAAAGTGGAAAATTGTGGG | |
| *FLAs13* | Solyc01g091530 | Fp | TGTGTTGGAAGTTAGTGCTCCGA | 117 |
| | | Rp | GAAGCAAATGTTTTACACCCAGCT | |
| *β-ACTIN* | FJ532351.1 | Fp | GTCCCTATTTACGAGGGTTATGC | 108 |
| | | Rp | CAGTTAAATCACGACCAGCAAGATT | |
| *UBIQUITIN 3* | X58253.1 | Fp | GCCGACTACAACATCCAGAAGG | 110 |
| | | Rp | TGCAACACAGCGAGCTTAACC | |

*ADCS*, Aminodeoxychorismate synthase; *ADCL*, Aminodeoxychorismate lyase; *GCHI*, GTP cyclohydrolase I; *DPP*, Dihydroneopterin triphosphate pyrophosphatase; *DHNA*, Dihydroneopterin aldolase; *DHFS*, Dihydrofolate synthetase; *DHFR*, Dihydrofolate reductase; *HPPK-DHPS*, Hydroxymethyldihydropterin pyrophosphokinase – dihydropteroate synthase; *FPGSm*, Folylpolyglutamate synthase (mitochondrial); *FPGSp*, Folylpolyglutamate synthase (plastidial); *GGH*, γ-glutamyl hydrolase; *FLAs*, Fasciclin-like arabinogalactan proteins; *MYB*, myeloblastosis.

**Table S8.** List of primers used for qRT-PCR analysis for carotenoid biosynthesis pathway and ripening regulators in AV and *tf-5* fruit. The primers were designed using SOL ITAG 2.3.

| Gene | SGN id. | Primer Sequence (5′→3′) | | Amplicon size (bp) |
|---|---|---|---|---|
| *DXS* | Solyc01g067890 | FP | AAATGGGATCGGTGTAGAGC | 115 |
| | | RP | TGCTGAGCCATATCCCAATA | |
| *GGPPS2* | Solyc04g079960 | FP | ATCAATGGAGCAGCTTTGTG | 128 |
| | | RP | GCGGTTGATAAAACGACGTA | |
| *PSY1* | Solyc03g031860 | FP | TGAATTAGCACAGGCAGGTC | 140 |
| | | RP | TCAATTCTGTCACGCCTTTC | |
| *PDS* | Solyc03g123760 | FP | TATCATCAACGTTCCGTGCT | 122 |
| | | RP | TATCGGTTTGTGACCAGCAT | |
| *ZISO* | Solyc12g098710 | FP | AGAGCGTGCTTTTCGTGTATTG | 107 |
| | | RP | ATTGCCATAACTGCACTCCATC | |
| *ZDS* | Solyc01g097810 | FP | TCCAAAAGGGCTATTTCCAC | 115 |
| | | RP | TTGATCCAAGAGCTCCACAG | |
| *CRTISO* | Solyc10g081650 | FP | GAGATCGCCAAATCCTTAGC | 118 |
| | | RP | CAGAAAGCTTCACTCCCACA | |
| *LCYB1* | Solyc04g040190 | FP | CGATGCAACTGGCTTCTCTA | 149 |
| | | RP | AATGAGAATCTCGCCAATCC | |
| *CYCB* | Solyc06g074240 | FP | TCTTCTCAAGCCTTTTCCATC | 92 |
| | | RP | TGGTGGGACTTAGAAAAGAAGG | |
| *LCYE* | Solyc12g008980 | FP | TTAGTCGCCATTTTCTGCAC | 130 |
| | | RP | TCACCCTCGCACTCTACAAG | |
| *ZEP2* | Solyc02g090890 | FP | GGTCGTGTTACATTGCTTGG | 118 |
| | | RP | TGCATGCTTTTTCAAGTTCC | |
| *VDE* | Solyc04g050930 | FP | GTGCAGCTAATGTTGCCTGT | 126 |
| | | RP | GGGAGACTGCACACTCATTG | |
| *CYP97A* | Solyc04g051190 | FP | GTGCCATTGTACCAGCATTG | 101 |
| | | RP | TGCAGCAACATCAAGCTTTT | |
| *CYPC11* | Solyc10g083790 | FP | TGCTGAGAGAATGGTGGAGA | 107 |
| | | RP | GTGCAAGGCCAATAACATCA | |
| *NCED* | Solyc07g056570 | FP | TGACACCACCAGACTCCATT | 130 |
| | | RP | ACTTGTTCATCCGGGTTTTC | |
| *NOR* | Solyc10g006880 | FP | GGCAATATTCGGAGAGCAAG | 113 |
| | | RP | TTCCGGTAGCCTTCCAATAA | |
| *RIN* | Solyc05g012020 | FP | ATGGCATTGTGGTGAGCAAAG | 147 |
| | | RP | GTTGATGGTGCTGCATTTTCG | |
| *CNR* | Solyc02g077920 | FP | AAATTTGGGCTGAAGAAGCA | 115 |
| | | RP | AATCCGGGAATTGACAGAAG | |

*DXS*, deoxy-xylulose 5-phosphate synthase; *GGPPS*, geranylgeranyl diphosphate synthase; *PSY1*, phytoene synthase 1; *PDS*, phytoene desaturase; *ZISO*, ζ-carotene isomerase; *CRTISO*, carotenoid isomerase; *ZDS*, ζ-carotene desaturase; *LCYB1*, lycopene β-cyclase1; *CYCB*, chromoplast specific lycopene β-cyclase; *LCYE*, lycopene ε-cyclase; *ZEP*, zeaxanthin epoxidase; *VDE*, violaxanthin deepoxidase; *CYP97A29*, cytochrome P450 carotenoid β-hydroxylase A29; *CYP97C11*, cytochrome P450 carotenoid ε-hydroxylase C11; *NCED*, 9-cis-epoxycarotenoid dioxygenase; *NOR*, Nonripening; *RIN*, ripening inhibitor; *CNR*, Colorless nonripening.

**Table S9.** The filtered read statistics of the transcriptome of AV and *tf-5* red ripe fruit. The sequencing reads with low quality, adaptor-polluted, and high content of unknown base (N) reads were removed before downstream analyses using SOAPnuke (v1.5.2).

| **Sample** | **Total Raw reads** | **Reads with unknown bases(N) > 5%** | **Adaptor contaminated reads** | **Low quality reads (Q < 15)** | **Total Clean reads (fastq files, %)** | **Clean Reads Q20 (%)** | **Clean Reads Q30 (%)** |
|---|---|---|---|---|---|---|---|
| **AV** | 115454306 | 723794 (0.63%) | 536406 (0.46%) | 4657532 (4.03%) | 109536574 (94.9) | 98.7 | 92.8 |
| **AV** | 115426526 | 709804 (0.61%) | 640238 (0.55%) | 4346184 (3.77%) | 109730300 (95.1) | 98.7 | 92.9 |
| **AV** | 116323850 | 322854 (0.28%) | 502002 (0.43%) | 4270100 (3.67%) | 111228894 (95.6) | 98.0 | 91.5 |
| ***tf-5*** | 115315664 | 731266 (0.63%) | 901176 (0.78%) | 4587066 (3.98%) | 109096156 (94.6) | 98.7 | 92.8 |
| ***tf-5*** | 116318562 | 314798 (0.27%) | 880638 (0.76%) | 4480406 (3.85%) | 110642720 (95.1) | 97.9 | 91.0 |
| ***tf-5*** | 115498828 | 280080 (0.24%) | 711710 (0.62%) | 3698334 (3.2%) | 110808704 (95.9) | 98.8 | 93.3 |

**Chen Y, Chen Y, Shi C, Huang Z, Zhang Y, Li S, Li Y, Ye J, Yu C, Li Z, Zhang X, Wang J, Yang H, Fang L, Chen Q**. (2018). SOAPnuke: a MapReduce acceleration-supported software for integrated quality control and preprocessing of high-throughput sequencing data. *Gigascience* **7:**1– 6. https://doi.org/10.1093/gigascience/gix120

**Table S10.** Summary of genome mapping statistics using HISAT2 in AV and *tf-5* red ripe fruit.

| Sample | Total clean reads | Unique mapping (%) | Multiple map reads (%) | Total mapping Ratio % |
|---|---|---|---|---|
| **AV** | 109536574 | 51132528 (93.36%) | 1252894 (2.29%) | 97.99% |
| **AV** | 109730300 | 51264664 (92.50%) | 1530276 (2.76%) | 97.75% |
| **AV** | 111228894 | 50626596 (91.03%) | 1576893 (2.84%) | 96.59% |
| ***tf-5*** | 109096156 | 50432228 (91.46%) | 1868529 (3.39%) | 97.66% |
| ***tf-5*** | 110642720 | 50878425 (91.45%) | 1517283 (2.73%) | 97.18% |
| ***tf-5*** | 110808704 | 51546774 (93.04%) | 1654381 (2.99%) | 98.37% |


**Kim D, Paggi JM, Park C, Bennett C, Salzberg SL.** (2019) Graph-based genome alignment and genotyping with HISAT2 and HISAT-genotype. *Nature Biotechnology*. **37**: 907-15.

**Table S11.** Genes name and their promoter sequence coordinate used for MYB binding site. The 2000 bp upstream promoter sequence of folate and carotenoids biosynthesis pathway genes were retrieved from SGN (https://solgenomics.net/). The MYB binding sites were and analyzed using PlantCARE (http://bioinformatics.psb.ugent.be/webtools/plantcare/html/)

| Gene Abbreviation | SOLYC Number | SL2.50 Coordinates |
|---|---|---|
| *ADCS:* | Solyc04g049360.2 | (>SL2.50ch04:41633624…41630625) |
| *ADCL1:* | Solyc11g071280.1 | (>SL2.50ch11:54816405…54813406) |
| *GCH1:* | Solyc06g083230.2 | (>SL2.50ch06:48746174…48743175) |
| *DPP (DHPN):* | Solyc03g043860 | (>SL2.50ch03:7532694...7529695) |
| *DHNA (DHPA):* | Solyc10g079830.1 | (>SL2.50ch10:61312710...61315709) |
| *DHFS:* | Solyc06g051900.2 | (>SL2.50ch06:35577695...35580694 |
| *DHFR:* | Solyc01g109830 | (>SL2.50ch01:96682187...96679188) |
| *FPGSm:* | Solyc04g016550 | (>SL2.50ch04:7383851…7380852 |
| *FPGSp:* | Solyc05g052920 | (>SL2.50ch05:63106376…63103377) |
| *GGH1:* | Solyc07g062270 | (>SL2.50ch07:65047840…65050839) |
| *GGH2:* | Solyc10g007410 | (>SL2.50ch10:1782740…1779741) |
| *DHPS* | Solyc05g012090 | (>SL2.50ch05:5313864…5310865) |
| *DXS1:* | Solyc01g067890 | (>SL2.50ch01:76875760…76872761) |
| *DXR:* | Solyc03g114340.2 | (>SL2.50ch03:64344674…64347673) |
| *GGPPS2:* | Solyc04g079960.1 | (>SL2.50ch04:64280892...64277893) |
| *PSY1:* | Solyc03g031860.2 | (>SL2.50ch03:4323134…4326133) |
| *PDS:* | Solyc03g123760 | (>SL2.50ch03:70498011…70501010) |
| *ZISO:* | Solyc12g098710.1 | (>SL2.50ch12:66122700…66125699) |
| *ZDS:* | Solyc01g097810 | (>SL2.50ch01:88526734…88523735) |
| *CRTISO:* | Solyc10g081650.1 | (>SL2.50ch10:62689988…62686989) |
| *LCYB1:* | Solyc04g040190.1 | (>SL2.50ch04:11944053…11947052) |
| *CYCB:* | Solyc06g074240.1 | (>SL2.50ch06:45902723...45899724) |
| *LCYE:* | Solyc12g008980.1 | (>SL2.50ch12:2282372...2285371) |
| *ZEP:* | Solyc02g090890 | (>SL2.50ch02:52378308...52375309) |
| *VDE:* | Solyc04g050930.2 | (>SL2.50ch04:48990853...48987854) |
| *NCED:* | Solyc07g056570.1 | (>SL2.50ch07:64366163…64363164) |
| *MYB117:* | Solyc05g007870.1 | (>SL2.50ch05:2309616…2312615) |
| *RIN:* | Solyc05g012020 | (>SL2.50ch05:5233962…5230963) |
| *NOR:* | Solyc10g006880 | (>SL2.50ch10:1311510...1308511) |
| *CNR:* | Solyc02g077920 | (>SL2.50ch02:42748257...42745258) |
