## Supplementary material for "Seeing the unseen: A trifoliate (MYB117) mutant allele fortifies folate and carotenoids in tomato fruits": Figure S1

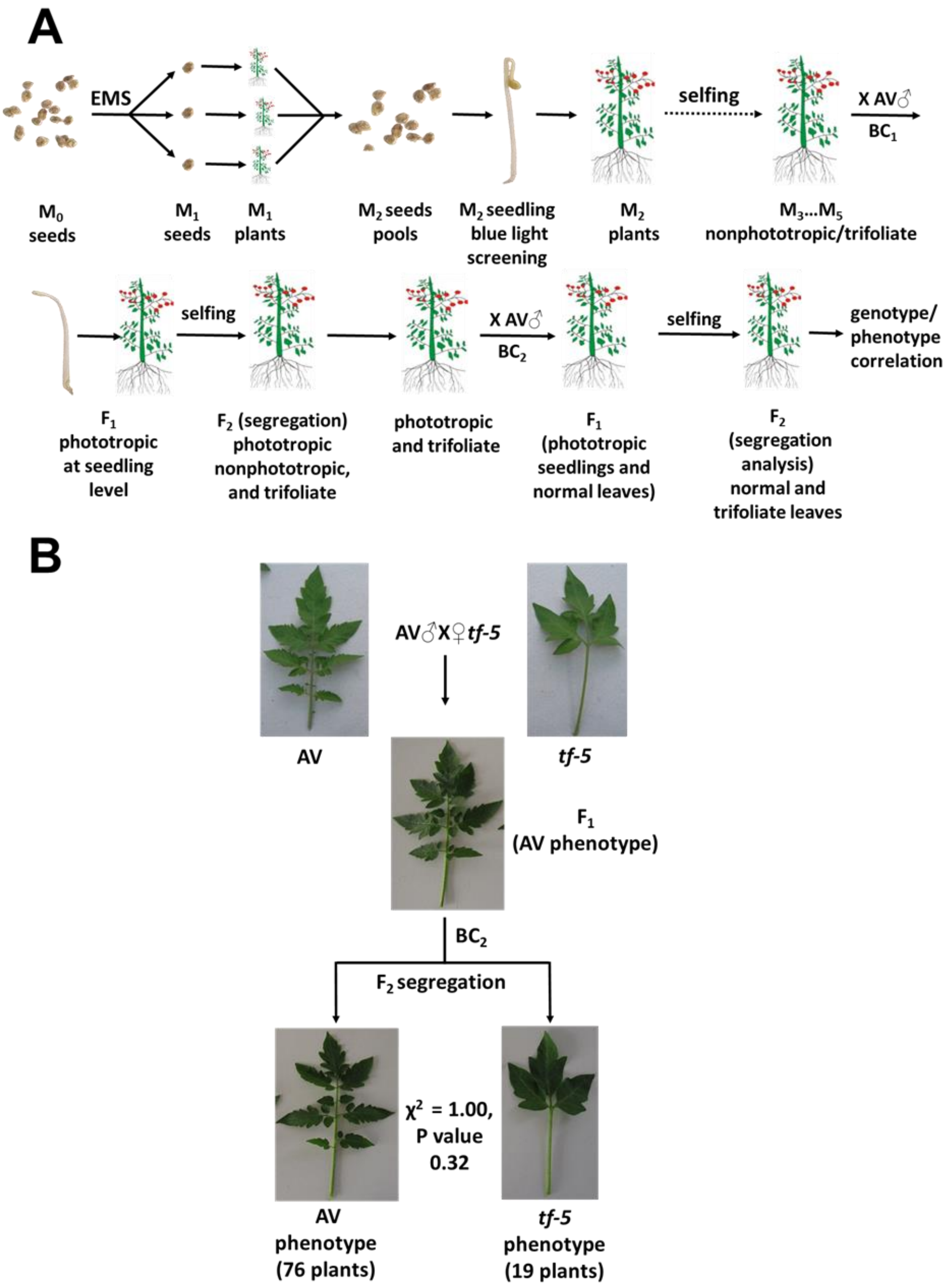


**Figure S1.** *Trifoliate* mutant stabilization and backcrossing.

**A.** The *tf-5* mutant identified in $M_2$ generation was taken forward till $M_5$ generation for ascertaining the stability of the phenotype. It was backcrossed with its parent Arka Vikas in the $M_5$ generation, followed by a second backcross. $BC_1F_2$ and $BC_2F_2$ homozygous plants were identified by trifoliate phenotype. **B**. Genetic segregation of *tf-5* mutant in $BC_2F_2$ generation. The normal leaf and trifoliate phenotype segregated in Mendelian ratio of 3:1 respectively showing that the *tf-5* is a monogenic recessive allele. The stability of the trifoliate phenotype was again confirmed in $BC_2F_3$ generation. In addition, the presence of the mutation in the *MYB117* gene and its homozygosity was also confirmed by CEL-I endonuclease mismatch assay (Figure S2) and Sanger sequencing of *MYB117* gene from wild type and *tf-5*.

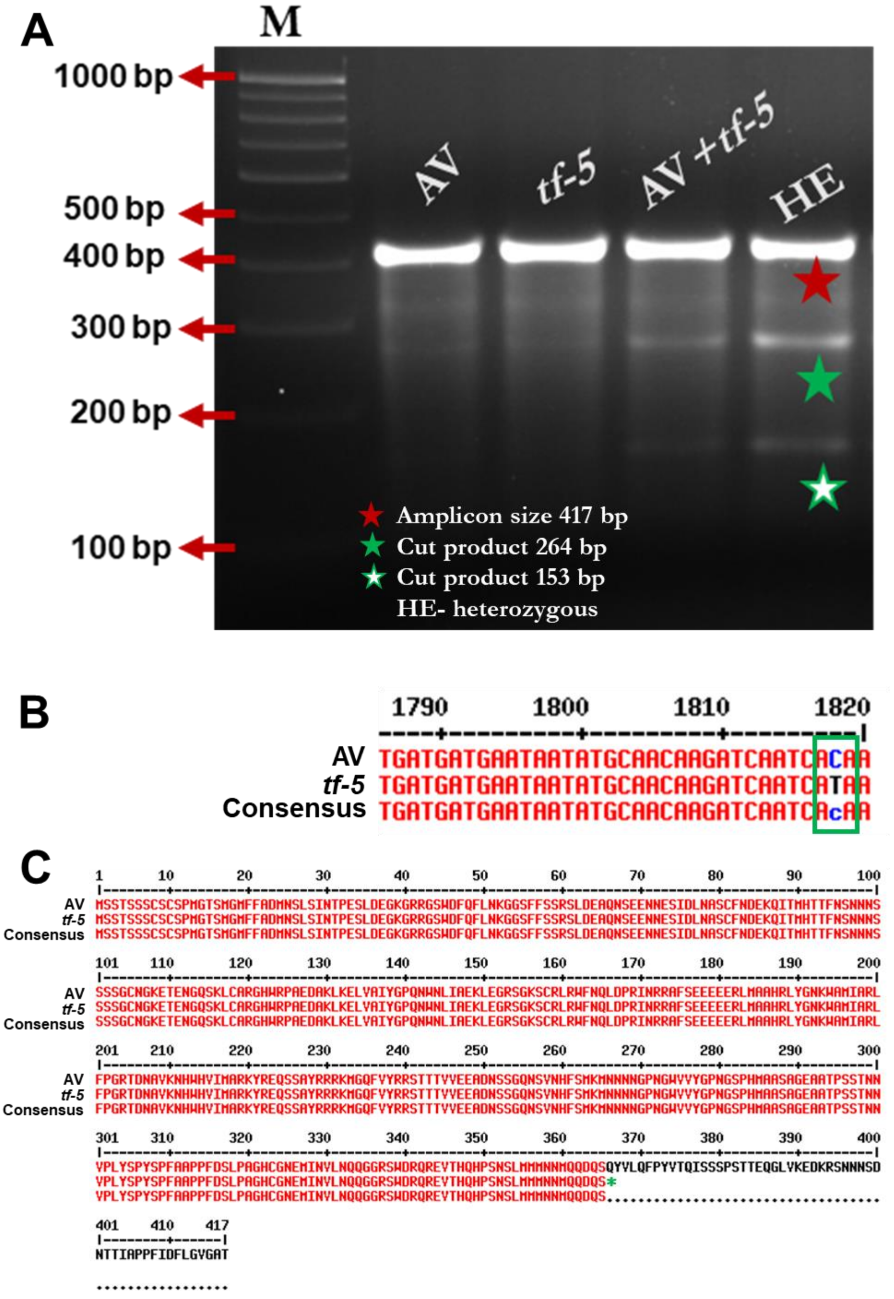


**Figure S2.** Mutation detection in *tf-5* by Sanger Sequencing, and CEL-I endonuclease assay.

**A.** Representative figure showing CEL-I endonuclease assay used for detection of the *tf-5* mutation and its zygosity in backcrossed plants. The genomic DNA from the wild type (AV) and the *tf-5* mutant was PCR amplified (417 bp) with primer set II, denatured, and renatured to make heteroduplex (AV + *tf-5* PCR products) before CEL-I digestion, and gel electrophoresis. The cleavage product of 264 bp and 153 bp generated by CEL-I cleavage is seen only in heteroduplexed CEL-I treated PCR product, but not in AV and *tf-5* mutant PCR products (homoduplex) incubated with CEL-I. The zygosity of backcrossed plants was determined by CEL-I assay followed by Sanger sequencing. **B.** *MYB117* gene sequence displaying the nucleotide change in *tf-5* (C1818T). **C.** The alignment of the MYB117 protein sequence of WT and *tf-5* showing that a truncated protein of 365 amino acids is formed in *tf-5*. The change in a nucleotide at position C1818T resulted in the replacement of glutamine at 366 position by a stop codon (Q366*). For CEL-I assay *see* Sreelakshmi et al. (2010) NEATTILL: A simplified procedure for nucleic acid extraction from arrayed tissue for TILLING and other high-throughput reverse genetic applications. *Plant Methods* **6:** 3

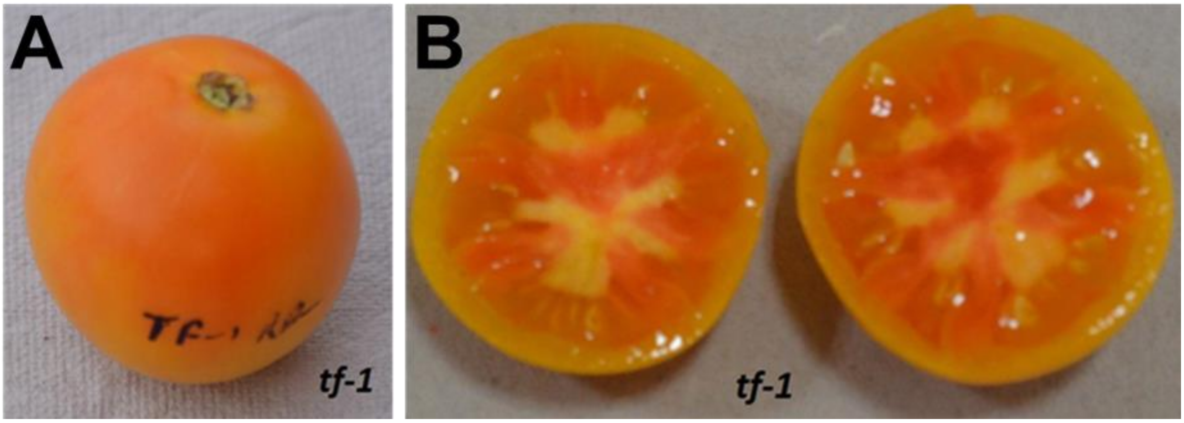


**Figure S3**. The phenotype of ripe *tf-1* fruit.

The *tf-1* fruits are tangerine-coloured and accumulate little lycopene, but have a normal level of β-carotene and lutein. The *tf-1* (LA0512) is not a monogenic mutant, it also harbors *macrocalyx*, *obscuravenosa*, and *wilty* mutations, which may contribute to the absence of carotenoids.

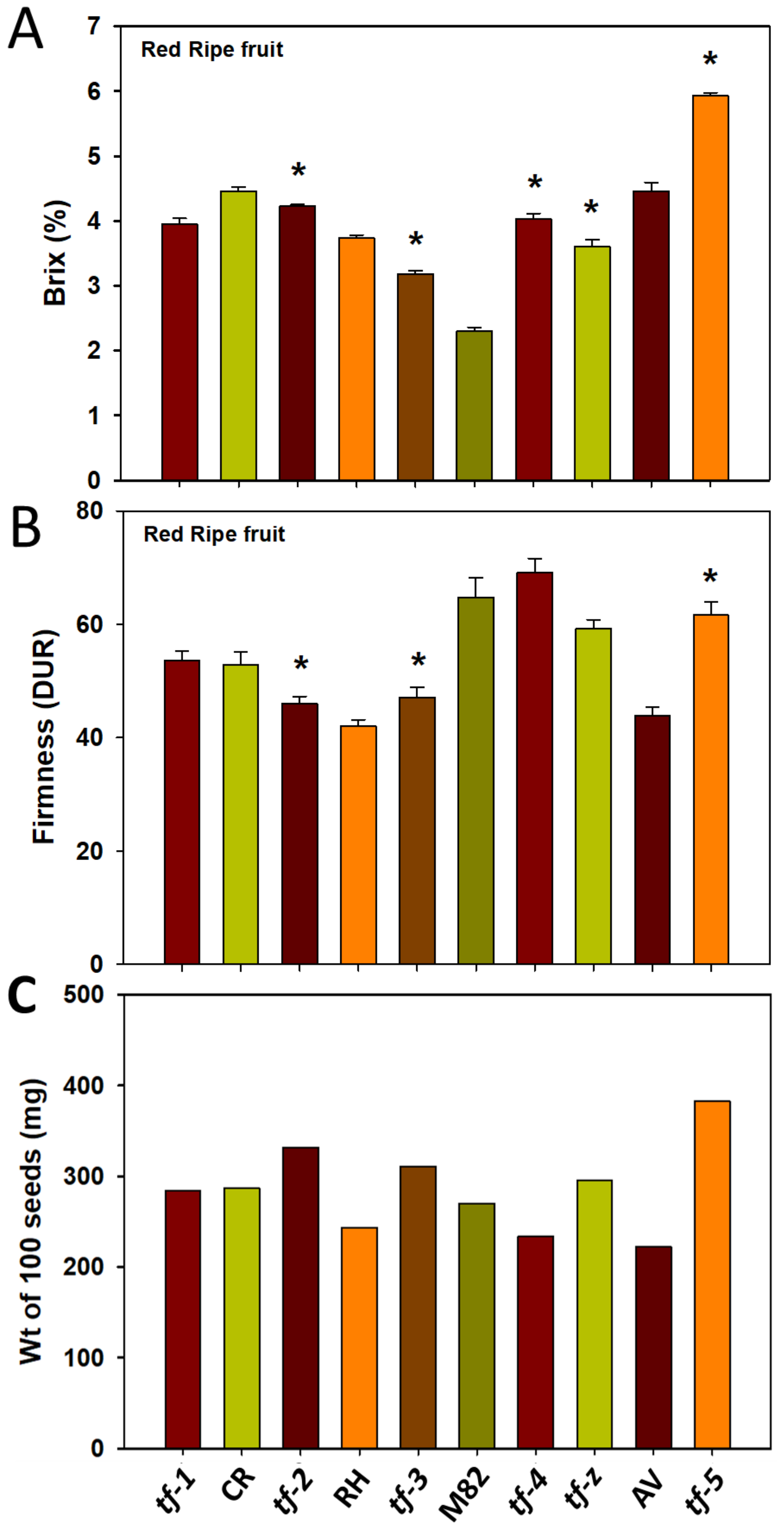


**Figure S4. C**hanges in total soluble sugar content (°Brix), fruit firmness and seeds weigh in different *tf* alleles.

**A.** Total soluble sugar content in different *tf* alleles was measured at red ripe stage along with respective wild-types. **B.** The fruit firmness in different *tf* alleles was measured at red ripe stage along with respective wild-types. **C.** Seeds weight in different tf alleles. Note *tf-5* red ripe fruits have significantly high °BRIX and firmness than its wild type Arka Vikas. Data are means ± SE (n = 3 ), *p ≤ 0.05.

**AV**, Arka Vikas; **RH**, Rheinland Ruhm; **CR**, Condine Red.

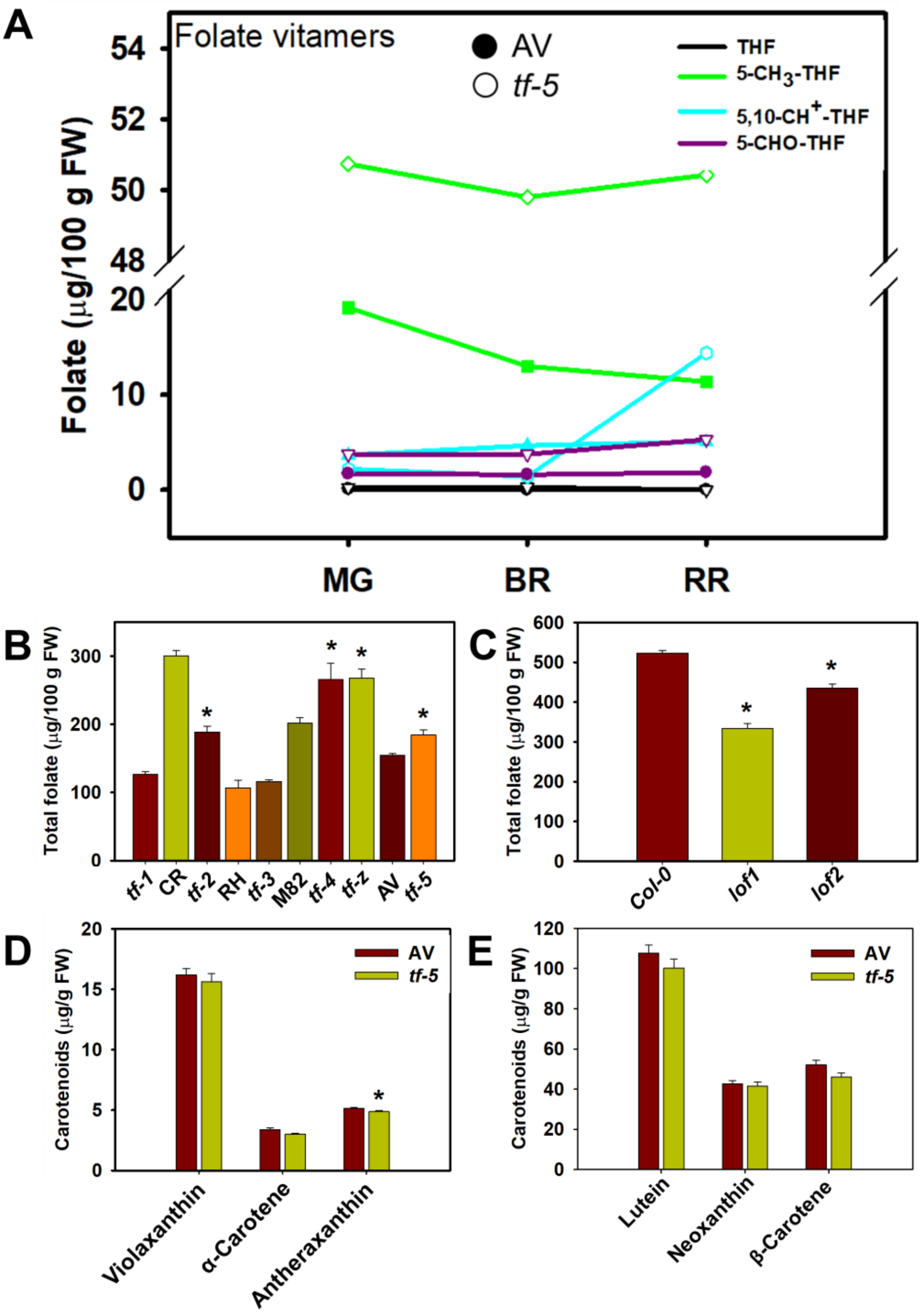


**Figure S5**. Total folate levels in Arabidopsis *lof* mutants, different *tf* alleles, carotenoids level of *tf-5* in leaves and different folate forms in *tf-5* fruit.

**A.** Different folate forms in *tf-5* and AV fruit at MG, BR and RR stage. **B.** Total folate levels in leaf of different *tf* alleles was measured along with respective wild-types. The leaves harvested from 5$^{th}$ node of 7-week-old *tf* mutants and wild type plants were used for folate estimation. **C.** Total folate levels in leaf of *lof1* (*MYB117*) and *lof2* (*MYB105*) mutants of Arabidopsis. The Arabidopsis wild type (CS_70000) and *lof1* (Salk_025235) and *lof2* (Salk_064076) mutant seedlings were grown at 22°C for 21 days under long day (16 h light/8 h dark). **D-E.** The levels of levels different carotenoids in leaf of *tf-5* and AV. The leaves harvested from 5$^{th}$ node of 7-week-old *tf-5* and wild type plants were used for carotenoids estimation. Data are means ± SE (n=3), *$p \leq 0.05$.

*Abbreviations*: THF, tetrahydrofolate; 5-$CH_3$-THF, 5-methyl-THF; 5,10-$CH^+$-THF, 5,10-methenyl-THF; 5-CHO-THF, 5-formyl-THF; AV, Arka Vikas; RH, Rheinland Ruhm; CR, Condine Red; MG, Mature green; BR, Breaker; RR, Red ripe.

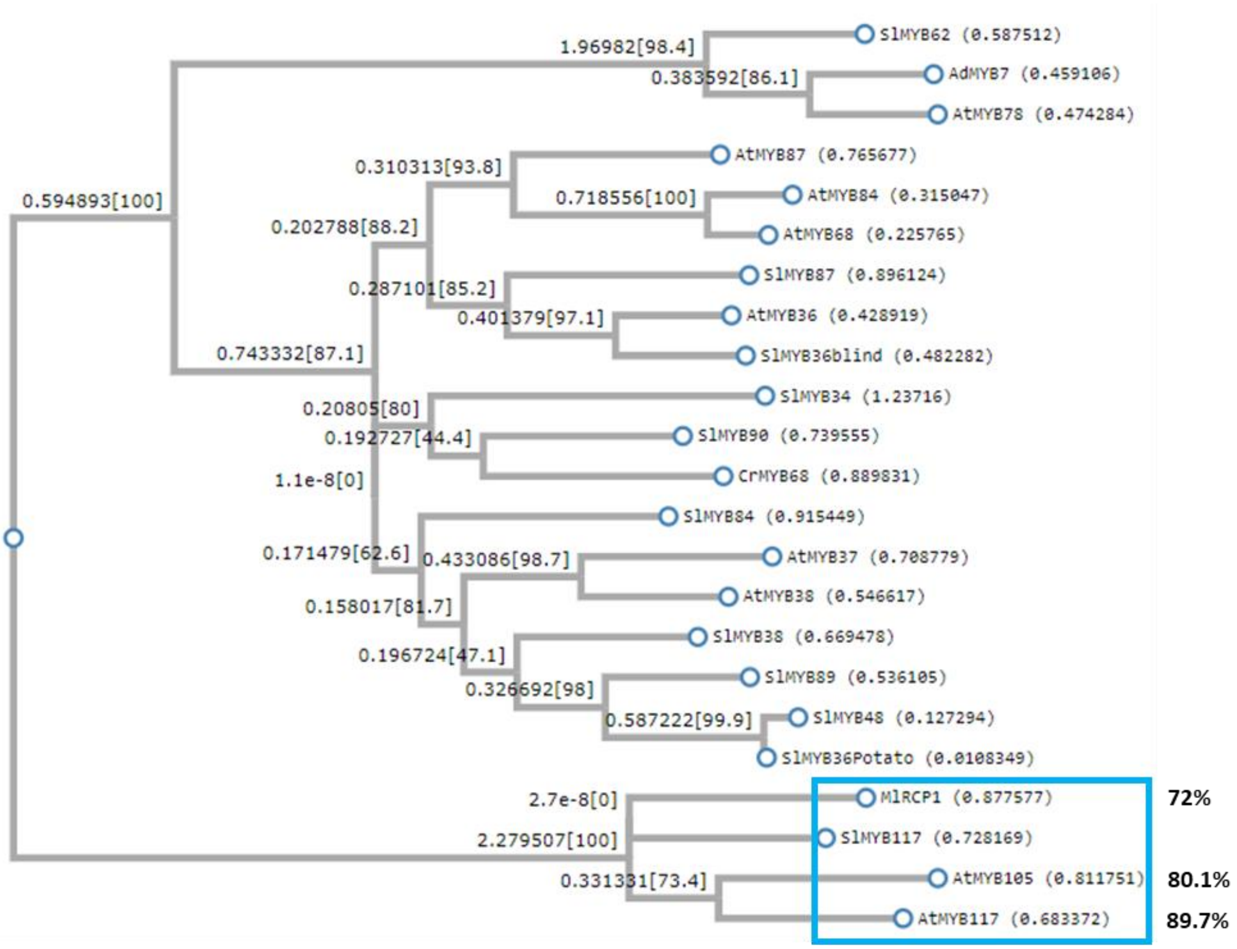


**Figure S6**. Dendrogram showing maximum likelihood phylogeny of selected R2R3-MYBs.

The rectangle encloses the Arabidopsis *lof1 (*MYB117) and *lof2* (MYB105), tomato MYB117, and Mimulus RCP1. The percent homology of the Arabidopsis and Mimulus MYB to tomato MYB 117 is shown on the right side of the rectangle. The effect of loss-of-function mutations in these MYBs widely vary in a species-specific fashion. Arabidopsis *lof1* and *lof2* mutants show reduced folate levels in leaves (this study), whereas *tf-5* mutant shows higher folate level in leaves and fruits. While *rcp1* mutants show reduced carotenoids levels in corolla tube of flowers, the *tf-5* mutant has higher carotenoids levels in fruits.

Lee DK et al. (2009) LATERAL ORGAN FUSION1 and LATERAL ORGAN FUSION2 function in lateral organ separation and axillary meristem formation in Arabidopsis. *Development*. **136**: 2423–32.

Sagawa JM et al. (2016) An R2R3-MYB transcription factor regulates carotenoid pigmentation in Mimulus lewisii flowers. *New Phytologist*. **209**: 1049-57.

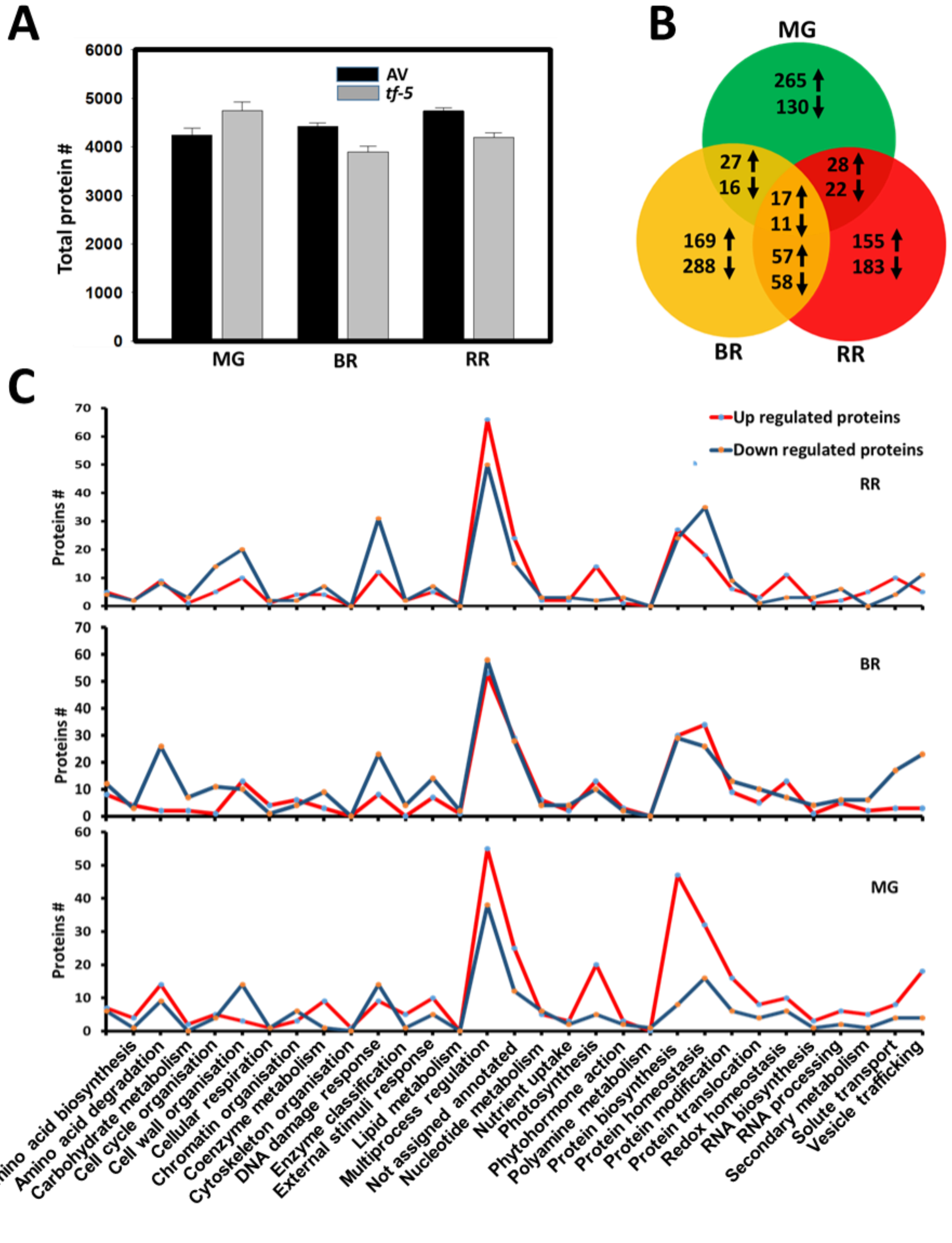


**Figure S7**. Proteome profiling of *tf-5* and its wild type Arka Vikas fruits at different ripening stages**.**

**A.** Number of proteins identified in *tf-5* and Arka Vikas. **B.** Venn representation of differentially expressed proteins in *tf-5* in comparison with Arka Vikas (Log2 fold change ≥ ± 0.58 and p ≤ 0.05). Upregulated (↑), Downregulated (↓). ***Note*** only a few proteins are common between different ripening stages. **C.** Functional classification of differentially expressed proteins in *tf-5* and Arka Vikas. The differentially expressed proteins in the fruits of *tf-5* compared to Arka Vikas were functionally classified using Go-MapMan. See Dataset S2. **MG**, Mature green; **BR**, Breaker; **RR**, Red ripe.

A

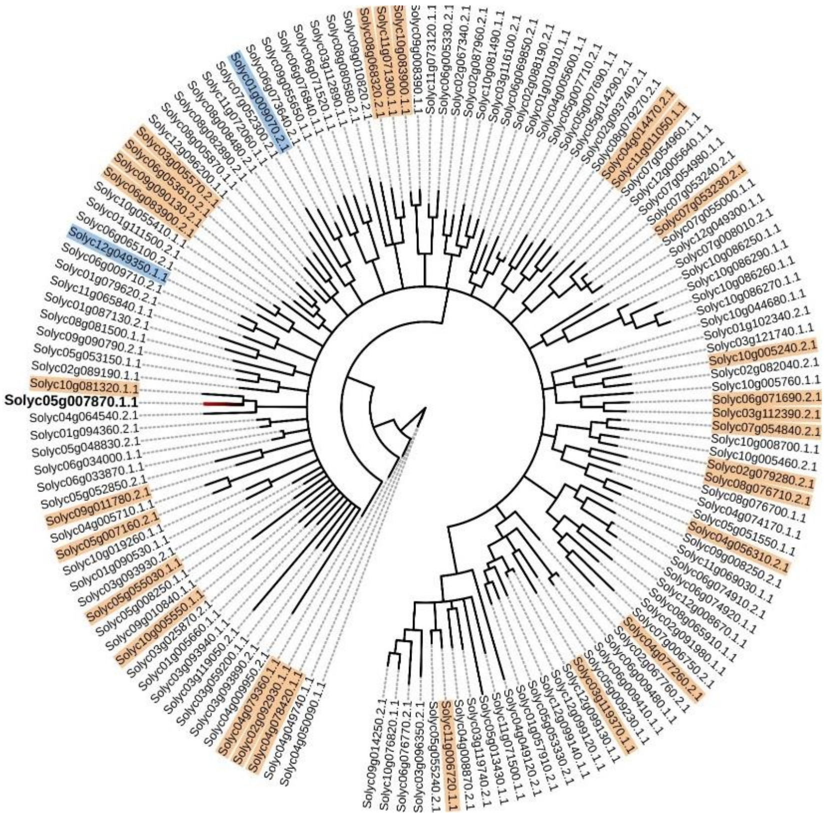


B

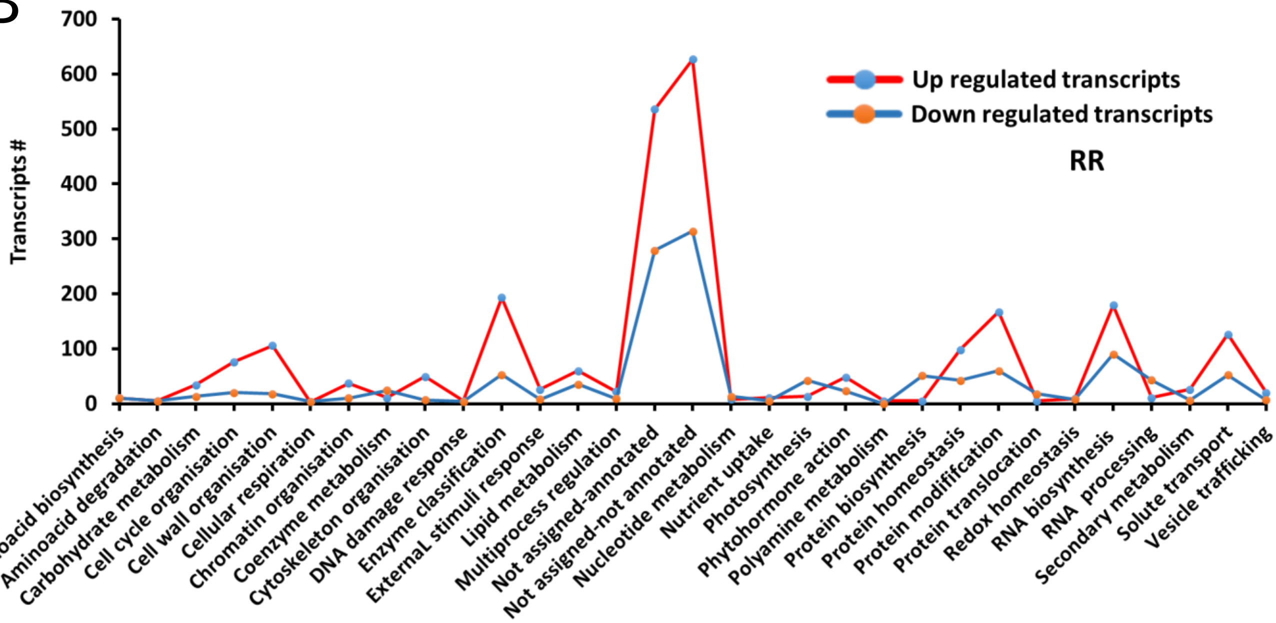


**Figure S8**. Transcriptome profiling of *tf-5* and its wild type Arka Vikas fruits at red ripe stage**.**

**A**. The dendrogram shows the R2R3MYB TF family in tomato. The significantly up- and down-regulated *MYBs* in *tf-5* transcriptome are highlighted in brown and blue color respectively (Log2 fold ≥ ±0.584 and p-value ≤0.05). The *SlMYB117* is shown in bold letter. **B.** Functional classification of differentially expressed transcripts in *tf-5* and Arka Vikas. The differentially expressed transcripts in the red ripe fruits of *tf-5* compared to Arka Vikas were functionally classified using Go-MapMan. RR- Red ripe,

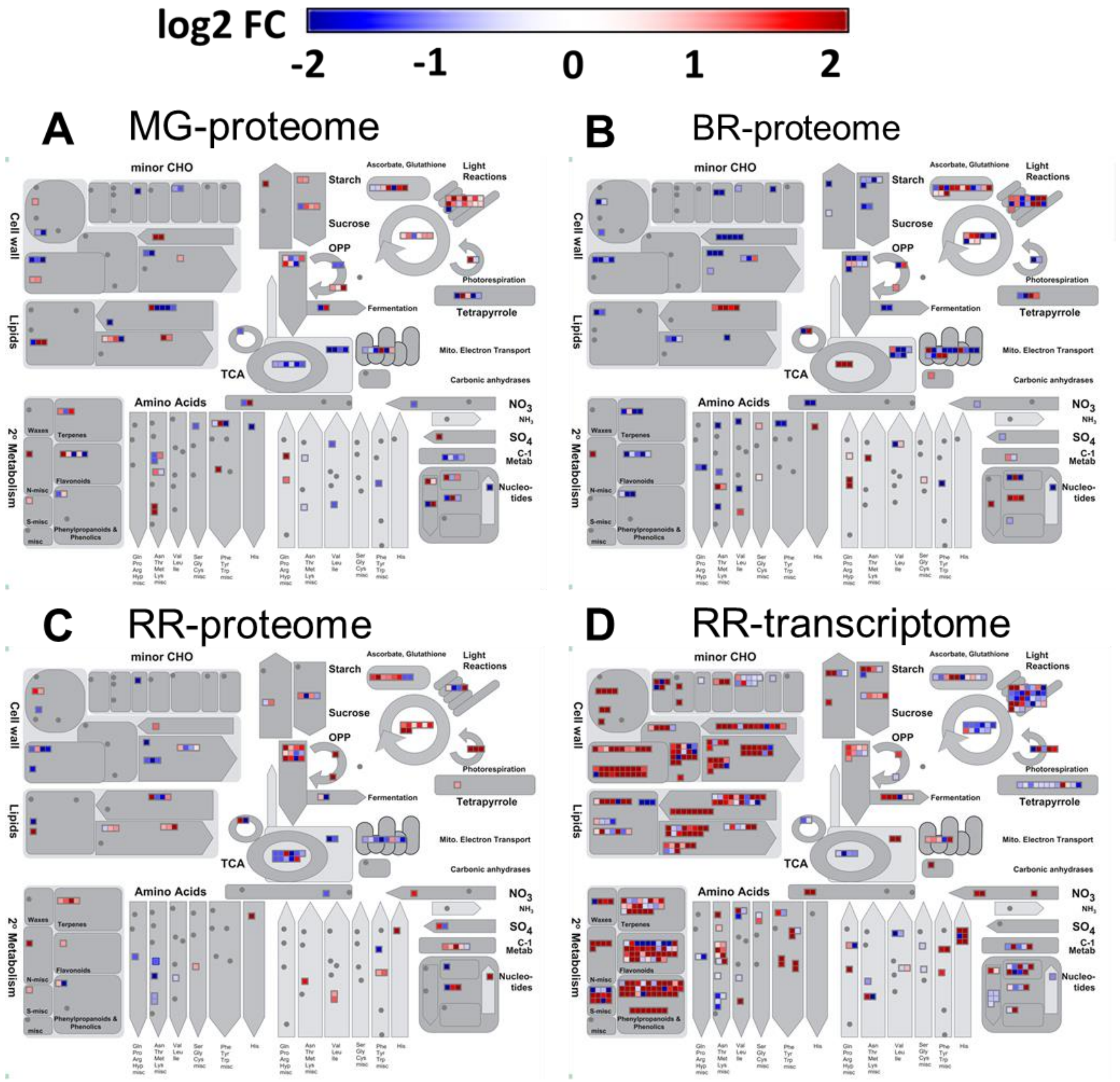


**Figure S9.** Mapman representations of proteome and transcriptome changes in *tf-5* fruits. **A-D.** Mapman overview of metabolic processes affected in *tf-5* at different stages of fruit ripening. **A-C** shows processes affected by differentially expressed proteins at MG **(A)**, BR **(B),** and RR **(C)** stages. **D** shows processes affected by differentially expressed transcripts at RR. Each square represents one protein or transcript. The red square indicates the upregulation of transcripts/proteins and the blue square indicates the downregulation of transcripts/proteins. Only significantly different transcripts/proteins (Log2 fold ≥±0.58, P ≤0.05) are depicted in heat maps. Data are means ± SE (n = 3), P ≤ 0.05. (For details, see Dataset S2, S3 and S4).

**MG**, Mature green; **BR**, Breaker; **RR**, Red ripe.

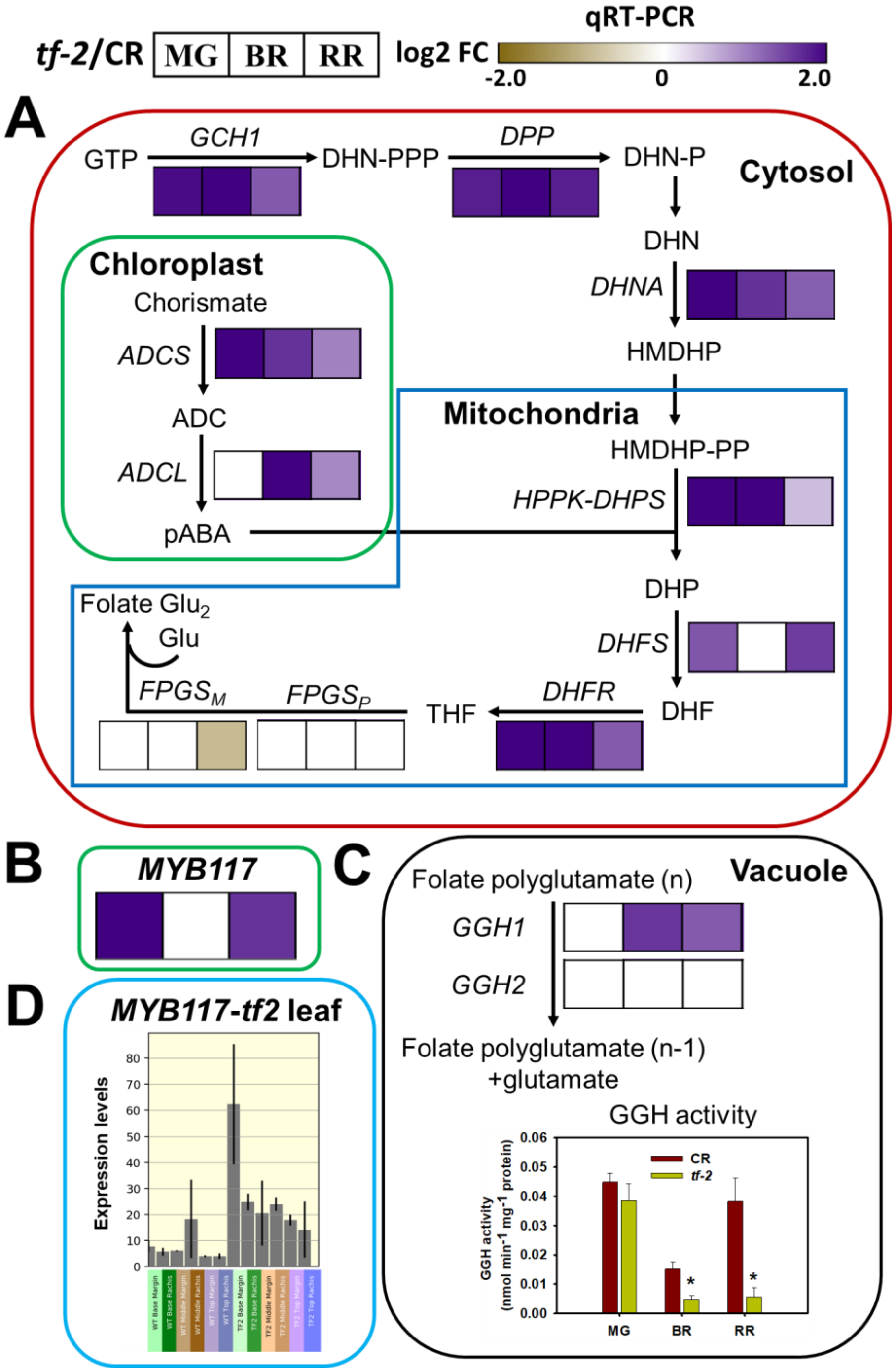


**Figure S11**. Folate biosynthesis pathway, *MYB117* expression, and GGH activity in *tf-2*. **(A-B)** The heat maps show log2 fold change in transcripts of folate biosynthesis pathway genes (**A**), and *MYB117* (**B**) in *tf-2* determined by qRT-PCR at MG, BR, and RR stage**. C** shows change in *GGH1*, *GGH2* transcripts, and total GGH enzyme activity in *tf-2* at MG, BR, and RR stage. **D**, Shows expression of *MYB117* in leaf of *tf-2* mutant and its WT (Data from Martinez et al., 2021, from efp browser http://bar.utoronto.ca/efp_tomato/cgi-bin/efpWeb.cgi?dataSource=Tomato_Meristem. For A, B, and C only significantly different transcripts (Log2 fold ≥±0.58, P ≤ 0.05) are marked with Asterisk on the graphs. *(*For details, see Dataset S1). Data are means ± SE (n = 3), P ≤ 0.05. Abbreviations: GCH1, GTP cyclohydrolase I; DPP, dihydroneopterin (DHN) triphosphate diphosphatase; DHNA, DHN aldolase; ADCS, aminodeoxychorismate (ADC) synthase; ADCL, aminodeoxychorismate lyase; *HPPK-DHPS*, 6-hydroxymethyl-7,8-dihydropterin (HMDHP) pyrophosphokinase (HPPK) and dihydropteroate (DHP) synthase; DHFS, dihydrofolate (DHF) synthase; DHFR, DHF reductase; FPGS, folylpolyglutamate synthase (p-plastic, m-mitochondrial), GGH, γ-glutamyl hydrolase.
MG, Mature green; BR, Breaker; RR, Red ripe.

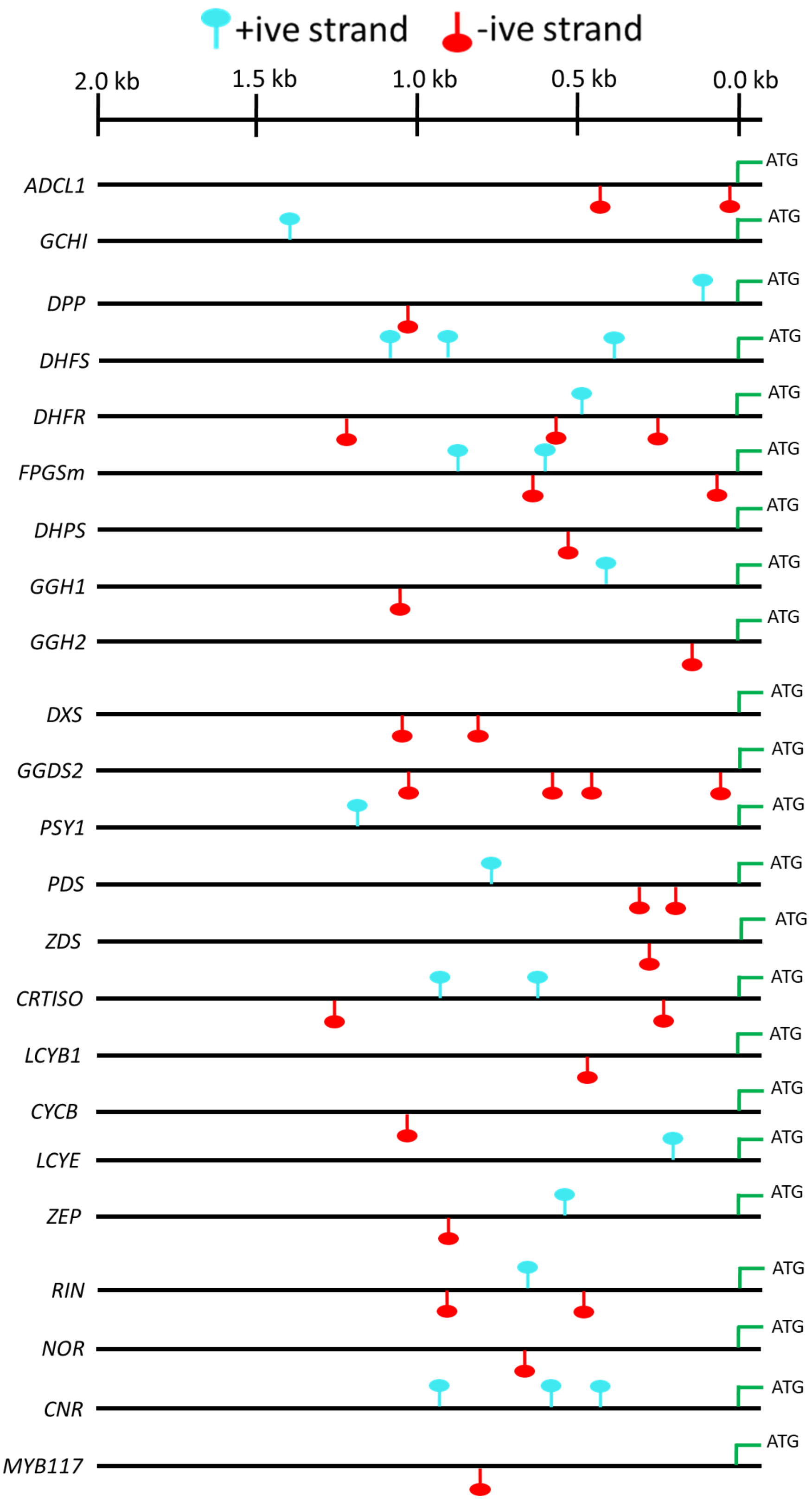


**Figure S12**. MYB binding domains in the 2 kB (-1 to -2000 bp) promoter of carotenoids and folate biosynthesis genes of tomato.

The MYB binding sites were identified in both positive and negative strands of the promoters by using PLACE software. Only genes that had MYB binding sites are shown.

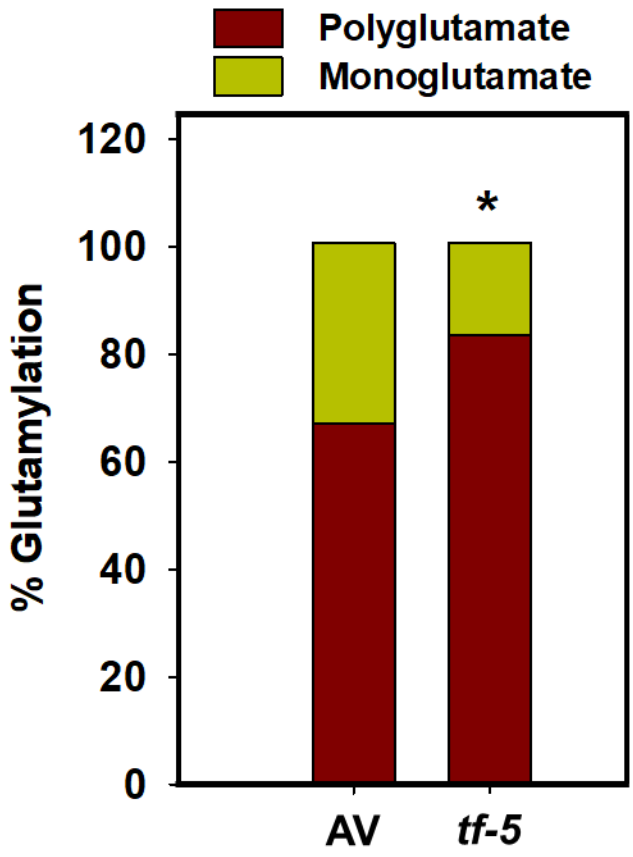


**Figure S13**. Percent polyglutamylation in red ripe fruits of Arka Vikas and *tf-5*.
The reduced GGH activity is associated with higher polyglutamylation of folate. Data are means ± SE (n≥4), *p ≤ 0.05.

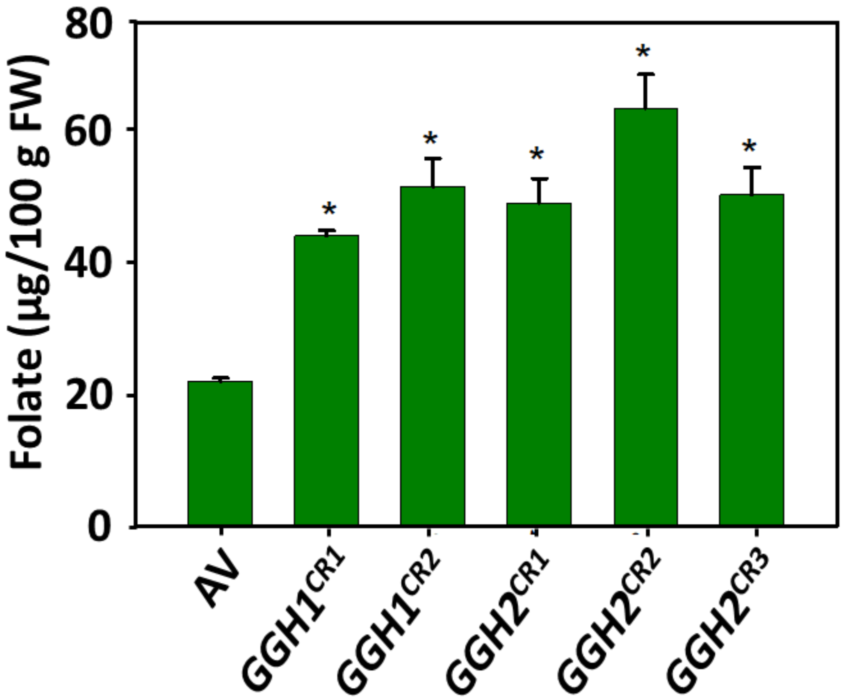


**Figure S14.** Total folate in red ripe fruits of $T_0$ *GGH1* and *GGH2* gene-edited lines. **Note:** Edited lines have 2-3 fold higher folate level than wild type (Arka Vikas). Data are means ± SE (n=3), *$p \leq 0.05$.

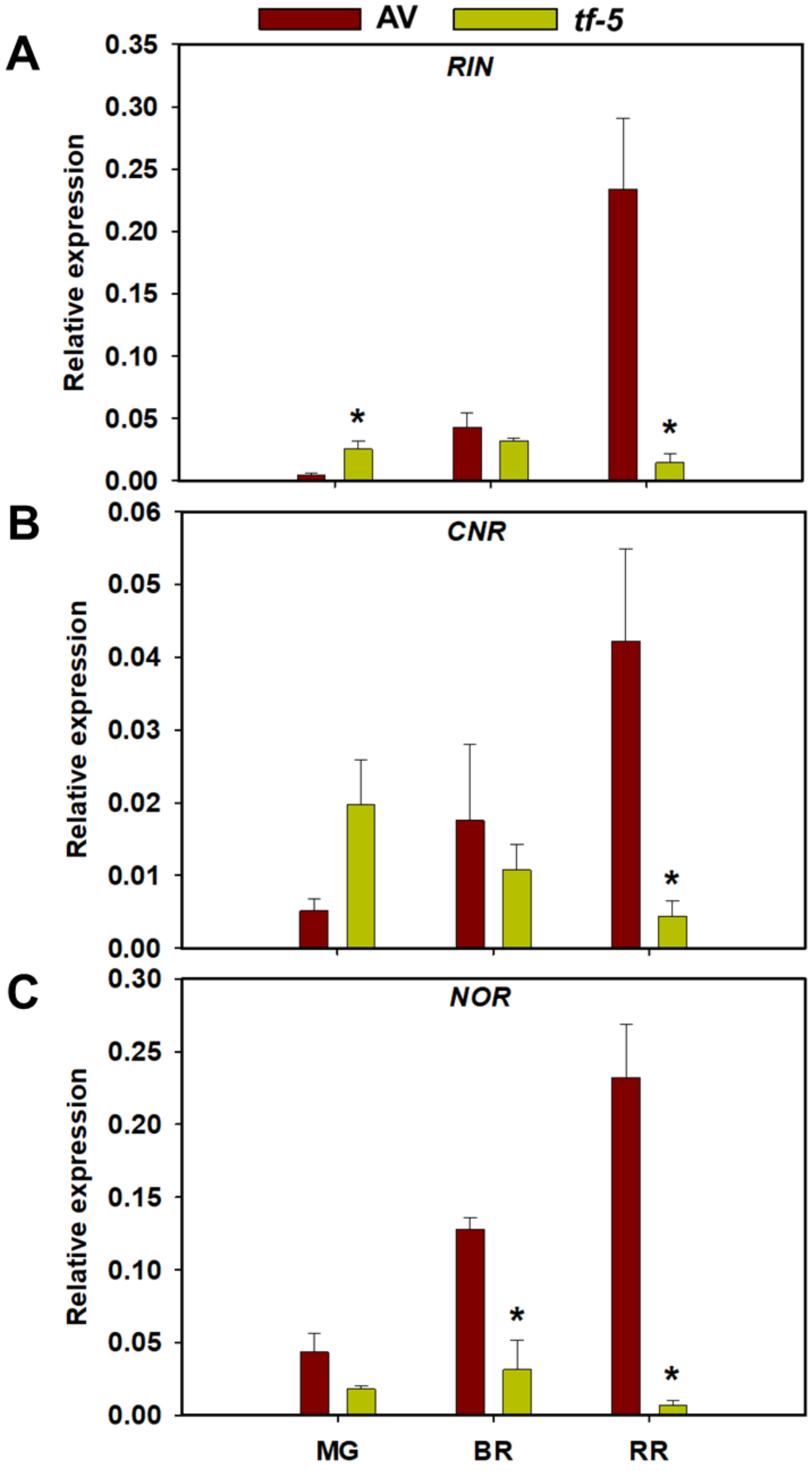


**Figure S15**. Expression of ripening regulator genes *RIN*, *NOR*, and *CNR* of *tf-5* and its wild type Arka Vikas fruits at different ripening stages**.**

The graphs depict data obtained after normalization with *β-actin* and *ubiquitin*. Data are means ± SE (n=3), *p ≤ 0.05. MG, Mature green; BR, Breaker; RR, Red ripe.

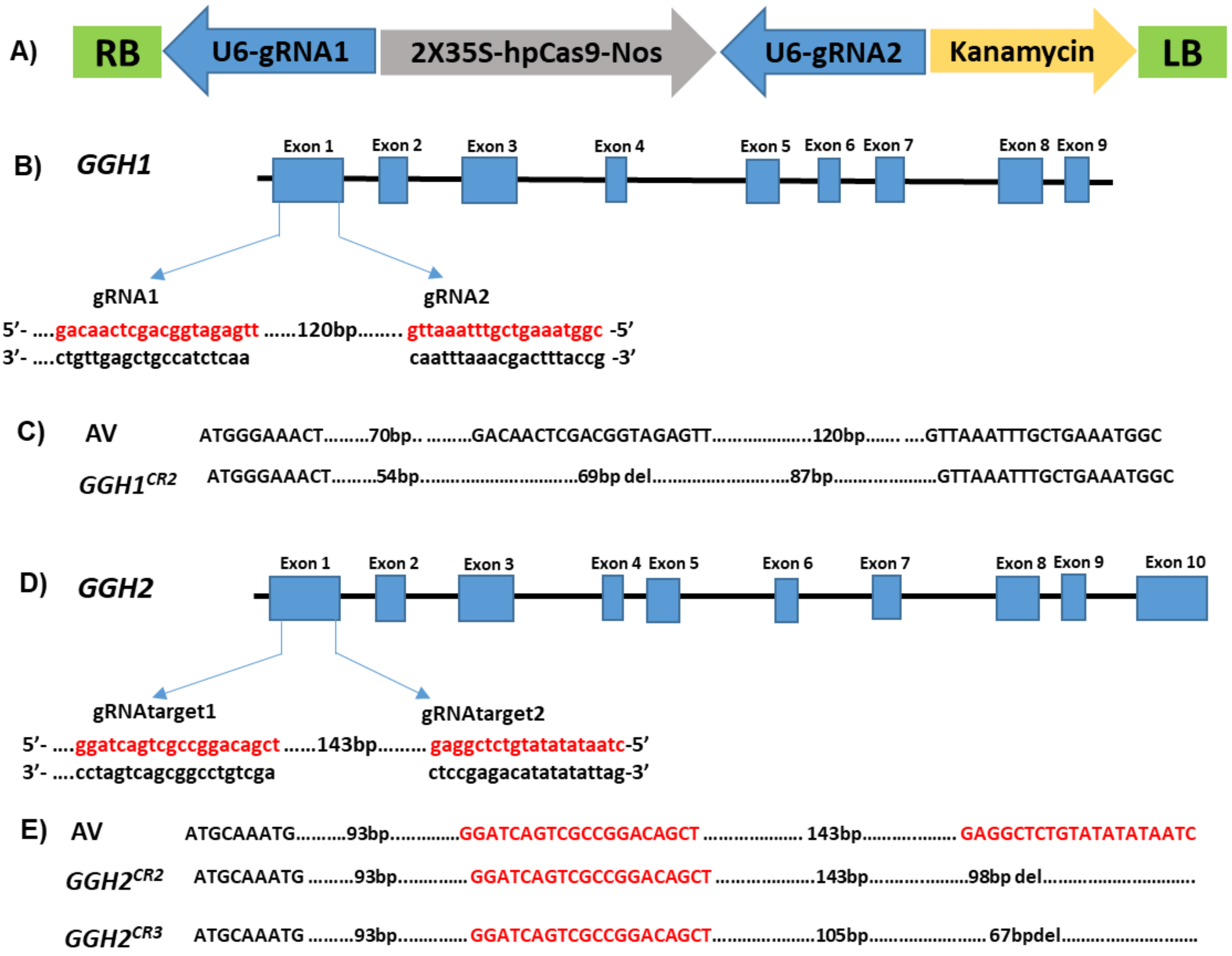


**Figure S16a.** Generation and confirmation of genome-edited *GGH1* and *GGH2* plants.

**A.** Schematic representation of binary vectors used in this study. U6-(*GGH1* and *GGH2* -gRNA-1 and 2: Arabidopsis U6 promoter and the gRNA sequence; 2X 35S-hpCAS9-Nos: 2X CaMV35S promoter sequence; hpCas9: human-codon optimized SpCas9; Nos: Nos terminator; kanamycin: the kanamycin-resistant marker expression cassette; RB: right border of T-DNA; LB: left border of T-DNA.

**B-E.** Schematic view of gRNA1 and gRNA2 target sites in the *GGH1* (**B)** *GGH2* gene (**D)**. Boxes indicate exons, red color indicates 20-bp target sequences. Sequence alignment of the target regions of *GGH1* (**C)** and *GGH2* (**E)** gene. The wild-type sequence is shown at the top, with the target sequence in red. Nucleotide variations at the targets of $T_0$ mutant lines, 'del' are refereed as base deletions.

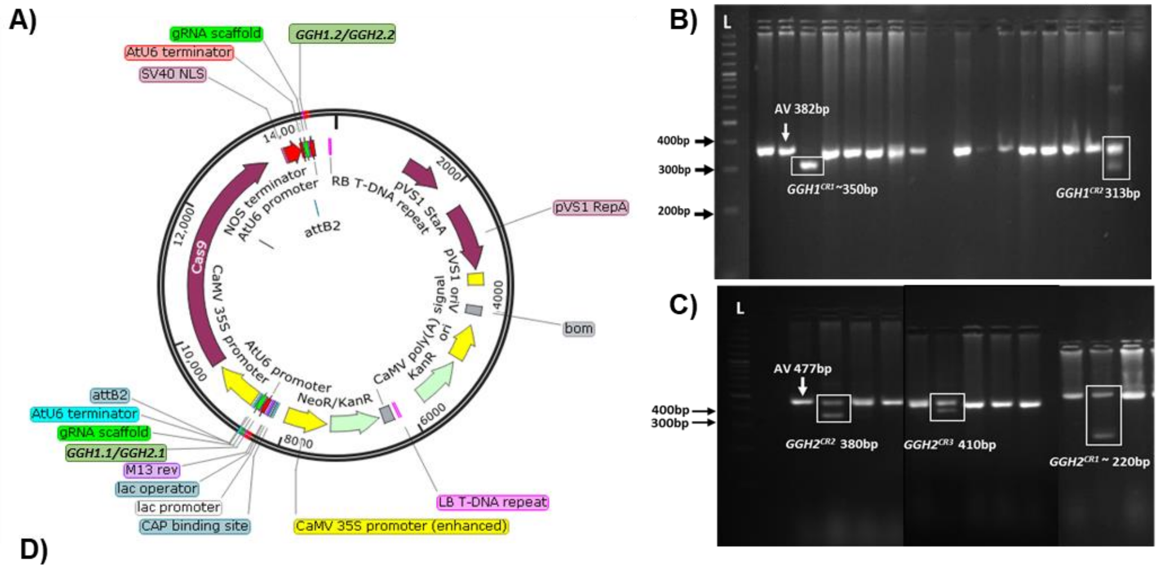


D)

| Alleles | Zygosity in $T_0$ | Mutation description | Effect on translated protein |
|---|---|---|---|
| ***GGH1$^{CR1}$*** | Homozygous | Sequence awaited | |
| ***GGH1$^{CR2}$*** | Heterozygous | 69bp deletion from 93$^{rd}$ -162$^{nd}$ position in the exon1 | Change of 5 amino acids (QLFNS to RSSEL) from 22$^{nd}$ to 26$^{th}$ position and truncation of amino acid from L26$^{*}$ position |
| ***GGH2$^{CR1}$*** | Heterozygous | Sequence awaited | |
| ***GGH2$^{CR2}$*** | Heterozygous | Deletion from 412$^{th}$-450$^{th}$ in exon1 and 451$^{th}$-510$^{th}$ position in the subsequent intron | Deletion from 89$^{th}$ to 112$^{th}$ position |
| **GGH2$^{CR3}$** | Heterozygous | Deletion from 376$^{th}$ -443$^{th}$ position in exon1 | Change of 4 amino acid (VKFA to QSKT) from 104$^{th}$ to 107$^{st}$ position and truncation of amino acid from T108$^{*}$ position |

**Figure S16b.** Generation and confirmation of genome-edited *GGH1* and *GGH2* plants.

**A.** Schematic representation of *γ-glutamyl hydrolase (GGH1* or *GGH2)* dgRNA construct in pCAMBIA2300-cas9 vector showing the location of *GGH1.1*, *GGH1.2* or *GGH2.1*, *GGH2.2:* gRNAs. The above constructs were generated and transformed as described by **Kilambi et al. (2021)**. The double guide RNA (dgRNA) of the *GGH1* (Solyc07g062270) and *GGH2* (Solyc10g007410) gene was selected using the CRISPR-P web tool based on GC content and folded *in silico* by RNAfold server for stable hairpin structure.

**B-C.** Representative image shows the presence of a deleted band in the *GGH1* gene in agarose gel for *GGH1$^{CR1}$* and *GGH1$^{CR2}$* **(B)** and *GGH2$^{CR1}$*,*GGH2$^{CR2}$*, and *GGH2$^{CR3}$* **(C)** Note the presence of a single deleted band in *GGH1$^{CR1}$* and the absence of the WT band indicate the homozygous state of the deletion. The remaining edited lines are in heterozygous conditions.

**D.** Position of deletion in nucleotide and amino acid sequence of *γ-glutamyl hydrolase* (*GGH1* and *GGH2*) edited lines.
