## Supplementary material for "Seeing the unseen: A trifoliate (MYB117) mutant allele fortifies folate and carotenoids in tomato fruits": Figure S10

**Figure S10.** LycoCyc representations of the tomato metabolic pathway with proteome and transcriptome changes in *tf-5* fruits

**A.** The likely changes in the global metabolic pathway in *tf-5* mature-green fruits by alteration in the proteome. **B.** The likely changes in the global metabolic pathway in *tf-5* breaker fruits by alteration in the proteome. **C.** The likely changes in the global metabolic pathway in *tf-5* red-ripe fruits by alteration in the proteome **D.** The likely changes in the global metabolic pathway in *tf-5* red ripe fruits by alteration in the transcriptome

The upregulated (Red-highlight) and downregulated (Blue-highlight) proteins/transcripts of *tf-5* are depicted on the LycoCyc pathway. Only proteins/transcript with log2 fold value ≥ ± 0.58 and p-value of ≤ 0.5 are mapped on the pathway. For full details of protein/transcript SOYC number and changes see Datasets of proteome and transcriptome.

Authors: Bo Xue, Angela Xu, Pascal Schlapfer, Charles Hawkins, Seung Yon Rhee, Emily Fryer, Peifen Zhang

Figure S10A

### TomatoCyc: *Solanum lycopersicum* Cellular Overview

An online version of this diagram is available at BioCyc.org. Biosynthetic pathways are positioned in the left of the cytoplasm, degradative pathways on the right, and reactions not assigned to any pathway are in the far right of the cytoplasm. Transporters and membrane proteins are shown on the membrane. Periplasmic (where appropriate) and extracellular reactions and proteins may also be shown. Pathways are colored according to their cellular function. Connections between pathways are omitted for legibility.

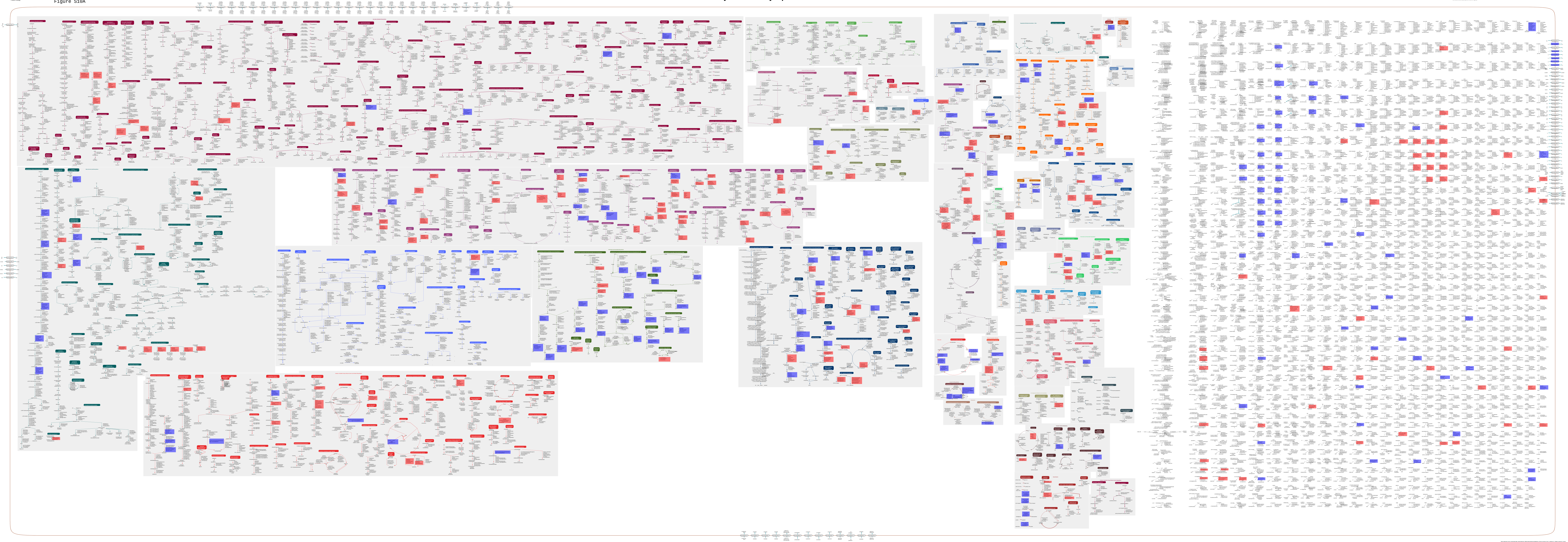

This diagram was automatically generated by SRI International Pathway Tools version 24.0, authors S. Paley and P.D. Karp.

Authors: Bo Xue, Pascal Schlapfer, Seung Yon Rhee, Peifen Zhang, Angela Xu, Charles Hawkins, Emily Fryer

Figure S10B

### TomatoCyc: *Solanum lycopersicum* Cellular Overview

An online version of this diagram is available at BioCyc.org. Biosynthetic pathways are positioned in the left of the cytoplasm, degradative pathways on the right, and reactions not assigned to any pathway are in the far right of the cytoplasm. Transporters and membrane proteins are shown on the membrane. Periplasmic (where appropriate) and extracellular reactions and proteins may also be shown. Pathways are colored according to their cellular function. Connections between pathways are omitted for legibility.

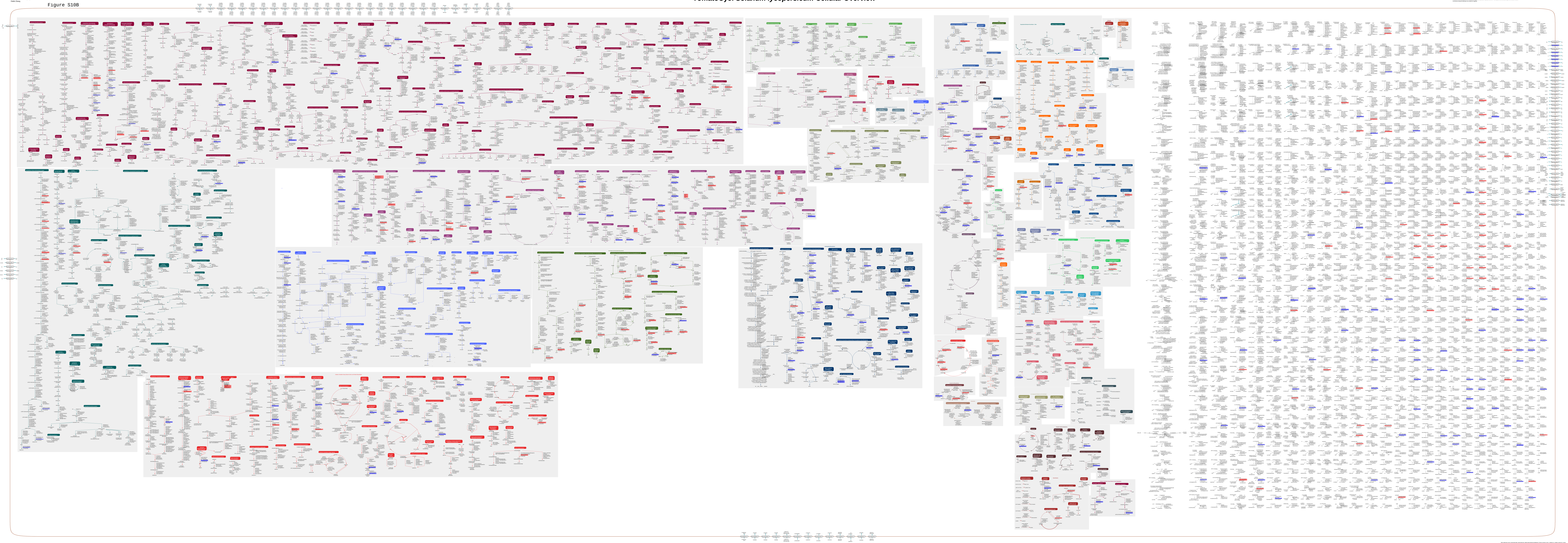

This diagram was automatically generated by SRI International Pathway Tools version 24.0, authors S. Paley and P.D. Karp.

Authors: Bo Xue, Angela Xu, Pascal Schlapfer, Charles Hawkins, Seung Yon Rhee, Emily Fryer, Peifen Zhang

Figure S10C

### TomatoCyc: *Solanum lycopersicum* Cellular Overview

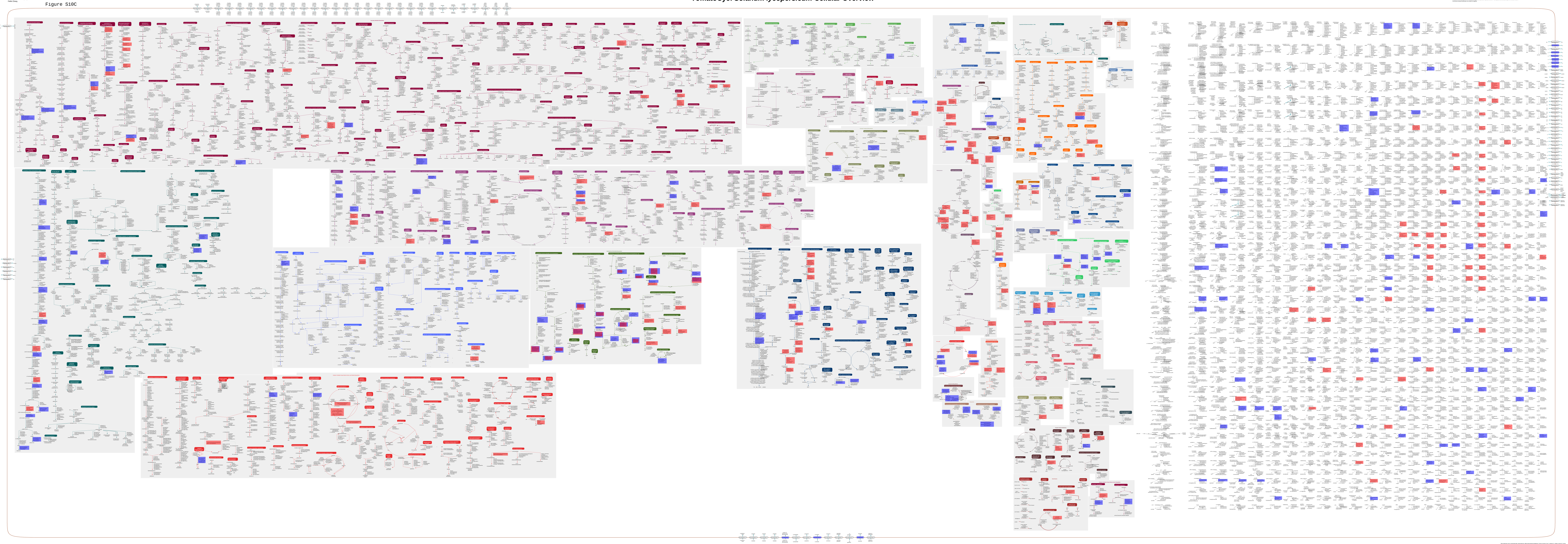

Authors: Bo Xue, Angela Xu, Pascal Schlapfer, Charles Hawkins, Seung Yon Rhee, Emily Fryer, Peifen Zhang

Figure S10D

### TomatoCyc: *Solanum lycopersicum* Cellular Overview

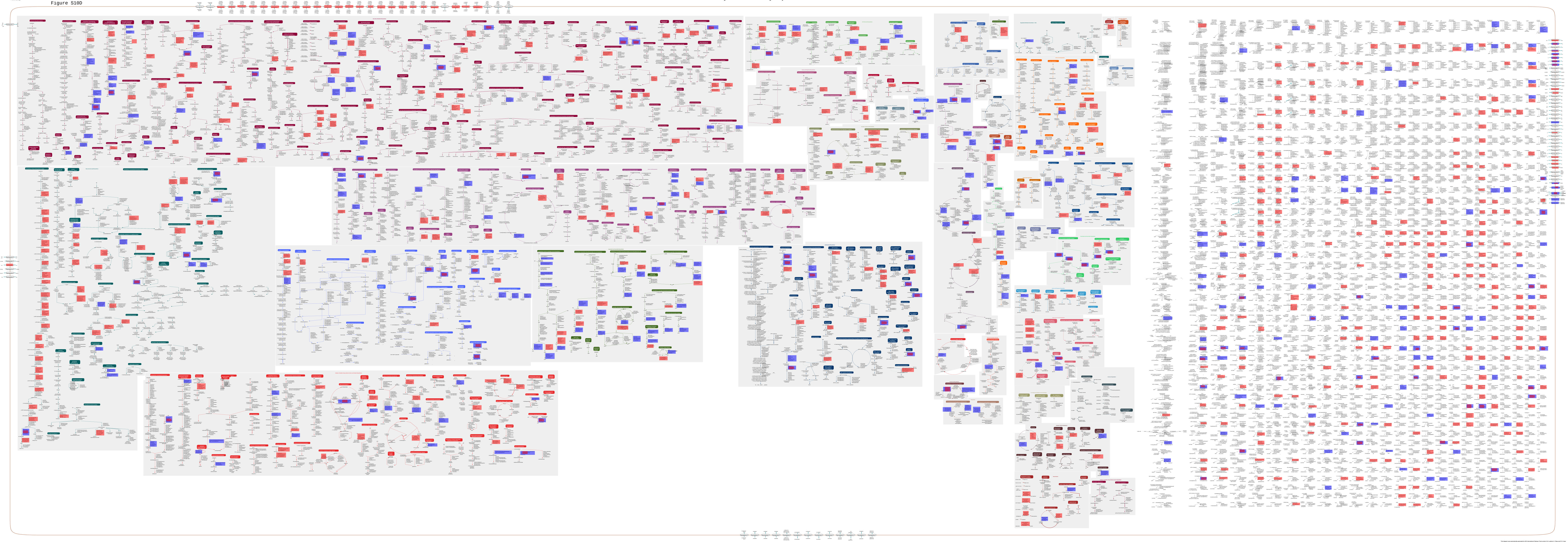
